# Understanding Thioamitide Biosynthesis Using Pathway Engineering and Untargeted Metabolomics

**DOI:** 10.1101/2020.12.17.423248

**Authors:** Tom H. Eyles, Natalia M. Vior, Rodney Lacret, Andrew W. Truman

**Affiliations:** Department of Molecular Microbiology, John Innes Centre, Norwich Research Part, Norwich, NR4 7UH, UK

## Abstract

Thiostreptamide S4 is a thioamitide, a family of promising antitumour ribosomally synthesised and post-translationally modified peptides (RiPPs). The thioamitides are one of the most structurally complex RiPP families, yet very few thioamitide biosynthetic steps have been elucidated, even though the gene clusters of multiple thioamitides have been identified. We hypothesised that engineering the thiostreptamide S4 gene cluster in a heterologous host could provide insights into its biosynthesis when coupled with untargeted metabolomics and targeted mutations of the precursor peptide. Modified gene clusters were constructed, and in-depth metabolomics enabled a detailed understanding of the biosynthetic pathway, including the identification of an effector-like protein critical for amino acid dehydration. We use this biosynthetic understanding to bioinformatically identify new widespread families of RiPP biosynthetic gene clusters, paving the way for future RiPP discovery and engineering.

## INTRODUCTION

Thioviridamide is an apoptosis-inducing compound that was isolated from *Streptomyces olivoviridis* during a screen for cytotoxic compounds^1^ and represents the founding member of the thioamitides, a structurally complex family of ribosomally synthesised and post-translationally modified peptides^2^ (RiPPs). RiPPs derive from a ribosomally synthesised precursor peptide that is modified by a series of tailoring enzymes encoded in a biosynthetic gene cluster (BGC). The discovery of the thioviridamide BGC initiated the genomics-led discovery of other thioamitides, including thioholgamide^3^ (also known as neothioviridamide^4^), thioalbamide, thiostreptamide S87, and thiostreptamide S4^5^ (**1**, Figure 1A). It was recently determined that thioamitides inhibit mitochodrial ATP synthase^6^, which induces mitochondrial dysfunction and triggers apoptosis^7^.

**Figure 1.**
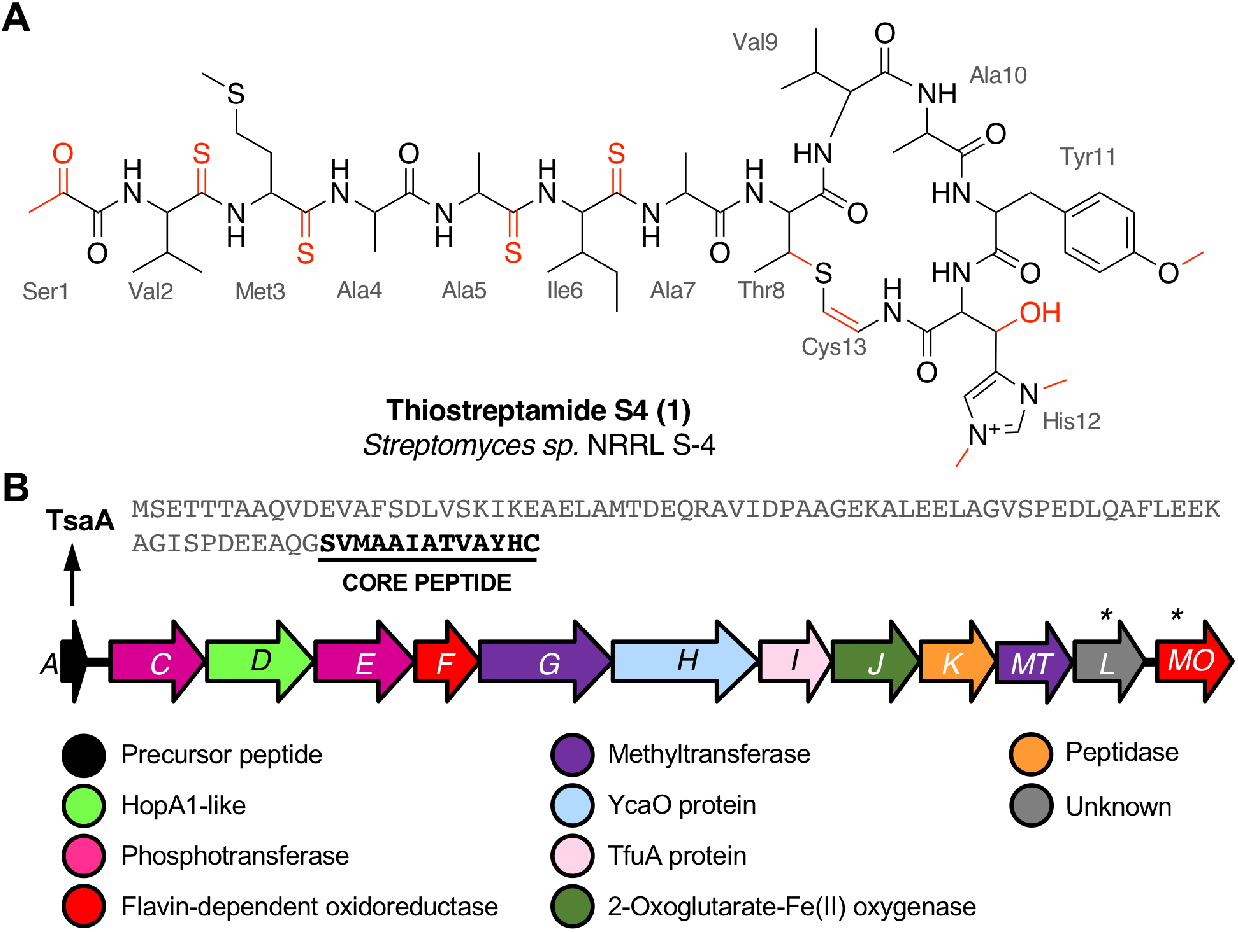
A. Thiostreptamide S4 (**1**) structure with posttranslational modifications highlighted in red and core peptide numbering. B. Thiostreptamide S4 (*tsa*) BGC and precursor peptide sequence. * indicates genes tested in this study that are unlikely to be involved in biosynthesis.

**1** features multiple post-translational modifications that are common to most thioamitides but are otherwise rare in nature, including four thioamide bonds, a 2-aminovinyl-3-methyl-cysteine (AviMeCys) macrocycle^8^, histidine bis-N-methylation, histidine β-hydroxylation, and an N-terminal pyruvyl group. **1** also features tyrosine O-methylation, which is not found in other thioamitides^5,9^. These features are interesting due to their structural and biosynthetic rarity, along with the possible influence they have on bioactivity. For example, histidine bis-N-methylation is a modification not found in other RiPPs, as is the installation of multiple thioamide bonds. However, there was limited data on thioamitide biosynthesis^10,11^. Therefore, we hypothesised that understanding thiostreptamide S4 biosynthesis would reveal new biosynthetic machinery involved in RiPP maturation. This could inform future pathway engineering and genome mining for new RiPPs with related biosynthetic machinery. Notably, thioamitide biosynthesis is predicted to require lanthipeptide-like Ser/Thr dehydrations, but homologues of the Lan proteins that usually catalyse this step are not encoded in thioamitide BGCs^2^.

Gene deletions are commonly used to understand natural product biosynthesis, as they can lead to the production of intermediates and therefore reveal the role of a gene, especially as there can be substantial challenges in the *in vitro* reconstitution of complex multi-step pathways from Actinobacteria. However, there are significant difficulties in using gene deletions to understand RiPP biosynthesis^12^. If the deleted biosynthetic gene produces a protein that acts early in a biosynthetic pathway, then the resulting precursor peptide intermediate is often unstructured and minimally modified. These peptides can be readily digested by endogenous proteases and acetylated endogenously^12^. Therefore, the identification of intermediates and shunt metabolites can be very challenging, especially if these issues are combined with low productivity in complex media (Figure S1).

Here, we use a combination of heterologous expression, gene deletions, untargeted metabolomics, and yeast-mediated core peptide engineering to understand the biosynthesis of thiostreptamide S4. This provides a genetic basis for almost every post-translational modification in thioamitide biosynthesis. In addition, the identification of genes associated with Ser/Thr dehydration enables the discovery of new widespread RiPP BGC families across multiple bacterial taxa.

## RESULTS AND DISCUSSION

### Identification of essential biosynthetic genes

We had previously cloned the thiostreptamide S4 (*tsa*) BGC (Figure 1B, Table S1) from *Streptomyces* sp. NRRL S-4 using transformation associated recombination (TAR) cloning^13,14^ in yeast to produce plasmid pCAPtsa^5^. Heterologous expression of pCAPtsa in *Streptomyces coelicolor* M1146^15^ produced complete thiostreptamide S4, providing evidence that every gene required to produce thiostreptamide S4 was present. However, the region captured via TAR cloning covered a larger region than the predicted BGC (*tsaA*-*tsaMO*; Figure S2A). Three genes, *tsa-3*, *tsa-2*, and *tsa-1*, were captured upstream from the predicted gene cluster and were predicted to encode a DNA polymerase III δ subunit, a hydrolase, and a threonine tRNA respectively. Three genes were captured downstream from the predicted BGC, *tsa+1*, *tsa+2*, and *tsa+3*, predicted to encode a transporter, sulphurtransferase, and serine/threonine protein kinase respectively.

All genes in the predicted *tsa* BGC were independently deleted in pCAPtsa using PCR targeting, replacing most of the target gene with an in-frame 81 bp scar sequence while retaining the original start and stop codons^16^. The up- and downstream regions described above were also deleted. These mutated plasmids were then expressed in *S. coelicolor* M1146. This revealed that *tsaA-tsaJ* and *tsaMT* were required for the biosynthesis of **1**, whereas the molecule was still produced in Δ*tsaK*, Δ*tsaL* and Δ*tsaMO* (Figure S2B; for simplicity, each *S. coelicolor* M1146 strain harbouring a mutated version of pCAPtsa will herein be referred to by the mutation only). Production of **1** following deletion of *tsaK* was surprising given that this gene is conserved amongst thioamitide BGCs^5^ and encodes a cysteine protease that we predicted was involved in leader peptide removal. It is possible that native peptidases from the heterologous host, *S. coelicolor* M1146, can complement this deletion as there was a small drop in the production of **1** (Figure S3). TsaK may only be necessary when **1** is produced in the native host. Similarly, *tsaL*-like genes are conserved amongst almost all thioamitide BGCs, although there is no clear catalytic domain in TsaL (Table S1). This analysis also demonstrated that *tsaMO* is not required for the biosynthesis of **1**.

The deletion of the block of upstream genes, *tsa-3*, *tsa-2*, and *tsa-1*, caused a significant drop in production (Figure S3), whereas production was unaffected by deletion of *tsa+1*, *tsa+2*, and *tsa+3*. With the exception of Δ*tsaA*, each deletion that abolished production was successfully complemented (Figure S2B), which ensured there were no unwanted polar effects of gene deletions. Genetic complementation experiments were carried out by expressing the gene from the strong constitutive promoter PermE*^17^ in pIJ10257^18^, which integrates into a ϕBT1 site in the *S. coelicolor* M1146 genome. This enabled us to determine the correct start codon of each gene (Table S3, Figure S4), which revealed that there are two series of genes with overlapping start and stop codons within the BGC, *tsaC*-*G* and *tsaH*-*MT*, with an untranslated 28 bp region between *tsaG* and *tsaH*.

### Identifying the genetic basis for macrocycle modifications

The *C*-terminal macrocycle of **1** features a bis-N-methylated and β-hydroxylated histidine, as well as an O-methylated tyrosine. There are homologues of TsaG (methyltransferase, pfam06325) and TsaJ (2-oxoglutarate-Fe(II) oxygenase, pfam05721) encoded across almost all thioamitide BGCs, so these were predicted to install the conserved histidine N-methylations and β-hydroxylation, respectively. Amongst characterised thioamitide BGCs, TsaMT (methyltransferase, pfam13649) is only encoded in the thiostreptamide S4 BGC, so was predicted to install the unique O-methylation. We therefore searched liquid chromatography-mass spectrometry (LC-MS) spectra of wild type (WT), Δ*tsaG*, Δ*tsaJ* and Δ*tsaMT* cultures for masses matching the loss of one to three methyl groups, and/or one hydroxyl group. In total, five of these masses were detected: **1** (*m/z* 1377.55), **2** (*m/z* 1363.53, −1 methyl), **3** (*m/z* 1361.55, −1 hydroxyl), **4** (*m/z* 1347.54, −1 methyl and −1 hydroxyl), and **5** (*m/z* 1319.50, −3 methyl and −1 hydroxyl). Tandem MS (MS/MS) fragmentation data confirmed that the mass differences were on the macrocycle and accurate masses are consistent with these proposed structures (Figure S5). **1**, **3**, and **4** are seen in the WT, **5** is seen in Δ*tsaG*, **3** and **4** are seen in Δ*tsaJ*, and **2** and **4** are seen in Δ*tsaMT* (Figure 2).

**Figure 2.**
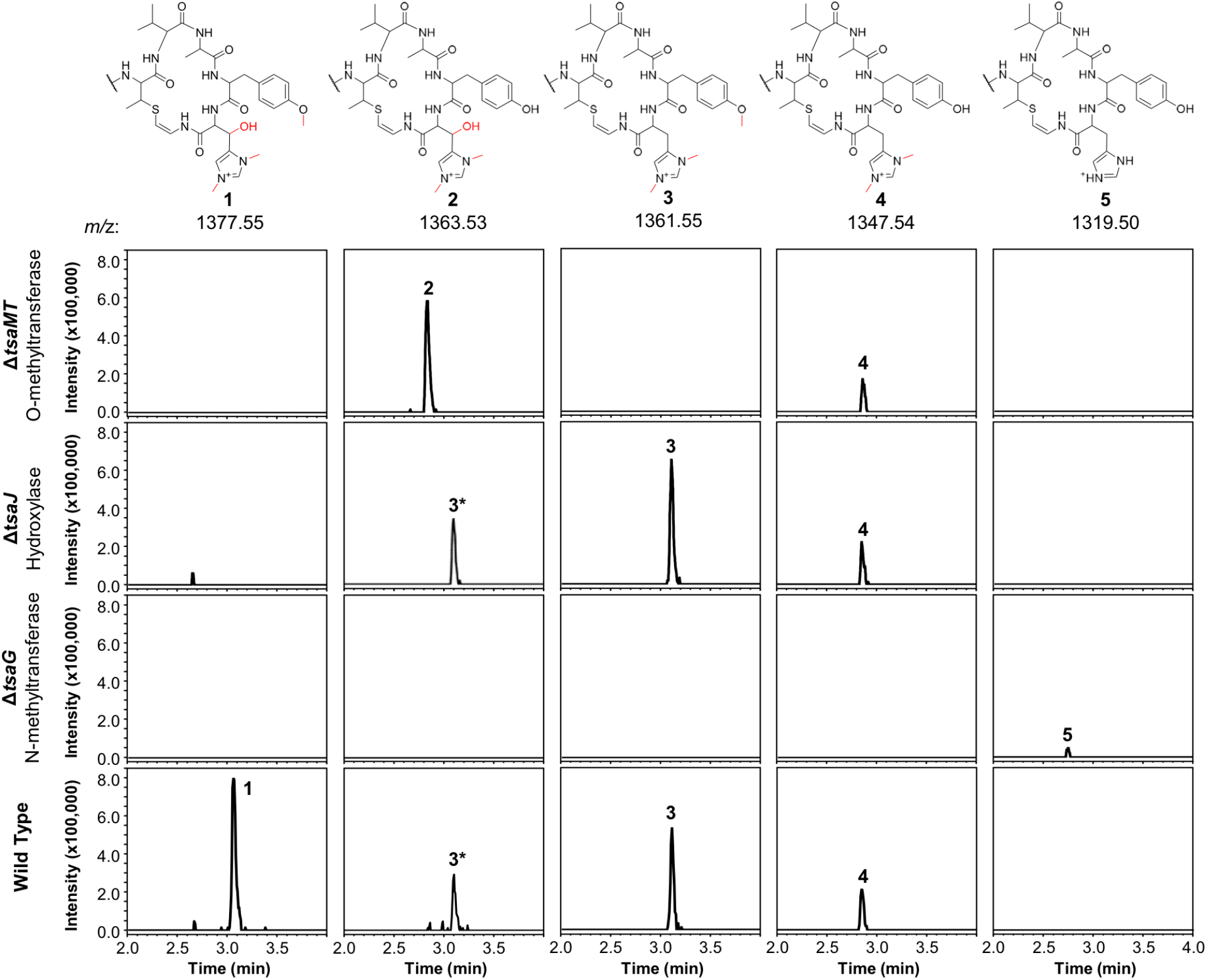
Extracted ion chromatograms (EICs) normalised for intensity showing the varied methylation and hydroxylation patterns of thiostreptamide-like molecules produced in *S. coelicolor* M1146 expressing the WT, *ΔtsaG*, *ΔtsaJ* and *ΔtsaMT* BGCs. Structures were inferred using detailed MS/MS analysis (Figure S5), which are consistent with **2**-**5** featuring full modifications on the N-terminal linear peptide portion (as in **1**). The structure of the macrocycle from each metabolite is shown above the relevant traces; in each case the rest of the molecule is identical to **1**. The **3\*** label indicates the +2 isotope peak of **3**.

Deletion of *tsaMT*, which encodes a class I SAM-dependant methyltransferase, resulted in the loss of **1** and **3** (Figure 2). Instead, **2** was produced, which lacks the tyrosine methylation but is otherwise identical to **1**, therefore confirming that TsaMT is the protein responsible for this modification. A retro-aldol MS/MS fragmentation that provides a loss of *m/z* 125.07 is consistent with histidine hydroxylation and bis-N-methylation in **2** (Figure S5). This shows that the tyrosine methylation is not required for the histidine hydroxylase and methyltransferase to function. Δ*tsaJ* produces **3** and **4** (Figure 2), which both lack the histidine hydroxylation. This indicates that TsaJ, a non-heme Fe(II) and α-ketoglutarate dependent dioxygenase, is responsible for histidine hydroxylation. The production of **3** shows that histidine hydroxylation is not a prerequisite for tyrosine methylation or histidine bis-N-methylation.

Deletion of *tsaG*, which encodes a SAM-dependant methyltransferase, abolished production of **1** and instead led to production of **5**, a version of **1** that lacks all modifications to the macrocycle but is otherwise fully mature (Figure 2). This means that histidine bis-N-methylation is a prerequisite for TsaJ-catalysed histidine hydroxylation and TsaMT-catalysed tyrosine methylation. This indicates that TsaJ and TsaMT are unable to recognise a TsaA-derived substrate without the histidine methylations, which provide a permanent positive charge. The thioamitides are the only RiPPs that features a bis-N-methylated histidine. Given that the macrocycle is correctly formed in each mutant, these results are consistent with a biosynthetic model where these modifications to the macrocycle are among the final steps in thiostreptamide S4 biosynthesis. TsaG-catalysed histidine methylation occurs first, which is then followed by TsaJ-catalysed histidine hydroxylation and TsaMT-catalysed tyrosine methylation in an undefined order. The role of TsaJ is consistent with a parallel study on the homologue in thioholgamide biosynthesis, ThoJ^11^.

To see if other thiostreptamide-like metabolites were produced by these mutants, the characteristic fragmentation pattern of these metabolites was used to search the LC-MS/MS data from the WT, Δ*tsaG*, Δ*tsaJ*, and Δ*tsaMT* strains. The macrocycle is one of the main fragments of **1**-**5**, and so the masses of the different macrocycle fragments seen in **1**-**5** (*m/z* 687.33, 673.31, 671.33, 657.32, and 629.29, respectively) were used to search all fragmentation events in LC-MS/MS spectra. This enabled the preliminary identification of six new metabolites, **6**-**11** (Figures S7-S8). **6**-**10** are proposed to be versions of **1**-**5** that are hydrolysed between Ala4 and Ala5 (Figure S7), while **11** is predicted to be a version of **5** where the other non-thioamide bond between Ala7 and the macrocycle is hydrolysed (Figure S8). These therefore result from hydrolysis of the only non-thioamidated peptide bonds in the tail portion of the molecule, which supports previous evidence that thioamide bonds protect molecules from proteolysis^19^.

### Identifying thiostreptamide-related metabolites using untargeted metabolomics

In contrast to the genes encoding macrocycle-modifying enzymes, it was difficult to predict likely pathway products for all other gene deletions (Δ*tsaC*, Δ*tsaD,* Δ*tsaE*, Δ*tsaF*, Δ*tsaH* and Δ*tsaI*), and the targeted analysis described above was unable to identify any macrocycle-containing molecules from these mutants. To address this challenge, MS-based untargeted metabolomics was employed to detect any pathway-associated metabolites. Mutants were compared to *S. coelicolor* M1146-pCAPtsa, Δ*tsaA*, and a medium only control. By filtering out all metabolites present in Δ*tsaA*, we were able to identify multiple metabolites across almost all mutant strains that were likely to derive from TsaA, the precursor peptide (Figure 3, Table S3).

**Figure 3.**
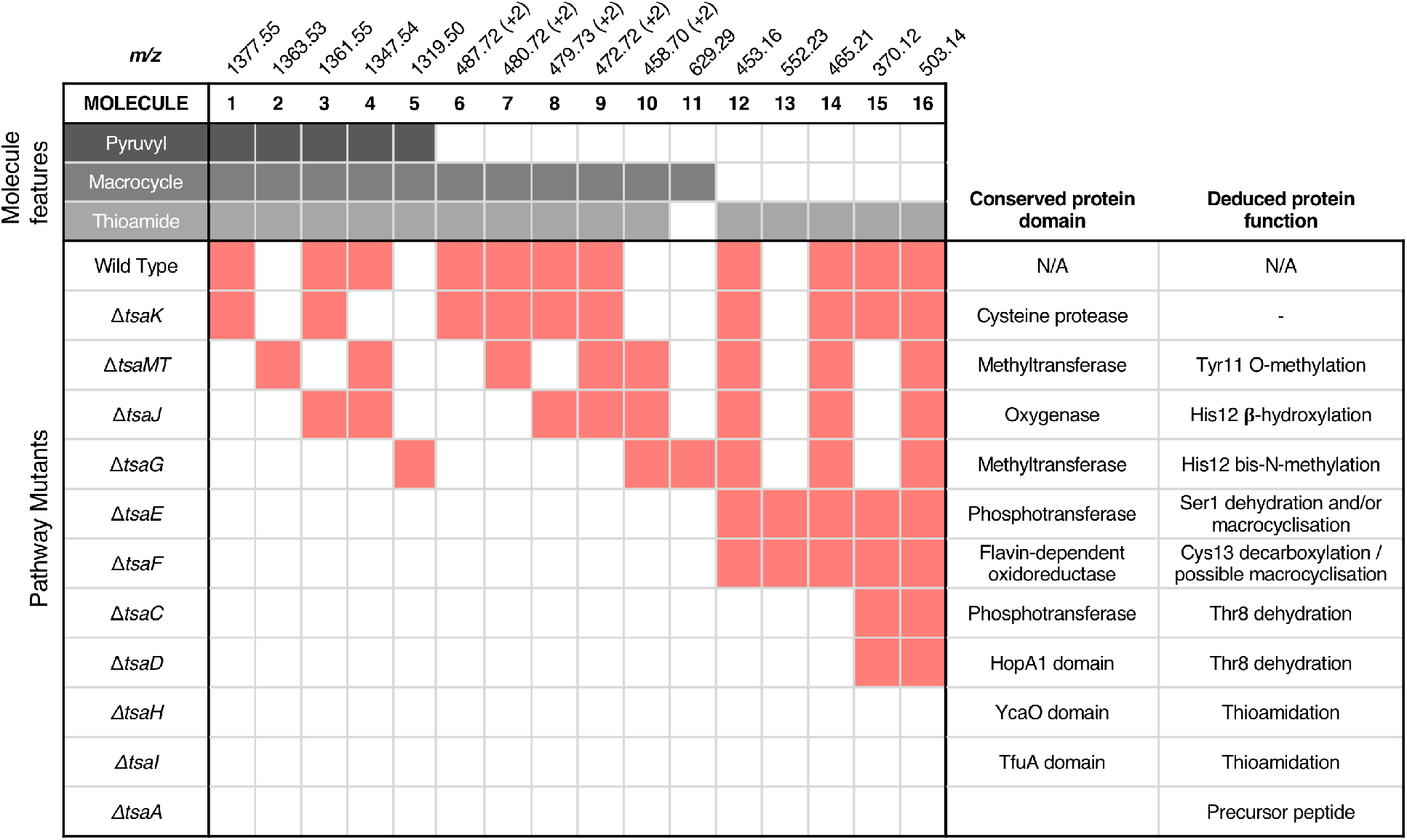
Untargeted metabolomic analysis of thiostreptamide S4 biosynthesis. Red shading indicates the production of a molecule in a given mutant. Proposed structures of molecules **2**-**16** are summarised in Figure S17, where MS/MS and accurate mass data is shown in Figures S5-S10. “+2” indicates that the doubly charged *m/z* is shown.

There were numerous difficulties in interpreting these data. Identification by MS/MS initially proved difficult due to limited fragmentation, and fragments that were observed could not be accounted for by the simple loss of proteinogenic amino acids. Notably, molecules containing thioamide bonds can undergo fragmentation to lose SH2; corresponding to a mass loss of 33.9877 Da that does not break the backbone of the molecule^20^. This signature loss can be seen very clearly in the fragmentation of **1** (Figure S6) and can be used as a tool to identify metabolites that contain thioamides. This indicated that previously unidentified metabolites (*m/z* 552.23, 503.15, 465.22, 453.16, 392.11, 370.12, 348.14, 330.13 and 259.09) have thioamide bonds in their structure due to this signature fragmentation and are not produced by the Δ*tsaA* mutant (Table S4). These molecules are hypothesised to be short shunt metabolites that are protected from proteolytic degradation by thioamidation^19^.

### Chemical characterisation of a thioamidated shunt metabolite

To support the preliminary interpretation of thioamidated shunt metabolites, the signature SH2 loss was used to target metabolites for detailed chemical characterisation. **12** (*m/z* 453.16) was targeted for purification due to its high production levels and because numerous metabolites featured comparable MS/MS fragmentation (Figure S9). **12** was purified from *S. coelicolor* M1146 expressing the Δ*tsaE* BGC, yielding 0.7 mg of pure compound. This was characterised by NMR (^1^H, COSY, DEPTQ, HSQCed and HMBC, Figures S11-S16, Table S5). ^13^C shifts of 206.0 and 203.7 ppm were indicative of two thioamides, while a ^13^C shift of 136.4 ppm was consistent with an olefinic methine. Two-dimensional experiments established the amino acid connectivity to show that **12** is a modified tetrapeptide, *N*-acetyl-Ala^S^Ile^S^AlaDhb (Dhb = 2,3-dehydrobutyrine; superscript S = thioamidated amino acid) (Figure 4), whose molecular formula of C_18_H_30_N_4_O_4_S_2_ was consistent with a high-resolution MS peak of *m/z* 453.1584 ([M+Na]+, calc. *m/z* 453.1601). **12** is therefore a portion of TsaA (Ala5 to Thr8, Figure 1A) that has undergone expected thioamidation of Ala5-Ile6 and dehydration of Thr8 to Dhb8, but has not been macrocyclised and instead has been hydrolysed at unmodifed peptide bonds. These modifications help explain the difficult to interpret MS/MS data, as does an atypical MS/MS fragmentation pattern that occurs for sodiated peptides^21,22^ (Figure S9B).

**Figure 4.**
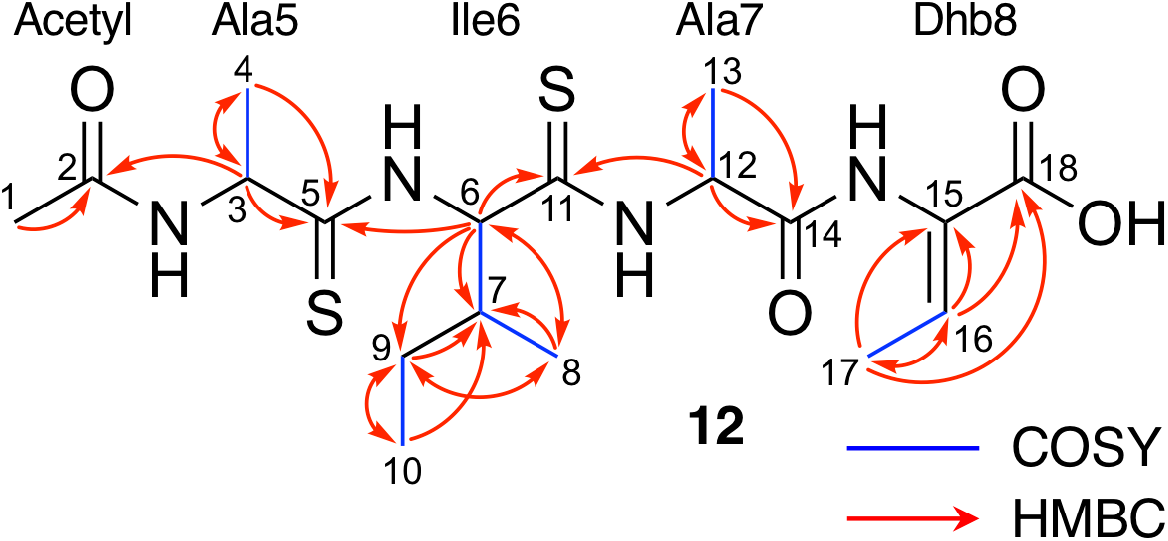
NMR characterisation of **12** in CD3OD. See Figures S11-S16 and Table S5 for NMR assignments.

Following the characterisation of **12**, the structures of **13** (*m/z* 552.23), **14** (*m/z* 465.21), and **15** (*m/z* 370.12) could also be proposed to be related acetylated and thioamidated short peptides, based on similar MS/MS fragmentation patterns, accurate mass data and predicted thioamidations (Figure S9A). This similarly enabled us to propose the structure of **16** (*m/z* 503.14), a metabolite produced by the WT and all Δ*tsaC-F* mutants (Figure 3 and Table S4). **16** is proposed to be N-acetyl-SerVal^S^Met^S^Ala, which we hypothesise derives from the precursor peptide (Ser1 to Ala4, Figure 1A) that has undergone expected thioamidation of Val2-Met3 (Figure S9). Further support for this structure was provided by precursor peptide modifications (S1T and M3I), which led to expected mass shifts to this metabolite (Figure S10; see later section for a description of site-directed mutagenesis).

### Thioamidation requires a YcaO protein and a TfuA protein

Prior studies have demonstrated that YcaO and TfuA domain proteins are required for thioamidation in archaea^23,24^ and bacteria^20,25^, although the precise role of the TfuA domain protein is not known. We therefore predicted that YcaO protein TsaH and TfuA protein TsaI would iteratively introduce the four thioamides in **1**. Deletion of either *tsaH* or *tsaI* led to the abolition of every detectable metabolite associated with the BGC (Figure 3). This is in contrast to every other mutant, which were all able to make thioamidated peptides (Figure 3). This supports a biosynthetic model where TsaH and TsaI cooperate to catalyse thioamidation. The absence of detectable metabolites is consistent with TsaH/TsaI functioning as essential steps at a very early stage in the pathway, as other modifications that could protect TsaA from proteolysis, such as macrocyclisation, were not detected in these mutants.

### Identification of a new amino acid dehydratase

Along with thioamidation, a characteristic feature of thioamitides is a *C*-terminal Avi(Me)Cys macrocycle. In **1**, this is predicted to be generated by the Michael-type addition of oxidatively decarboxylated Cys13 with Dhb8, which is formed by dehydration of Thr8. Another conserved thioamitide feature is an N-terminal pyruvyl or lactyl moiety that we previously predicted to be derived from dehydrated Ser1^5^. In lanthipeptide biosynthesis, the dehydration of Thr to Dhb is catalysed by Lan proteins^26^. However, no Lan proteins are encoded in thioamitide BGCs. We were unable to detect full-length core peptides featuring Thr8, but we hypothesised that the presence/absence of metabolites **12**-**16** would help identify genes involved in dehydration and potentially cyclisation. **12**-**16** were therefore mapped to the metabolomes of mutants of genes that had not yet been functionally annotated (Δ*tsaC*, Δ*tsaD,* Δ*tsaE*, Δ*tsaF*; Figures 3 and 5).

**Figure 5.**
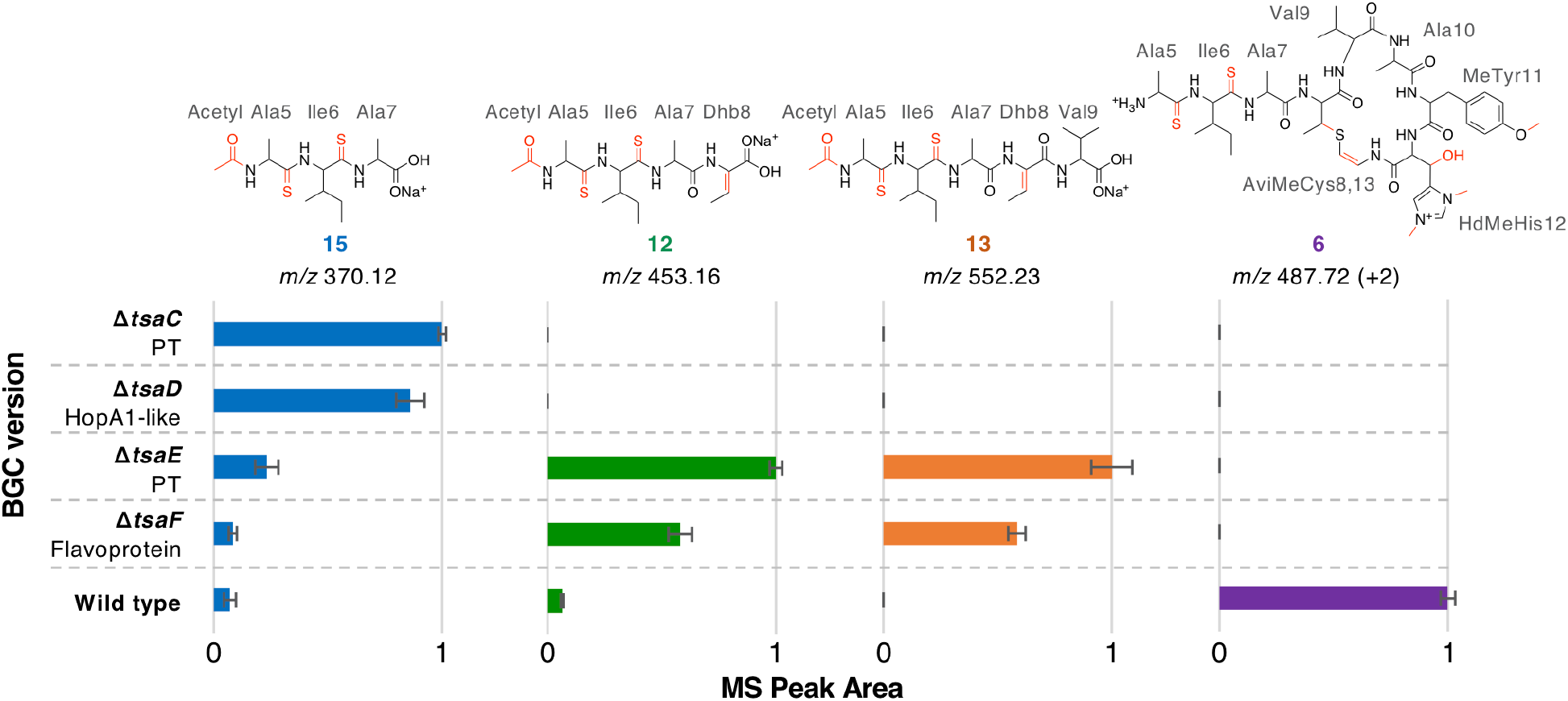
MS peak areas of selected metabolites produced by mutants (*ΔtsaC-F*) and WT thiostreptamide S4 BGC. Each bar chart is normalised to the highest mass spectral peak area for that metabolite. The error bars represent the standard error of three biological repeats. PT = phosphotransferase; “+2” indicates that the doubly charged *m/z* is shown.

The structure of **12** (Figure 4), and the predicted structures of **13**-**16** (Figure S17), provide key information towards the proteins involved in dehydration (Figure 5). **12-14** are shunt metabolites of an intermediate that lacks the macrocycle but contains the Dhb8 residue that is required for macrocycle formation. In contrast, **15** can derive from an intermediate that contains an unmodified Thr8; the lack of modification making it susceptible to proteolysis. In all strains containing deletions of any of *tsaC-F,* all detected metabolites lack the macrocycle (**12**-**16**), which implies that they are involved in steps previous to its formation. Of these mutants, Δ*tsaC* and Δ*tsaD* produce thioamidated **15** and **16** (Figure 3) but do not produce shunt metabolites containing Dhb8. We hypothesise that these metabolites derive from a modified TsaA that is not yet dehydrated at Thr8 and is therefore more susceptible to proteolysis at that position. This would indicate that TsaC and TsaD cooperate to catalyse dehydration of Thr8 to Dhb8.

TsaC contains an aminoglycoside phosphotransferase (APH)-like domain (pfam01636). APHs are structurally similar to eukaryotic protein kinases^27^ and it has been shown that some APH enzymes can also phosphorylate serine residues^28^. We therefore propose that TsaC is responsible for phosphorylation of Thr8, allowing for a subsequent elimination reaction to dehydrate Thr8 (Figure 4). The role of TsaD in threonine dehydration is currently unclear, although TsaD contains a HopA1 effector family domain (pfam17914). HopA1 itself is a type III effector from *Pseudomonas syringae*^29,30^. HopA1 aids plant infection by this pathogen, although the mechanistic basis for this activity is unknown. Other effectors, such as the OspF family^31^, do function as lyases to inactivate protein kinases in the host cell. The N-terminal lyase domain of the LanL family of lanthionine synthetases has sequence homology with OspF proteins^32^. TsaD may therefore act as a lyase to catalyse the elimination of a TsaC-installed phosphate group to dehydrate Thr8. Alternatively, TsaD may have a non-catalytic role that is essential for TsaCD-catalysed dehydration, such as precursor peptide binding. Very recently, two RiPPs containing lanthionine cross-links and AviMeCys macrocycles were reported (cacaoidin^33^ and lexapeptide^34^), whose BGCs encode homologues of TsaC and TsaD. Intriguingly, our results are in contrast to a very recent report on the biosynthesis of the thiosparsoamide, which indicated that lanthipeptide synthetases encoded outside of the BGC catalyse thioamitide dehydration^35^.

### Macrocyclisation is dependent on TsaE and TsaF

In contrast to Δ*tsaC* and Δ*tsaD*, **12**, **13** and **14** are produced by both Δ*tsaE* and Δ*tsaF* (Figures 3 and 5). Therefore, neither gene is required for the dehydration of Thr8 to Dhb8. However, no macrocyclised molecules (**1**-**11**) are produced by either Δ*tsaE* or Δ*tsaF*. TsaF is predicted to be a flavoprotein (pfam02441) that has similarity to flavin-dependent cysteine decarboxylases that catalyse the formation of AviCys^36^, AviMeCys, and avionin macrocycles^37^ in RiPP biosynthesis. In Avi(Me)Cys-containing natural products this oxidative decarboxylation forms the thioenolate moiety required for Avi(Me)Cys formation^38^. The lack of macrocyclised molecules when *tsaF* is deleted supports the role of TsaF as a cysteine decarboxylase that decarboxylates Cys13. This is consistent with a recent co-expression study using the thioviridamide orthologue, TvaF, which was shown to catalyse oxidative decarboxylation of the thioviridamide precursor peptide^10^. In lanthipeptide biosynthesis, a cyclase is required to catalyse cyclisation, where one of the roles of the cyclase is to stabilise the thiolate involved in macrocycle formation^26^. However, the formation of the AviMeCys thioether may be spontaneous, as the enethiolate that results from cysteine decarboxylation has a significantly lower p*K*_a_ than the thiol side chain of cysteines^39^. This would explain the lack of a cyclase homologue encoded in thioamitide BGCs or in other Avi(Me)Cys containing RiPP BGCs, such as the linaridins^40^.

TsaE possesses weak homology to the APH-like phosphotransferase domain (pfam01636) that is also found in TsaC. However, Δ*tsaE* has a very similar metabolite profile to Δ*tsaF* (Figures 3 and 5, Table S4). It is somewhat surprising that the macrocycle cannot form in Δ*tsaE*, given that Dhb8-containing molecules are produced by this mutant and the cysteine decarboxylase, TsaF, is present. The lack of macrocycle could be explained if TsaE assists with AviMeCys cyclisation. However, it is unclear what role a phosphotransferase could play in cyclisation and there are no *tsaE* homologues encoded in BGCs for other Avi(Me)Cys RiPPs, such as the linaridins. An alternative hypothesis is that TsaE functions as the phosphotransferase involved in Ser1 dehydration to 2,3-dehydroalanine (Dha), which we predicted to be necessary for the formation of the *N*-terminal pyruvyl group of **1**.

To demonstrate that the pyruvyl group originates from a dehydrated Ser1 instead of a pyruvyl transferase^41^, a S1T mutant of TsaA was generated, producing the construct pCAPtsaS1T. *S. coelicolor* M1146-pCAPtsaS1T produced a molecule with *m/z* 1391.5598 (**17**) that was absent in the WT (Figure S18). This mass reflects an extra methyl group compared to **1** and MS/MS fragmentation is consistent with **17** containing an *N*-terminal 2-oxobutyryl moiety instead of a pyruvyl moiety (Figure S18). This therefore confirms that the natural *N*-terminal pyruvyl group originates from a dehydrated serine. Hydrolytic removal of the leader peptide generates an enamine in equilibrium with an imine that is predicted to spontaneously hydrolyse to the pyruvyl group (Figure 6E). A Thr1 residue is seen naturally in the predicted core peptides for uncharacterised thioamitides from *Micromonospora eburnea* and *Salinispora pacifica*^5^. The amino acid origin of the pyruvyl group is consistent with previous co-expression studies on epilancin 15X^42^ and polytheonamide dehydratases^43^. The serine origin of the pyruvyl group means that it is plausible that TsaC, TsaD and/or TsaE are involved in Ser1 dehydration, given that **16** is proposed to contain an unmodified Ser1 residue and is produced by each of these mutants (Figure 3, Figure S10), but further experimental work is required to confirm the serine dehydratase.

**Figure 6.**
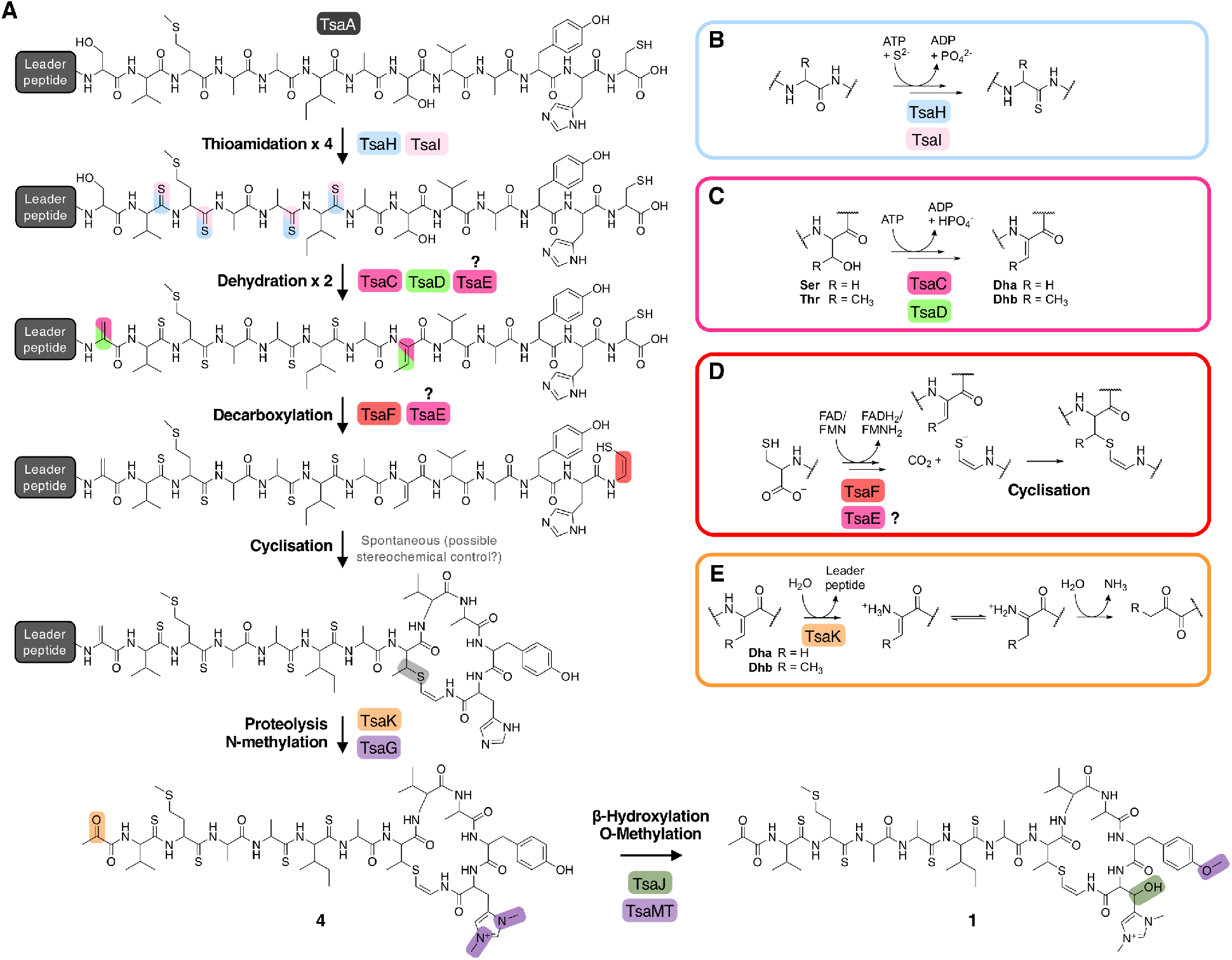
A. Proposed biosynthetic pathway to thiostreptamide S4 (**1**). B. Proposed route to thioamidation based on prior studies of archaeal thioamidation. C. Proposed route to dehydration of serine (R = H) or threonine (R = CH3) via phosphorylation and elimination. D. Proposed route to cysteine decarboxylation and cyclisation. E. Proposed production of pyruvyl or 2-oxobutyryl moiety from an N-terminal Ser1 or Thr1 (S1T mutant) following leader peptide proteolysis.

### Proposed biosynthetic pathway to 1

The metabolomic data from the gene deletions enables a plausible biosynthetic pathway to be proposed (Figure 6A). The absence of any detectable metabolites in Δ*tsaH* and Δ*tsaI*, as well as the presence of thioamidated peptides in all other mutants, indicates that the first step is the thioamidation of the TsaA core peptide by TsaH (YcaO-like) and TsaI (TfuA-like) (Figure 6B). The absence of metabolites containing Dhb8 (or macrocycles) in Δ*tsaC* and Δ*tsaD* is consistent with TsaC/TsaD-catalysed dehydration of Thr8 to Dhb8. TsaC is a phosphotransferase, which would be consistent with a phosphorylation and elimination mechanism typical of class II, III, and IV lanthipeptide dehydratases (Figure 6C). Our data suggest that HopA1-like TsaD is either a lyase responsible for phosphate elimination, or a non-catalytic partner protein. At some stage following this step, Ser1 is dehydrated, which is predicted to proceed via the same mechanism. This may involve TsaE-catalysed phosphorylation, but it also cannot be ruled out that TsaC catalyses this step. The lack of macrocyclic molecules detected in Δ*tsaE* indicates that this precedes macrocycle formation, but could alternatively indicate a cryptic role for TsaE in macrocyclisation.

We propose that the AviMeCys macrocycle is then formed, which first involves Cys13 decarboxylation to generate a reactive thioenolate. This is proposed to be catalysed by flavoprotein TsaF, given the absence of cyclised molecules produced in Δ*tsaF* and the prior characterisation of the TsaF orthologue from the thioviridamide pathway. Macrocyclisation itself may be non-enzymatic (Figure 6D), given the lack of an obvious cyclase protein, although the apparent absence of multiple diastereoisomers of **1** suggests stereochemical control during AviMeCys formation.

The next step is bis-N-methylation of His12. Deletion experiments show that TsaG is responsible for this step. Histidine bis-methylation is not present in any other natural product family and provides a positive charge that may be important for biological activity^44^. TsaG has predicted structural homology to protein arginine N-methyltransferases (Table S2). Gene deletion experiments show that this methylation acts as a gatekeeper for subsequent modifications: His12 β-hydroxylation and Tyr11 O-methylation, installed by TsaJ and TsaMT respectively. Our data indicate that these proteins preferentially act on substrates containing a bis-methylated histidine. Whilst mature thiostreptamide S4-like molecules detected in this study rarely lack the histidine N-methylations, we could readily detect significant quantities of mature thiostreptamide S4-like molecules lacking the histidine β-hydroxylation and tyrosine O-methylation (Figure 2). Therefore, these modifications are not a prerequisite for leader peptide cleavage and associated pyruvyl formation, and may happen following leader peptide removal.

There is no clear data that allows the assignment of an enzyme for leader peptide cleavage. This was unexpected, as bioinformatic analysis shows that that TsaK is a C1A family cysteine protease and homologues are encoded in other thioamitide BGCs. This protease family is rare in bacteria, although a C1A family protease catalyses removal of the leader peptide in polytheonamide biosynthesis^45^. It is possible that the small change in production observed when *tsaK* is deleted (Figure S3) is because endogenous proteases catalyse hydrolysis of the leader peptide, as in the biosynthesis of many class III lanthipeptides^46^. Following proteolysis, the pyruvyl group is likely to be formed spontaneously from Dha1 (Figure 6E). This is supported by the production of **17** (featuring an N-terminal 2-oxobutyryl group) when the precursor peptide contains a S1T mutation (Figure S18).

### Yeast assembly enables site-directed mutagenesis of precursor peptide TsaA

Precursor peptide mutagenesis can be important to probe specific biosynthetic steps or hypotheses, to test the substrate specificity of enzymes, and to generate RiPP libraries. However, it was not possible to complement the precursor peptide Δ*tsaA* mutant with an intact copy of the *tsaA* gene present in the integrative plasmid pIJ10257, which may be due to insufficient levels of expression from the non-native PermE* promoter. Therefore, we employed a yeast-mediated recombination strategy^47^ to introduce clean modifications to the precursor peptide in pCAPtsa. Here, the vector was digested using naturally occurring unique restriction enzyme sites near *tsaA* (AflII and SrfI) and then reassembled in a single step in yeast using PCR fragments and a single-stranded synthetic oligonucleotide that contains a mutated core peptide sequence (Figure S19A).

This strategy was used to generate the S1T mutant that was discussed earlier. To test the tolerance of the biosynthetic enzymes to modifications to the macrocycle amino acids, we made four further mutants of the TsaA core peptide: T8S, Y11V, H12A and H12W (Figure S19B). T8S was constructed to assess whether dehydration and macrocyclisation takes place when Thr8 is swapped with a serine residue, which is found in this position in some related precursor peptides^5^. This led to the production of **18**, which has a mass (calc. *m/z* 1363.5311, obs. m/z 1363.5261) and MS/MS fragmentation that is consistent with a fully modified derivative of **1** featuring the expected AviCys moiety (Figure S20). In contrast, the other modifications were not tolerated, as no macrocyclised molecules were detected with the Y11V, H12A and H12W mutants. However, an increase in the production of **12** (Figure S21) in each mutant indicated that early stage thioamidation and Thr8 dehydration took place, but either TsaE or TsaF would not function.

A common metabolite detected throughout growth and extraction of thiostreptamide S4 (**1**) is the methionine sulphoxide derivative (**19**; Figure S22). Met3 is particularly susceptible to oxidation, which is problematic if this molecule was used in a clinical setting, as the methionine sulphoxide version of a similar molecule, thioholgamide, is around ten times less active than un-oxidised thioholgamide^3^. To engineer thiostreptamide S4 into a more stable molecule, a version was made with Met3 swapped for an isoleucine (M3I), which is naturally found at this position in the thioalbamide precursor peptide^5^. This modification was tolerated and led to the production of **20** (*m/z* 1359.59, Figure S23). As with a site-directed mutagenesis study on thioviridamide^48^, these data indicate that precursor peptide mutagenesis represents a viable route to novel thioamitides, although the complexity of these pathways means that there are mutants that are not tolerated by all tailoring enzymes (Figure S19B).

### Understanding thioalbamide biosynthetic modifications

The *Amycolatopsis alba* thioalbamide BGC contains genes that encode for a predicted cytochrome P450 (TaaCYP; pfam00067) and a NAD(P)H-dependent reductase (TaaRed; pfam00106) that are absent from almost all other thioviridamide-like BGCs (Figure S24). The function of these genes could not be tested directly in *A. alba* because attempts to genetically manipulate this strain were unsuccessful. Therefore, *taaCYP* and *taaRed* were expressed in *S. coelicolor* M1146-pCAPtsa to test if their activity could be reconstituted on a similar molecule. Thioalbamide contains an N-terminal lactyl group, and we previously predicted that TaaRed catalyses the reduction of the Ser1-derived pyruvyl group to a lactyl group^5^. This would be analogous to the generation of a lactyl group in epilancin 15X biosynthesis by ElxO, a NAD(P)H-dependant reductase^49^. Co-expression of TaaRed with pCAPtsa generated a thiostreptamide S4 derivative (**21**) that is 2 Da bigger than 1. The accurate mass (*m/z* 1379.5624) and MS/MS fragmentation data for **21** is consistent with an *N*-terminal lactyl group (Figure S25). Our data provide preliminary evidence that TaaRed has broad substrate tolerance, given the significant differences between thioalbamide and **1**.

Thioalbamide has a hydroxylated Phe5 not seen in other characterised thioviridamide-like compounds, so it was hypothesised that TaaCYP is responsible for this hydroxylation. To test this, we used yeast-mediated assembly to generate two new versions of the thiostreptamide S4 BGC with mutated *tsaA* genes: one encoding a core peptide with a containing a phenylalanine at position 5 (A5F), and TsaCoreTaa, where the entire thiostreptamide S4 core peptide was replaced with the thioalbamide core peptide (Figure S19B). Unfortunately, no related metabolites could be detected when these clusters were expressed in *S. coelicolor* M1146, meaning that these modifications were not tolerated by the thiostreptamide S4 tailoring enzymes.

### Genome mining reveals that the HopA1 and phosphotransferase protein pair define a widespread RiPP family

A key step of thiostreptamide biosynthesis is dehydration, which we propose is catalysed by phosphotransferase TsaC and HopA1-like protein TsaD. We were therefore interested in determining whether this represented an overlooked but widespread RiPP modification and could therefore be used to identify new RiPP BGCs. A similarity-based search for TsaD-like proteins in GenBank identified 1,340 non-redundant HopA1 domain proteins across multiple bacterial phyla. This was filtered using a 95% identity cut-off to 742 proteins (see Supplementary Methods for full details). 41% of these are in Cyanobacteria, approximately two thirds of the rest come from Actinobacteria. The remaining representatives are distributed between Proteobacteria, Bacteroidetes, Firmicutes, Acidobacteria and some rare bacterial phyla. In almost all cases, the HopA1 protein is encoded alongside a phosphotransferase protein, supporting the theory that these are partner proteins that cooperate to catalyse one reaction (Figure 7).

**Figure 7.**
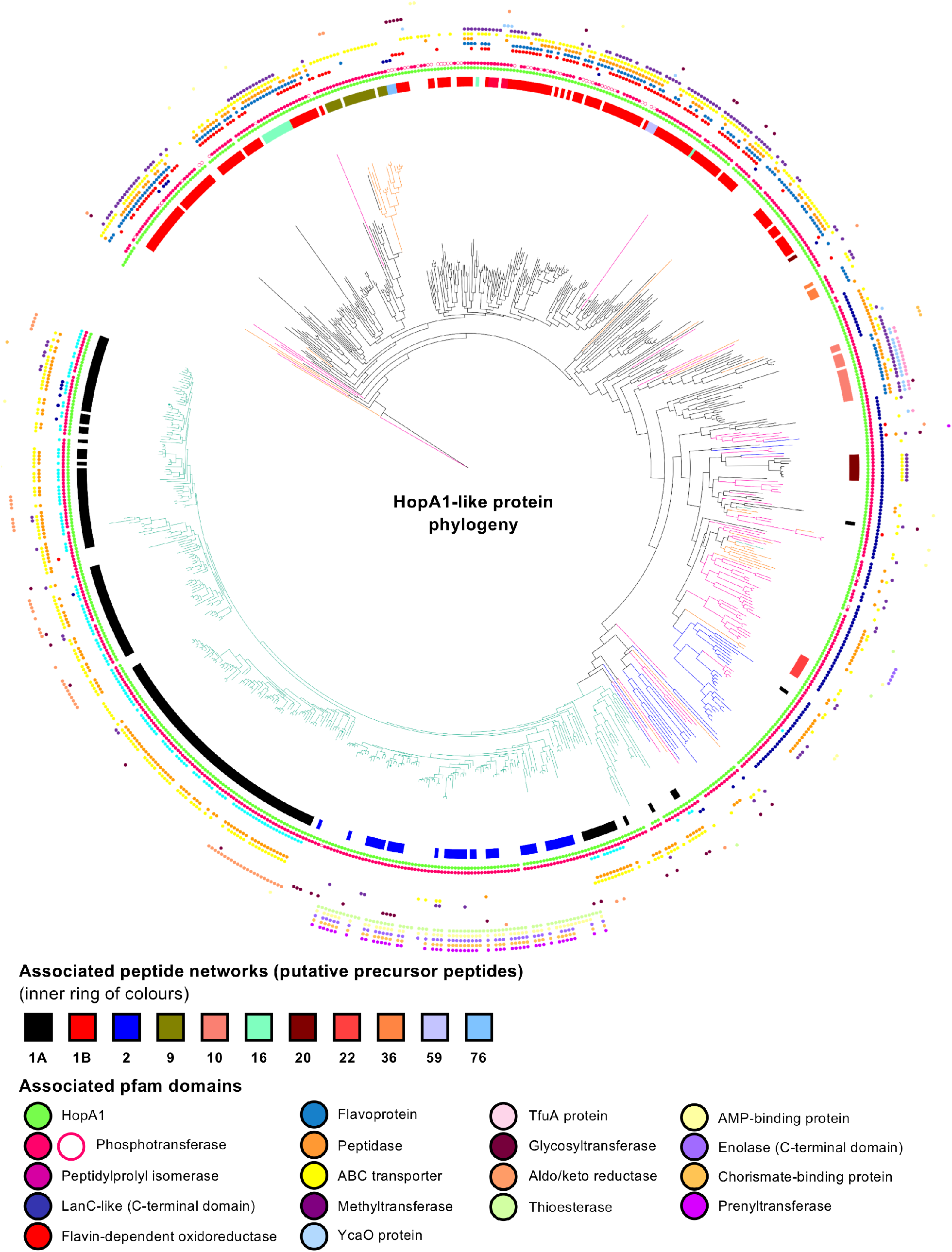
Maximum likelihood phylogenetic tree of HopA1 domain containing proteins analysed in this work. The tree branches are colour coded according to the taxonomic origin of the protein: Cyanobacteria (turquoise), Actinobacteria (black), Proteobacteria (pink), Bacteroidetes (blue) and others (orange). Association of selected RiPPER-generated peptide networks to the HopA1 proteins is depicted as coloured strips surrounding the tree. Conserved pfam domains in the genetic context of each HopA1 proteins are shown as coloured dots in the outer rings of the tree. Tree visualised using iTOL^50^. See Supplementary Dataset 3 for a high-resolution tree that includes further mapping of peptide networks.

We hypothesised that if conserved short peptides were encoded near these proteins then they could represent novel RiPP BGCs. To assess this, we used RiPPER, which is a tool previously developed in our group to identify conserved short peptides encoded near a series of user-defined “bait” proteins^20^. Groups of related peptides were identified by generating sequence similarity networks^51^ with an identity cut-off of 40% (Figure S26). Using the 742 HopA1-like proteins as bait for a RiPPER analysis, we found that 77% of these HopA1-like proteins are associated with short peptide networks in the RiPPER output (Supplementary Dataset 1). The thioamitides themselves belong to peptide Family 10. These networks were then mapped to a HopA1-like protein phylogeny and the associated genomic loci were assessed for conservation (Figure 7). These putative BGCs were manually assessed for characteristic features of RiPP BGCs: co-linearity of putative biosynthetic genes and a position at the beginning of biosynthetic genes for the short peptide gene. A co-occurrence analysis^52^ was used to identify proteins commonly encoded in these putative BGCs (Figure 7).

Most of the putative precursor peptides from Cyanobacteria and Actinobacteria belong to two major sub-networks that are loosely related between themselves (Families 1A and 1B). The main precursor peptide network associated to Actinobacteria (272 precursor peptides, Family 1B) includes the precursor peptide for the antibiotic cacaoidin, the first member or the recently described lanthidin RiPP family^33^. Genes *cao7*, *cao9* and *caoD* in the cacaoidin BCG encode a HopA1-like protein, a phosphotransferase and a cysteine decarboxylase homologous to TsaD, TsaC and TsaF respectively, which suggests that the AviMeCys group found in this molecule is installed following a similar mechanism as in the thioamitides. Co-occurrence analysis shows that Family 1B BGCs also encode flavin-dependent oxidoreductases and peptidases, and occasionally other putative tailoring enzymes, such as glycosyltransferases and methyltransferases (Figure 8A). The precursor peptides in this network share higher conservation on their likely leader N-terminal region than in their C-terminal region (Figure S27), although the C-terminus contains a Ser/Ala rich region and two highly conserved Thr and Cys residues (Figure 8A), which supports the theory that these BGCs produce diverse AviMeCys containing RiPPs. In parallel with our study, a new RiPP genome mining algorithm, decRiPPter, also identifies the discovery of a similar set of actinobacterial RiPP BGCs encoding HopA1-like proteins and phosphotransferases^53^.

**Figure 8.**
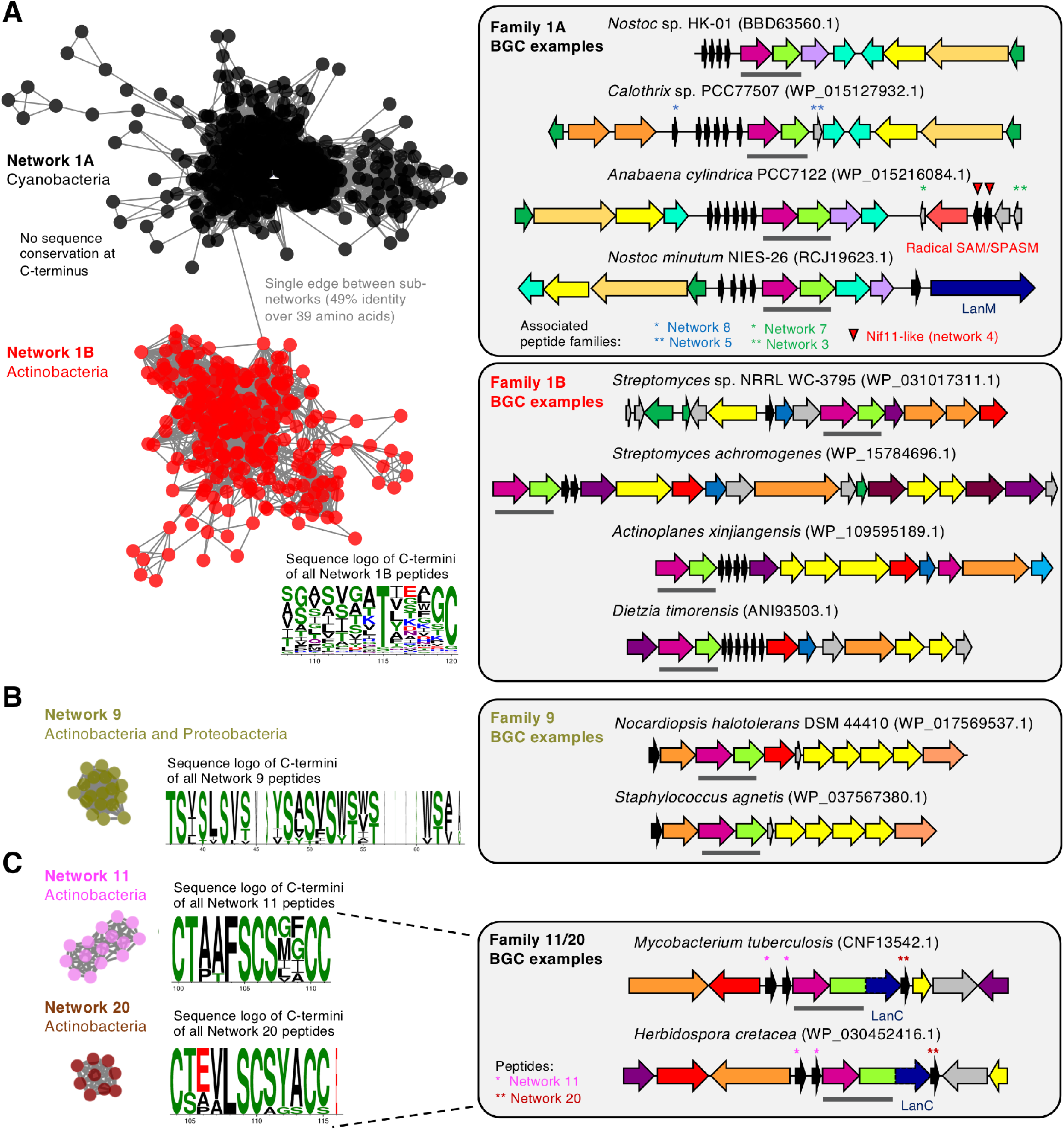
Selected HopA1-associated precursor peptides and examples of corresponding BGCs. Networks^54^ represent short peptide networking output from RiPPER with a 40% identity cut-off (Figure S26). Sequence logos^55^ are shown for selected portions of the C-termini of each family. In each BGC, the HopA1/phosphotransferase pair is highlighted with a grey bar and the HopA1-like protein accession is listed. See Figure 7 for a key to BGC colour-coding. A. Family 1A and 1B. B. Network 9. C. Network 11 and 20, which co-occur in the same BGCs. Additional associated peptide networks are highlighted (see Figure S33 for sequence logos).

Family 1A precursor peptides (570 peptides across 183 BGCs) are exclusive to Cyanobacteria and are encoded in partially conserved BGCs (Figure 8A). These BGCs also encode peptidylprolyl isomerases and peptidase-containing ABC transporter genes, which points to the coupling of leader peptide cleavage and peptide export, as in lanthipeptide biosynthesis^56^. These precursor peptides have high conservation of their leader region, which features a conserved double glycine motif that is a common cleavage motif in lanthipeptides^26^. In contrast, their C-terminal regions, which are predicted to correspond to the core peptide regions, are highly variable and do not feature conserved Ser, Thr or Cys residues (Figure S27). These BGCs typically encode multiple non-identical precursor peptides (up to 10), which is common occurrence in cyanobacterial RiPPs^57^. A small subset of these seem to be adjacent to additional RiPP BGCs containing Nif11-like precursor peptides and radical SAM maturases^58,59^, as well as to class II lanthipeptide clusters containing LanM proteins (Figure 8A). The next biggest peptide network identified by RiPPER (Family 2) is also exclusively cyanobacterial (Figure S28). The associated BGCs are conserved, although these short peptides contain a transmembrane protein domain (pfam03647), so it is unclear whether these represent true RiPP clusters.

A large group of HopA1-like proteins are associated with LanC-like cyclases (Figure 7). These are mostly found in Proteobacteria, but also in some Bacteroidetes and Actinobacteria. This further reinforces the role of the HopA1-phosphotransferase pair mediating the dehydration required for lanthionine bond formation, which could then be catalysed by LanC-like proteins or other cyclases. For example, peptides in network 22 feature conserved Cys and Ser/Thr residues and are found in proteobacterial BGCs that encode discrete LanC-like proteins (Figure S29). Supporting this functional association, there are instances of HopA1-like domains that are physically fused to either phosphotransferases or to the LanC like domain-containing proteins (Figure 8C and Figure S30). HopA1-LanC fusions could represent a new uncharacterised lanthionine synthetase, where the dehydratase and cyclase are fused in a single protein, as in LanM. Unexpectedly, this subgroup of proteins is not consistently associated with putative precursor peptides, particularly in Proteobacteria, which suggests that in some cases HopA1-like/phosphotransferases may be involved in the posttranslational modification of larger proteins or precursor peptides that are not clustered. There are some very good candidates for putative RiPP BGCs encoding HopA1-LanC fusions, such as the BGCs encoding peptide families 11 and 20, which appear in *Herbidospora*, *Actinomadura* and *Mycobacterium* species (Figure 8C and Figure S31). There are also further peptide networks that are associated with specific clades of HopA1-like proteins and represent likely RiPP BGCs, such as Family 9, which spans Actinobacteria and Proteobacteria. Family 9 peptides feature Ser/Thr rich C-termini and are encoded within conserved BGCs (Figure 8B and Figure S32).

## CONCLUSION

The apoptosis-inducing thioamitides represent some of the most complex RiPPs identified, where thiostreptamide S4 (**1**) contains 11 post-translational modifications (Figure 1A). BGC-wide gene deletions can be a powerful method to understand biosynthetic pathways. However, this process can be particularly complicated in RiPP pathways, where a partially modified precursor peptide may rapidly degrade if the pathway stalls in the absence of an essential modification step^12^. Here, we inactivated every gene in the thiostreptamide S4 BGC and used MS-based untargeted metabolomics and precursor peptide mutations to inform a model of how thioamitides are biosynthesised in the bacterial cell. Our analysis of the metabolites produced by expressing the thiostreptamide S4 BGC in *S. coelicolor* M1146 resulted in the identification of **2**-**16** (Table 1, Figure S17), mainly by detailed LC-MS/MS characterisation. This LC-MS characterisation was supported by detailed NMR characterisation of **12**, a key thioamidated and dehydrated shunt metabolite.

These data include a number of key findings about thioamitide biosynthesis, which enables a biosynthetic pathway to be proposed (Figure 6A). Our work confirms that YcaO and TfuA domain proteins (TsaH and TsaI) are required for iterative thioamidation and this functions as a gatekeeper for all subsequent biosynthetic steps. Prior studies of archaeal YcaO proteins indicate that this is an ATP-dependent process^24,60^. We define the proteins responsible for histidine hydroxylation and bis-methylation, as well as the reductase required for N-terminal reduction in the thioalbamide pathway. Yeast-mediated assembly provided a route to site-directed mutagenesis of the thiostreptamide S4 precursor peptide, which demonstrated that the pathway is tolerant to precursor peptide mutations, but does stall at an early stage in the biosynthetic pathway with some mutations. This indicates that macrocyclisation is a bottleneck for engineering thiostreptamide S4 biosynthesis.

We show that a phosphotransferase and a HopA1-like protein (TsaC and TsaD) are required for dehydration, which represents a new route to α,β-dehydroamino acids. Our results contrast with a recent study indicating that lanthipeptide synthetases encoded outside of the BGC catalyse dehydration in thioamitide biosynthesis^35^. Metabolomic results show that a further phosphotransferase (TsaE) is essential for biosynthesis, where it may have a role in either dehydration or macrocyclisation. A detailed informatic analysis using RiPPER^20^ shows that the phosphotransferase/HopA1-like protein pair defines multiple new RiPP BGC families, with representatives across over 1,000 sequenced genomes. The variety of tailoring enzymes and precursor peptide sequences indicates that the products will be highly diverse. This is supported by the parallel identification of HopA1-containing BGCs by the decRiPPter algorithm^53^, which have been recently defined as lanthidins in antiSMASH 5.0 ^61^.

Our insights are supported by parallel studies of individual enzymes in other thioamitide pathways^10,11^, as well as the recent discoveries of cacaoidin^33^ and lexapeptide^34^, which contain AviMeCys macrocycles, as predicted from our experimental and informatic analyses. We anticipate that the data reported here will inform further experimental work on the thioamitides and related RiPPs to determine key biosynthetic steps, including the true roles of HopA1 and TfuA domain proteins in their respective steps, given that neither protein features a known catalytic domain. Similarly, the true catalytic function of the HopA1 effector protein in *P. syringae* remains unknown^30^. More widely, understanding the diversity of products made by HopA1-like associated RiPP BGCs will be a substantial and exciting research effort, especially given the diversity of pathways identified.

## SUPPLEMENTARY MATERIAL

Supplementary Information (PDF): Methods, Supplementary Tables and Supplementary Figures

Supplementary Dataset 1 (CSV): Data for networked peptides from RiPPER

Supplementary Dataset 2 (Cytoscape file): Networked short peptides and associated data

Supplementary Dataset 3 (TIFF): High-resolution phylogenetic tree

Supplementary Dataset 4 (XLSX): Co-occurrence data for HopA1 proteins

## AUTHOR CONTRIBUTIONS

**Tom Eyles:** investigation, methodology, conceptualisation, visualisation, writing – original draft and review & editing. **Natalia Vior:** investigation, methodology, formal analysis, data curation, visualisation, writing – review & editing. **Rodney Lacret:** investigation, validation. **Andrew Truman:** project administration, supervision, funding acquisition, methodology, conceptualisation, visualisation, writing – original draft and review & editing.

## ACKNOWLEDGEMENTS

This work was funded by a Biotechnology and Biological Sciences Research Council (BBSRC) Norwich Research Park Doctoral Training Partnership grant (BB/M011216/1) for T.H.E., a Royal Society University Research Fellowship (A.W.T.), and BBSRC MET and MfN Institute Strategic Programme grants (BB/J004596/1 and BBS/E/J/000PR9790) for the John Innes Centre (JIC). We are very grateful for the technical assistance at JIC provided by Lionel Hill and Gerhard Saalbach for LC-MS analysis, Sergey Nepogodiev for NMR assistance, and Govind Chandra for assistance with informatics. We thank Vladimir Larionov (National Cancer Institute, NIH, USA) for *S. cerevisiae* VL6-48N and Bradley Moore (Scripps Institution of Oceanography, University of California San Diego, USA) for pCAP03. We are thankful to Javier Santos-Aberturas for helpful discussions.

