## Supplementary Information for "Understanding Thioamitide Biosynthesis Using Pathway Engineering and Untargeted Metabolomics"

### MATERIALS AND METHODS

#### Materials

Unless otherwise noted, all chemicals and media components were purchased from Sigma Aldrich. All restriction enzymes were purchased from New England Biolabs. The final concentrations of antibiotics used were: 50 µg $mL^{-1}$ kanamycin, 50 µg $mL^{-1}$ hygromycin, 50 µg $mL^{-1}$ carbenicillin, 25 µg $mL^{-1}$ chloramphenicol, and 25 µg $mL^{-1}$ nalidixic acid. All primers and oligonucleotides were ordered at a High Purity Salt Free (HPSF) grade from Eurofins Genomics. Ultrapure water was obtained using a Milli-Q purification system (Merck) and all media and solutions were autoclaved prior to use, unless otherwise stated.

#### Propagation and storage of strains

*Streptomyces coelicolor* M1146[1] was cultured on SFM (2% soy flour (Holland & Barrett), 2% mannitol, and 2% agar (Formedium)). Spore stocks were stored in 20% glycerol at -20 °C. *Escherichia coli* DH5α (Invitrogen) was used for plasmid propagation and *E. coli* ET12567[2] was used for conjugations; these were grown in lysogeny broth (LB) and stored in 20% glycerol at -20 °C. Electrocompetent *E. coli* was stored in 10% glycerol at -80 °C. *Saccharomyces cerevisiae* VL6-48N[3] was grown on YPAD (1% yeast extract, 2% peptone (BD Biosciences), 2% glucose (Fisher), 20 µg $mL^{-1}$ adenine, and 2% agar), and was stored in 20% glycerol at -80 °C, or as single colonies on YPAD plates at 7 °C.

#### Standard protocols

*Streptomyces* genomic DNA extraction was carried out using the salting out procedure (Kieser ref). Polymerase chain reactions (PCRs) of fragments for assemblies were carried out with Herculase II Fusion DNA Polymerase (Agilent) and analytical PCRs were carried out Go Taq G2 Flexi DNA Polymerase (Promega). These followed the manufacturers' protocols and annealing temperatures were based on predicted primer melting temperatures. Plasmid DNA extraction and purification was carried out using the Wizard plus SV Minipreps DNA purification system (Promega) and PCR products were purified using the Illustra GFX PCR DNA and gel band purification kit (GE Healthcare) following manufacturer instructions. All sequencing was carried out on plasmid miniprep templates using a Mix2Seq Kit (Eurofins Genomics). Sequences of all primers are listed in Table S2.

#### Bacterial transformations and conjugations

Electrocompetent *E. coli* were prepared using the following protocol. *E. coli* were inoculated into 10 mL SOB-Mg (2% tryptone, 0.5% yeast extract, 0.58% NaCl, and 0.186% KCl) and grown overnight at 250 rpm, 28 °C. A 1% inoculum of this starter culture was added to ten 250 mL conical flasks, each containing 50 mL SOB-Mg. These were incubated for 3 to 5 hours and at 250 rpm, 28 °C until an OD600 between 0.2 and 0.4 was reached. Cells were harvested by centrifugation at 800 x *g* for 20 minutes and resuspended in a total of 250 mL 10% glycerol. This was repeated three times, resuspending in totals of 50 mL, 50 mL, and 1 mL successively. This produced electrocompetent cells, which were separated into 50 µL aliquots in microcentrifuge tubes and flash frozen for storage at -80 °C. Electrocompetent *E. coli* were transformed using the following protocol: Electrocompetent cells were thawed and electroporation cuvettes (2 mm) were cooled on ice. 1 µL of DNA solution was added to each aliquot of cells. This was mixed gently by pipetting and transferred to the electroporation cuvette. The outside of the cuvette was dried, and it was inserted into a Gene Pulser (Bio-Rad) with the pulse generator set to 25 µFD, 2.5 kV, and 200 Ω. The pulse was delivered, and the cuvette was immediately placed on ice. 200 µL of LB was added. The cells were transferred back to a microcentrifuge tube and were incubated for an hour at 250 rpm, 37°C. The entire transformation mix was then plated on LB with the appropriate selection. If hygromycin selection was used cells were plated on DN-agar (2.3% Difco™ Nutrient Broth and 2% agar).

*E. coli* ET12567 carrying the helper plasmid pR9604[4] were used for intergenic conjugations into *Streptomyces* strains using an established protocol[5], with a few modifications as described here. *E. coli* ET12567-pR9604 transformants were selected on LB-agar containing carbenicillin, chloramphenicol, and either kanamycin or hygromycin for pCAP-based or pIJ10257-based plasmids, respectively. A single colony was used to inoculate 10 mL liquid LB containing the same antibiotics, and grown overnight at 37 °C, 250 rpm. A 1% inoculum was used to start a fresh growth in 10 mL of LB containing the same antibiotics. This was grown in the same conditions to an $OD_{600}$ of 0.4. Cells were washed twice in 2 mL LB and resuspended in 1 mL LB. 5 µL of *Streptomyces* spores were mixed with 0.5 mL 2xYT (1.6% tryptone (BD Biosciences), 1% yeast extract, and 0.5% NaCl; adjusted to pH 7.4 before autoclaving) and heat-shocked at 50 °C for 10 minutes. 0.5 mL of washed *E. coli* ET12567-pR9604 were added to the heat-shocked spores. This mixture was pelleted by centrifugation at 3000 x *g* for 5 minutes and resuspended in 0.2 mL water. Cells were plated on SFM with $MgCl_2$ (Fisher) added to a final concentration of 10 mM and incubated at 30 °C overnight. These plates were overlaid with nalidixic acid and kanamycin or hygromycin to select for *Streptomyces* pCAP-based or pIJ10257-based plasmid exconjugants, respectively.

**Yeast transformations**

Yeast transformations all followed a modified version a published lithium acetate/polyethylene glycol (LiAc/PEG) protocol[6]. Single colonies of *S. cerevisiae* VL6-48N were inoculated into 10 mL of liquid YPAD in a 50 mL conical centrifuge tube and grown overnight at 30 °C with shaking at 250 rpm. This starter culture was added to 40 mL liquid YPAD in a 250 mL conical flask and incubated for 4 hours at 250 rpm, 30 °C. These cells were harvested by centrifuging for 5 minutes at 1,789 x *g* and washed twice in equal volumes of sterile water. Cells were resuspended in 1 mL 0.1 M LiAc, transferred to a microcentrifuge tube and pelleted at 3,000 x *g* for 15 seconds. These cells were resuspended in 400 µL 0.1 M LiAc to a final volume of 500 µL and then transferred to microcentrifuge tubes as aliquots of 50 µL for single transformations. Prior to a transformation, an aliquot was briefly centrifuged, and the supernatant was removed. The following solutions were then added to the cells, in this order: 240 µL PEG solution (50% PEG 3350), 36 µL 1 M LiAc, 50 µL salmon sperm DNA (Invitrogen; 2 mg $mL^{-1}$; boiled for ten minutes and cooled in ice water), and 34 µL DNA to be assembled. Cells were resuspended by pipetting and incubated at 250 rpm, 30 °C for 30 minutes, which was followed by heat shock at 42 °C for 30 minutes. Cells were pelleted by centrifugation at 3,000 x *g* for 30 seconds, the supernatant was removed, and cells were resuspended in 200 µL sterile water. This final volume of 220 µL was separated into 200 µL and 20 µL aliquots that were plated on selective media SD+CSM-Trp (0.17% YNB-AA-$(NH_4)_2SO_4$ (Formedium), 0.5% $(NH_4)_2SO_4$, 2% glucose, 2% agar, 20 µg $mL^{-1}$ adenine and 740 µg $mL^{-1}$ CSM-TRP (Formedium)). Plates were then incubated for 3 days at 30 °C.

**Yeast colony screening**

To screen yeast colonies by PCR, colonies were picked using a pipette tip, and cells were resuspended in 50 µL 1 M sorbitol (Fisher). 2 µL of zymolyase (5 U $µL^{-1}$; Zymo Research) was added to each cell suspension and incubated at 30 °C for 1 hour. Cell suspensions were then boiled for 10 minutes, centrifuged for 15 seconds at 1,000 x *g* and 1 µL of the supernatant was used as a template for PCR.

**Yeast to *E. coli* plasmid shuttling**

To shuttle plasmids from yeast into *E. coli*, colonies of yeast were grown in 10 mL of liquid SD+CSM-TRP overnight at 250 rpm, 30 °C. Cells were harvested by centrifuging for 5 minutes at 1,789 x *g* and resuspended in 200 µL 1 M sorbitol plus 2 µL of zymolyase (5 U $\mu L^{-1}$). Cell suspensions were incubated at 30 °C for 1 hour to produce spheroplasts. Spheroplasts were pelleted at 600 x *g* for 10 minutes, and the supernatant was aspirated. Plasmid DNA was extracted from the spheroplasts using a standard Wizard miniprep protocol (Promega). 1 µL plasmid DNA was then transformed into *E. coli* by electroporation. Transformations were plated on LB-agar and transformants were selected for with kanamycin.

**Gene deletions**

To obtain gene deletions in pCAPtsa, PCR targeting was used following the published protocol[7]. The disruption cassette (apramycin resistance cassette bracketed by FLP-recombinase recognition sites) was amplified from pIJ773-oriT, a version of pIJ773 modified to have the oriT removed, with the following primers:

*tsaA* cassette = SPPCRtTsaA and ASPPCRtTsaA
*tsaC* cassette = SPPCRtTsaC and ASPPCRtTsaC
*tsaD* cassette = SPPCRtTsaD and ASPPCRtTsaD
*tsaE* cassette = SPPCRtTsaE and ASPPCRtTsaE
*tsaF* cassette = SPPCRtTsaF and ASPPCRtTsaF
*tsaG* cassette = SPPCRtTsaG and ASPPCRtTsaG
*tsaH* cassette = SPPCRtTsaH and ASPPCRtTsaH
*tsaI* cassette = SPPCRtTsaI and ASPPCRtTsaI
*tsaJ* cassette = SPPCRtTsaJ and ASPPCRtTsaJ
*tsaK* cassette = SPPCRtTsaK and ASPPCRtTsaK
*tsaMT* cassette = SPPCRtTsaMT and ASPPCRtTsaMT
*tsaL* cassette = SPPCRtTsaL and ASPPCRtTsaL
*tsaMO* cassette = SPPCRtTsaMO and ASPPCRtTsaMO
*tsa+1* cassette = SPPCRtTsa+1 and ASPPCRtTsa+1
*tsa+2* cassette = SPPCRtTsa+2 and ASPPCRtTsa+2
*tsa+3* cassette = SPPCRtTsa+3 and ASPPCRtTsa+3
*tsa-1*, *-2* and *-3* cassette = SPPCRtTsa-1,2,3 and ASPPCRtTsa-1,2,3

pCAPtsa was introduced to *E. coli* BW25113-pIJ790[7,8] by electroporation, and transformants were selected on LB-agar containing chloramphenicol and kanamycin at 30 °C. A single colony was used to make electrocompetent cells using the standard protocol (see *Bacterial transformations and conjugations* above), with two modifications: the subculturing on the second day was performed with 10 mM arabinose added to the medium, and the 10% glycerol washes were replaced with sterile water to facilitate immediate use, rather than flash freezing. These cells were then transformed with the gene specific disruption cassettes by electroporation. Transformants were selected for on LB-agar containing kanamycin and apramycin at 37 °C. Plasmids were extracted from *E. coli* colonies, and gene disruptions were first confirmed by PCR using the following primer pairs: Δ*tsaA* was assessed with SPTsaA and ASPTsaA; Δ*tsaC* was assessed with SPTsaC and ASPTsaC; Δ*tsaD* was assessed with SPTsaD and ASPTsaD; Δ*tsaE* was assessed with SPTsaE and ASPTsaE; Δ*tsaF* was assessed with SPTsaF and ASPTsaF; Δ*tsaG* was assessed with SPTsaG and ASPTsaG; Δ*tsaH* was assessed with SPTsaH and ASPTsaH; Δ*tsaI* was assessed with SPTsaI and ASPTsaI; Δ*tsaJ* was assessed with SPTsaJ and ASPTsaJ; Δ*tsaK* was assessed with SPTsaK and ASPTsaK; Δ*tsaMT* was assessed with SPTsaMT and ASPTsaMT; Δ*tsaL* was assessed with SPTsaL and ASPTsaL; Δ*tsaMO* was assessed with SPTsaMO and ASPTsaMO; Δ*tsa+1* was assessed with SPTsa+1 and ASPTsa+1; Δ*tsa+2* was assessed with SPTsa+2 and ASPTsa+2; Δ*tsa+3* was assessed with SPTsa+3 and ASPTsa+3; and Δ*tsa-1*,*-2*, and *-3* was assessed with SPTsa-1,2,3 and ASPTsa-1,2,3. Additionally, the primers SPaac(3)IVseq and ASPaac(3)IVseq were used to sequence outwards from the inserted selectable marker.

To remove the selectable marker, gene-disrupted pCAPtsa plasmids were transformed into *E. coli* DH5α-BT340[9] by electroporation. Transformants were selected for on LB-agar containing apramycin, chloramphenicol, and kanamycin at 30 °C. Individual colonies were picked and spread to single colonies on LB-agar without antibiotics and grown overnight at 42 °C. Isolated colonies were spread as patches on LB-agar containing kanamycin and then LB-agar containing kanamycin and apramycin. Patches that only grew on the LB-agar plates containing kanamycin were assumed to have lost the disruption cassette, leaving behind an 81 bp scar. This was additionally confirmed by sequencing using the following primers: Δ*tsaA* was assessed with SPTsaA and ASPTsaA; Δ*tsaC* was assessed with SPTsaC and ASPTsaC; Δ*tsaD* was assessed with SPTsaD and ASPTsaD; Δ*tsaE* was assessed with SPTsaE and ASPTsaE; Δ*tsaF* was assessed with SPTsaF and ASPTsaF; Δ*tsaG* was assessed with SPTsaG and ASPTsaG; Δ*tsaH* was assessed with SPTsaH and ASPTsaH; Δ*tsaI* was assessed with SPTsaI and ASPTsaI; Δ*tsaJ* was assessed with SPTsaJ and ASPTsaJ; Δ*tsaK* was assessed with SPTsaK and ASPTsaK; Δ*tsaMT* was assessed with SPTsaMT and ASPTsaMT; Δ*tsaL* was assessed with SPTsaL and ASPTsaL; Δ*tsaMO* was assessed with SPTsaMO and ASPTsaMO; Δ*tsa+1* was assessed with SPTsa+1 and ASPTsa+1; Δ*tsa+2* was assessed with SPTsa+2 and ASPTsa+2; Δ*tsa+3* was assessed with SPTsa+3 and ASPTsa+3; and Δ*tsa-1*,*-2*, and *-3* was assessed with SPTsa-1,2,3 and ASPTsa-1,2,3.

#### *In trans* expression of single genes

Constructs for the *in trans* expression of thioalbamide genes and thiostreptamide S4 complementation genes were assembled using standard digestion and ligation of PCR fragments into pIJ10257[10]. *taaRed* and *taaCYP* were amplified from *Amycolatopsis alba* DSM 44262 genomic DNA using the primer pairs: SPNdeITaaRed and ASPNdeITaaRed, and SPNdeITaaCYP and ASPPacITaaCYP, respectively. Thiostreptamide S4 genes *tsaA*, *tsaC*-*tsaJ* were amplified from pCAPtsa using the following primers to assess for the correct start codon (see Table S3 and Figure S4): for *tsaA* SPNdeITsaA, SPNdeITsaAv2, SPNdeITsaAv3 and ASPPacITsaA were used; for *tsaC* SPNdeITsaC, SPNdeITSaCv2, and ASPPacITsaC were used; for *tsaD* SPNdeITsaD, SPNdeITsaDv2, SPNdeITsaDv3, and ASPPacITsaD were used; for *tsaE* SPNdeITsaE and ASPPacITsaE were used; for *tsaF* SPNdeITsaF and ASPPacITsaF were used; for *tsaG* SPNdeITsaG, SPNdeITsaGv2, SPNdeITsaGv3, SPNdeITsaGv4, and ASPPacITsaG were used; for *tsaH* SPNdeITsaH and ASPPacITsaH were used; for *tsaI* SPNdeITsaI and ASPPacITsaI were used; for *tsaJ* SPNdeITsaJ and ASPPacITsaJ were used; and for *tsaMT* SPNdeITsaMT and ASPPacITsaMT were used.

PCR fragments and pIJ10257 were digested with NdeI and PacI. The PCR fragments were ligated in to pIJ10257 using T4 ligase (Invitrogen). Ligation reactions were transformed into *E. coli* DH5α by electroporation and transformants were selected for on DN-agar containing hygromycin. Colonies were screened for correct ligations by PCR using primers SPpIJ10257ins and ASPpIJ10257ins. Plasmids extracted from these colonies were additionally checked by sequencing using the same primers.

#### Precursor peptide mutations

Precursor peptide mutations were made to *tsaA* within pCAPtsa. These assemblies were all carried out using LiAc/PEG mediated transformation into yeast. Assemblies were designed in such a way as each core peptide modification was installed by a single oligonucleotide, which was linked to the backbone by a PCR fragment on each side. The backbone was AflII and SrfI-digested and gel-purified pCAPtsa. The parts used in each assembly are shown in Table S6, while Table S7 has a description of each part. A schematic of the assembly is shown in Figure S19.

The efficiency and flexibility of the system meant no strict adherence to concentration or molar ratios of DNA was necessary. As a guide, the following was sufficient for an assembly: 40 ng of linearized plasmid, 80 ng of each PCR product, and 500 pmol of each oligonucleotide. PCR products and linearised plasmids were purified as described in *Standard protocols* above. PCR products used in assemblies were produced with approximately 60 bp regions of overlap with other assembly parts. The oligonucleotides used in assemblies overlapped by between 30 and 60 bp, and if they were assembled to another oligonucleotide they were also complementary. Plasmid screening was accomplished using PCR and/or sequencing. The assembly of pCAPtsa *tsaA* mutant plasmids were assessed by PCR using the primers SPPTsaAseq and ASPPTsaCseq, and subsequently sequenced using the primers SPPTsaAseq, ASPPTsaCseq, and ASPSrfITsaCseq.

**Production cultures**

All production cultures were carried out using BPM-agar[11] (1% glucose, 1.5% starch (BD Biosciences), 0.5% yeast extract, 1% soy flour, 0.5% NaCl, 0.3% $CaCO_3$ (AnalaR), and 2% agar; adjusted to pH 7.0 before autoclaving; $CoCl_2$ was added after autoclaving to a final concentration of 20 µg $mL^{-1}$. 10 µL of spores were spread directly on BPM-agar and this was grown for 5 days at 28 °C.

**Liquid chromatography - mass spectrometry (LC-MS)**

Plugs were taken from production plates using the top of a 1 mL pipette and mixed with 500 µL methanol per plug. This was shaken for 1 hour at room temperature before being centrifuged at 20 000 x *g* for 15 minutes. The resulting supernatant was used for analysis. High throughput LC-MS/MS data were acquired using a Shimadzu Nexera X2 UHPLC connected to a Shimadzu ion-trap time-of-flight (IT-TOF) mass spectrometer and analysed using LabSolutions software (Shimadzu). 10 µL samples were injected onto a Phenomenex Kinetex 2.6 µm XB-C18 column (50 mm x 2.1 mm, 100 Å). The samples were eluted over 5 minutes using a 5 to 95% gradient of acetonitrile in water + 0.1% formic acid. After the first minute of each run, positive mode MS data were collected between *m/z* 200 and 2000, with an ion accumulation window of 20 ms and automatic sensitivity control of 70% of the base peak. The curved desolvation line (CDL) temperature was 250 °C and the heat block temperature was 300 °C. MS/MS data were collected between *m/z* 50 and 2000 in a data-dependent manner for parent ions between *m/z* 200 and 2000, using collision-induced dissociation energy of 50% and a precursor ion width of 3 Da. The instrument was calibrated using sodium trifluoroacetate cluster ions prior to every run.

LC-MS data acquired on the Shimadzu IT-TOF was additionally assessed using the statistical package Profiling Solution 1.1 to provide untargeted analysis. The following Profiling Solution parameters were used: ion *m/z* tolerance = 0.1 Da, ion intensity threshold = 50,000, LabSolutions compatible ion *m/z* tolerances = ON, De-Isotope matrix = ON. Metabolites detected in the negative controls were filtered out during post-processing in Microsoft Office 365 ProPlus Excel. For each quantification experiment, the relevant metabolite was measured by integration of LC–MS peak areas using Browser software (Shimadzu). MS quantification data are the average of triplicate cultures. Error bars on all graphs represent the standard error.

High resolution LC-MS/MS data were acquired by Gerhard Saalbach (John Innes Centre) using a Waters Synapt G2-Si mass spectrometer, and analysed using MassLynx software (Waters). The Synapt G2-Si was operated in positive mode with a scan time of 0.5 s in the mass range of *m/z* 50 to 1200. 7 µL samples were injected onto a Phenomenex Luna Omega 1.6 µm Polar C18 column (50 mm x 2.1 mm, 100 Å) and eluted with a linear gradient of 1 to 50 % acetonitrile in water + 0.1% formic acid over 20 minutes. Synapt G2-Si MS data were collected with the following parameters: capillary voltage = 3 kV; cone voltage = 40 V; source temperature = 120 °C; desolvation temperature = 350 °C. Leu-enkephalin peptide was used to generate a dual lock-mass calibration with *m/z* = 278.1135 and *m/z* = 556.2766 measured every 30 s during the run.

**Purification and NMR analysis of 12**

**12** was purified from *S. coelicolor* M1146-pCAPtsaΔ*tsaE*, where LC-MS was used to assess the progress of purification through much of the purification. 100 µL of spores were spread on 1.6 L of BPM-agar and were incubated at 28 °C for five days. This was extracted with 3.2 L of ethyl acetate, which resulted in an organic fraction containing a small proportion of **12** and the remaining solid material containing the rest of **12**. The solid material was subsequently extracted with 3.2 L of methanol and reduced on the rotary evaporator at 30 °C to 800 mL. A liquid-liquid extract with 3.2 L of ethyl acetate was performed and the remainder of **12** partitioned into the organic phase. This organic phase was combined with the original ethyl acetate extract and was evaporated on a rotary evaporator at 30 °C to produce a brown solid. This was resuspended in $CHCl_3$ and dried onto silica gel. This was packed into a silica column and separated into 12 fractions using $CHCl_3$ and an increasing concentration of $CH_3OH$. An initial 400 mL elution of 100% $CHCl_3$ was followed by 10 elutions of 200 mL, ranging from 5% to 25% $CH_3OH$ in $CHCl_3$. The final elution was with 100% $CH_3OH$. Fractions eluted with 10% $CH_3OH$ and upwards contained **12**, so were combined and evaporated to a brown solid using a rotary evaporator at 30 °C. This was resuspended in $CH_3OH$, dried onto a 1 g DSC-18 column (Discovery), and eluted with a step-wise gradient of water:acetonitrile. 14 elutions of 10 mL, ranging from 0% to 60% acetonitrile, were collected. All fractions with acetonitrile concentrations equal to or less than 20% contained **12**, and so were combined and evaporated using a rotary evaporator at 30 °C to give 493 mg of brown solid. This was resuspended in $CH_3OH$ and separated by HPLC on a Gemini-NX 2.6 µ C18 column (150 mm x 21.2 mm, 100 Å) using a gradient from 10% to 60% acetonitrile over 45 minutes. Compound collection was guided by absorbance at 272 nm (characteristic of thioamide bonds). **12** was collected and evaporated using a Genevac EZ-2 Elite. This provided 0.7 mg of pure **12** as a white powder.

NMR spectra were recorded on a Bruker Avance 600 MHz NMR spectrometer. Chemical shifts were reported in ppm using the signals of the residual solvents as internal references ($\delta_H$ 3.31 and $\delta_C$ 49.0 for $CD_3OD$). Data were analysed using Mnova 6.0.2 (Mestrelab). See Figures S11 - S16 for NMR spectra.

### BIOINFORMATIC METHODS

#### Identification and RiPPER analysis of HopA1 domain-containing proteins

TsaD (WP_159037419.1) was used as query for a BlastP search on the non-redundant NCBI database applying default parameters and setting the maximum possible number of hits to 1000. This search returned 854 hits, from which 8 selected entries were used for subsequent BlastP searches to capture additional HopA1 domain-containing proteins diversity: BAN83919.1 (TvaD, *Streptomyces olivoviridis*), WP_161634174.1 (*Nocardiopsis potens*), WP_012409787.1 (*Nostoc punctiforme*), WP_075693282.1 (*Streptomyces acidiscabies*), WP_008178764.1 (*Moorea producens*), WP_015197629.1 (*Calothrix parietina*), AGP61345.1 (*Streptomyces rapamycinicus* NRRL 5491), WP_151696789.1 (*Mycrocistis aeruginosa*) and WP_109595189.1 (*Actinoplanes xinjiangensis*). These searches yielded a total of 1384 non-redundant entries, whose sequences were retrieved from the "protein" and "identical group proteins" databases using Batch Entrez (https://www.ncbi.nlm.nih.gov/sites/batchentrez).

The resulting sequences were size-filtered to remove any proteins shorter than 150 amino acids to yield a dataset with 1340 sequences. This dataset was then submitted to the online CD-HIT suite[12] for identity-based filtering with a 95% identity cut-off to reduce redundancy in downstream analyses. This yielded a reduced dataset with 828 sequences whose identity to each other was lower than 95%. The accession numbers of these sequences were then used as input for RiPPER analysis[13], which was run in Docker using default RiPPER parameters (minPPlen = 20, maxPPlen = 120, flanklen = 17.5, sameStrandReward = 5, maxDistFromTE = 8, fastaOutputLimit = 3, prodigalScoreThresh = 7.5). RiPPER retrieved information for 742 of the 828 entries, as the remaining 86 did not have associated nucleotide information. The RiPPER output was then used for downstream analyses, including short peptide networking with EGN[14] (thresholds = 40% identity, 40% sequence coverage), which was visualised using Cytoscape[15] (version 3.8.0). The RiPPER output is provided as Supplementary Datasets 1 (retrieved peptides) and 2 (Cytoscape file of networked peptides). Peptides belonging to each of the main networks described in the text were further analysed performing multiple sequence alignment with MUSCLE[16]. The resulting alignment files were then visualised for residue conservation across within each peptide network using

WebLogo[17] (version 3.7.4) with the following parameters: logo size = medium, units = probability, scale stacks width = on, no adjustment for composition, color scheme = chemistry.

**Phylogenetic analyses of HopA1 domain-containing proteins**

The set of successful RiPPER entries (742 sequences) was used to carry out a phylogenetic analysis of the HopA1 domain-containing proteins. As an outgroup for this analysis, the HopA1 protein from *Pseudomonas syringae* pv. *syringae* (AAF71481.2) was added to the dataset. Multiple sequence alignment and tree construction for this dataset were carried out using MUSCLE[16] and RAxML[18] respectively, through the CIPRES science gateway[19]. The following MUSCLE parameters were used:

```
muscle -in infile.fasta -seqtype protein -maxiters 2 -maxmb 30000000 -hydro 5 -hydrofactor 1.2 -log logfile.txt -verbose -weight1 clustalw -distance1 kmer6_6 -cluster1 upgmb -sueff 0.1 -root1 pseudo -maxtrees 1 -weight2 clustalw -distance2 pctidkimura -cluster2 upgmb -sueff 0.1 -root2 pseudo -objscore sp -noanchors -fastaout output.fasta
```

The FASTA output from the MUSCLE alignment was then used as input for maximum likelihood tree construction with RAxML using the following parameters:

```
raxmlHPC-HYBRID -T 4 -N autoMRE -n HopA1_filtered_RiPR -s infile.txt -p 12345 -m PROTGAMMABLOSUM62 -k -f a -x 12345 -o AAF71481.2 HopA1 [Pseudomonas syringae pv. syringae] --asc-corr lewis
```

The resulting maximum likelihood tree was visualised and edited in iTOL[20] to map the most prevalent precursor peptide networks identified by RiPPER and protein conserved domains associated to the HopA1 proteins in the tree. Upon tree plotting, four outlier proteins were identified and removed from the final analysis (KXJ59479.1, WP_12384669.1, WP_141234752.1 and PNO53249.1).

**Genetic context analysis and conserved domain mapping**

The genetic context of putative BGCs was assessed by using the output files from the RiPPER analysis described above. The GenBank files containing the annotated region surrounding the genes of interest were used to generate local databases for MultiGeneBlast analysis[21]. Separate databases were created for each of the most prevalent short peptide networks. These databases were then queried (default MultiGeneBlast settings) with a representative example GenBank file from each network to assess for genetic conservation and therefore determine the composition of putative biosynthetic gene clusters associated to each peptide network.

In addition, the "main_co_occur.csv" files (containing co-occurring conserved pfam domains encoded in the genetic region of the query protein) generated by RODEO2[22] (incorporated into RiPPER) for each of the protein accessions were merged into a single file for conserved domain analysis. This file was then used to determine the most abundant pfam domains co-occurring with the HopA1 domain-containing proteins across the full dataset using the instructions described at http://ripp.rodeo/advanced.html. The 35 most abundant pfam domains, plus other selected pfam domains (see Supplementary Dataset 4 for full list) were then mapped to each of the HopA1 proteins to determine co-occurrence patterns. For those entries where a phosphotransferase pfam domain was not identified by this mapping process, the proteins encoded immediately upstream of the HopA1 gene were retrieved and individually inspected to detect putative conserved pfam domains which might have been above the previous E-value cut off threshold (1x $10^{-3}$) as well as conserved domains from non-pfam databases and annotated accordingly. For ease of visualisation, results for pfam domains corresponding to similar activities (e.g. transport-related domains) were merged prior to mapping to the HopA1 phylogenetic tree using the iTOL annotation editor.

### SUPPLEMENTARY TABLES

**Table S1** Representative BLASTP and pfam matches for each protein encoded in the thiostreptamide S4 BGC. Results shown correspond to the first non-identical hit to TsaA and to the top matching hits to the tailoring enzymes not found in putative thioamitide gene clusters.

| Query protein | Accession | Homologous gene annotation | Accession | Organism | Coverage/ Identity (%) | Conserved pfam domain |
|---|---|---|---|---|---|---|
| **TsaA** | WP_107418454 | Thioviridamide family RiPP peptide | WP_018962002* | *Streptomyces* sp. CNB091* | 100/92 | Thiovirid_RiPP NF033415** |
| **TsaC** | WP_158709571 | Phosphotransferase | WP_157987422 | *Jiangella* sp. KE 2-3 | 62/31 | APH pfam01636 |
| **TsaD** | WP_159037419 | Hypothetical protein | WP_057178057 | *Cylindrospermopsis* (multispecies) | 97/31 | HopA1 pfam17914 |
| **TsaE** | WP_053929452 | Phosphotransferase family protein | WP_007508017 | *Frankia* (multispecies) | 27/35 | APH pfam01636 |
| **TsaF** | WP_051834497 | Hypothetical protein | WP_052035990 | *Tumebacillus flagellatus* | 79/45 | Flavoprotein pfam02441 |
| **TsaG** | WP_031090931 | Methyltransferase domain-containing protein | RPA66618 | Cyclobacteriaceae bacterium YHN15 | 95/33 | PrmA pfam06325 |
| **TsaH** | WP_031090932 | YcaO-like family protein | WP_082115066 | *Lentzea aerocolonigenes* | 95/56 | YcaO pfam02624 |
| **TsaI** | WP_051834498 | Hypothetical protein | WP_045314885 | *Lentzea aerocolonigenes* | 85/59 | TfuA pfam07812 |
| **TsaJ** | WP_051834499 | Phytanoyl-CoA dioxygenase family protein | WP_069779378*** | *Streptomyces fodineus* | 99/69 | PhyH pfam05721 |
| **TsaK** | WP_031090940 | C1 family peptidase | WP_069779379*** | *Streptomyces fodineus* | 95/66 | Peptidase_C1 pfam00112 |
| **TsaMT** | WP_078850506 | Methyltransferase domain-containing protein | WP_158089767 | *Mycobacterium sherrisii* | 92/38 | Methyltransf_25 pfam13649 |
| **TsaL** | WP_042836196 | Hypothetical protein | WP_142154560 | *Streptomyces* sp. SLBN-31 | 94/58 | None |
| **TsaMO** | WP_031090946 | LLM class flavin-dependent oxidoreductase | WP_103938723 | *Actinomadura echinospora* | 97/60 | Bac_luciferase pfam00296 |

* Identical hits to TsaA found in *Streptomyces* sp. NRRL S-15 and *Streptomyces mutomycini* NRRL B-65393.
** Not a pfam domain.
*** Could correspond to a truncated thioamitide cluster.

**Table S2** Primers used in this study.

| Primer/Oligonucleotide | Sequence (5’-3’) |
|---|---|
| SPPCRtTsaA | ggttgaagcatgtctgaa accac gac agca gctca ggt gattcc ggg gatcc gtcg acc |
| ASPPCRtTsaA | ccttctgctcggaagcgctgcttgccatt gtcta actc atgt aggc tgg agct gcttc |
| SPPCRtTsaC | gttgcgattgatgcggcatttcg ga atg agg ag gcga tga ttccgg gg atccgtc gacc |
| ASPPCRtTsaC | ctgcgtcgtatctgctcatgcgcggctgctcc gccccttt gta ggctg ga gctgcttc |
| SPPCRtTsaD | caccgggagtggggtgacctgtcccc tgctgc tgg ggtg attcc ggg gatcc gtcg acc |
| ASPPCRtTsaD | cggtctcctgggtgctcatatgga ggtcg ag gggc gggct gta ggct gga gctgct tc |
| SPPCRtTsaE | gcgcccctatctcaacgcccgcccctcgacctccat atg attccg ggg atccgtc gacc |
| ASPPCRtTsaE | cctcagcggtctctttcatgcagtggtc acccttcg gta tgt aggct gg agctgc ttc |
| SPPCRtTsaG | accagtaacgaggtgaa gatcc acttc ga aggcc gggt gat tccgg gga tccgtcg acc |
| ASPPCRtTsaG | agtttcactggtgctccttgcgtggt ggt atcgg aa tcat gta ggctg ga gctgcttc |
| SPPCRtTsaH | cggcctgattccgataccaccacgca agg agc acca gtg attccg ggg atccgtc gacc |
| ASPPCRtTsaH | cggtggtggggatgctcatgcg ttgctctcc tgctcgcc tgt aggct gg agctgc ttc |
| SPPCRtTsaI | gcagcagcgcaccacgggcgagc ag gag agc aac gcat gattcc ggg gatcc gtcg acc |
| ASPPCRtTsaI | tgttcaccgccgtcctcatgccgctcctcgactgtc agct gta ggct gga gctgct tc |
| SPPCRtTsaK | ccggatcgaccacaccgtgcggcaccg ga gcacg aca tga ttccgg gg atccgtc gacc |
| ASPPCRtTsaK | atgcagcttctgccgtcatgccgtctgctccc agtcctc tgt aggct gg agctgc ttc |
| SPPCRtTsaL | gtgattctcttacaggttgtgctg gtgt ggtcc gag gtg attccg ggg atccg tcgacc |
| ASPPCRtTsaL | cagtttccggcaacggtgcgcccgg aac actgcg ggct atgt ag gctgg agct gcttc |
| SPPCRtTsaMO | cttgagaaaccgtcaggacgc ttcta aa attct gccat gat tccgg gga tccgtcg acc |
| ASPPCRtTsaMO | tcgcagggatgggatcgggc ggtcg gccgg gccgg atc atgt aggc tgg agct gcttc |
| SPPCRtTsa+1 | acggacgagcggaaggg tacg gggc ggtt ggccgcc gtg attccg ggg atccg tcgacc |
| ASPPCRtTsa+1 | cgtggttcaagggcacgcggtg gacc ggcac ggcg gcta tgt aggct gg agctgc ttc |
| SPPCRtTsa+2 | ggggagatgtctttcatgacc gtcg agcac ga gggc atg attccg ggg atccg tcgacc |
| ASPPCRtTsa+2 | cgtcgggcaggaggggctggc agg ag gggct gccgg tcat gta ggctg ga gctgcttc |
| SPPCRtTsa-1,2,3 | tcggtttgacgcctcccatggtata aat agt ggctc gag attcc ggg gatcc gtcg acc |
| ASPPCRtTsa-1,2,3 | ctgtggcgccgactcgttccggacga agt gtgg gggc ggt gta ggctg ga gctgcttc |
| SPPCRtTsa+3 | acgcgtccgccgggccggctcatcgcaccg gcgcg acgc attccg ggg atccgtc gacc |
| ASPPCRtTsa+3 | agtagcagcacgttccttatat gta gctttc gac atat gtgt ag gctgg agct gcttc |
| SPPCRtTsaF | cctcgccgccgtcgggtaccgaagggt gacc actgc atg attccg ggg atccg tcgacc |
| ASPPCRtTsaF | tggtcatgtctgggatcactgtccttc tgctcgc gcggc tgt aggct gg agctgc ttc |
| SPPCRtTsaJ | ggatgtgccacctcccgctgacagtcg ag gagc ggc atg attccg ggg atccgtc gacc |
| ASPPCRtTsaJ | cggccgtgctcgtgctcatgtcgtgctccgg tgccgc actgt ag gctgg agct gcttc |
| SPPCRtTsaMT | ctggatagtgatcgaag agg actg gga gca gacg gcat gat tccgg gga tccgtcg acc |
| ASPPCRtTsaMT | agaatcactgccggggcggccca acgt gccgccgcc ttat gta ggct gga gctgct tc |
| SPPTsaAseq | gcacccgagaggtttcagc |
| ASPPTsaCseq | ccatgacgagggttccactc |
| ASPSrfITsaCseq | gccgggctcggggtatc |
| SPTsaA | ggttgaagcatgtctgaa ac |
| ASPTsaA | ccttctgctcggaagcgctg |
| SPTsaC | gttgcgattgatgcggcatt |
| ASPTsaC | cctccgccatgctggcgaac |

| | |
|---|---|
| SPTsaD | caccgggagtggggtgacct |
| ASPTsaD | gcggtcgctcccggaggttc |
| SPTsaE | gcgcccctatctcaacgccc |
| ASPTsaE | gacagcgcctgctgtatctc |
| SPTsaF | tgcgtgcatgggcggcatc |
| ASPTsaF | gcttgagcacccggccttc |
| SPTsaG | accagtaacgaggtgaagat |
| ASPTsaG | agtttcactggtgctccttg |
| SPTsaH | cggcctgattccgataccac |
| ASPTsaH | cggtggtggggatgctcatg |
| SPTsaI | gcagcagcgcaccacgggcg |
| ASPTsaI | tgttcaccgccgtcctcatg |
| SPTsaJ | ggatgtgccacctcccgctg |
| ASPTsaJ | cggccgtgctcgtgctcatg |
| SPTsaK | aggcccgtcccctctc |
| ASPTsaK | atctcgccactgatacgtcg |
| SPTsaMT | ctggatagtgatcgaagagg |
| ASPTsaMT | agaatcactgccggggcggc |
| SPTsaL | gtgattctcttacaggttgt |
| ASPTsaL | cagtttccggcaacggtgcg |
| SPTsaMO | cttgagaaaccgtcaggacg |
| ASPTsaMO | tcgcagggatgggatcgggc |
| SPTsa+1 | acggacgagcggaagggtac |
| ASPTsa+1 | cgtggttcaagggcacgcgg |
| SPTsa+2 | gtcgcgtcggccgggtac |
| ASPTsa+2 | cacggcgaggacaagacggt |
| SPTsa-1,2,3 | ctctctcgctttcctcattc |
| ASPTsa-1,2,3 | gagaaatgcgtgggctctc |
| SPTsa+3 | acgcgtccgccgggccggct |
| ASPTsa+3 | agtagcagcacgttccttat |
| SPseqTsaG | accaagctctcaccacag |
| SPseqTsaH | ggcagcaagtcaaccttg |
| SPseqTsaI | gccgggcgcggagtcctg |
| SPseqTsaK | cgaggcccgtcccctctc |
| SPseqTsaL | agaccgtgaagaaatacg |
| SPseqTsa+2 | tgttccctgtctatgctg |
| SPseqTsaMT | gccctacgcctttctgac |
| SPaac(3)IVseq | cgctacggaaggagctgtgg |
| ASPaac(3)IVseq | cttcctgcatccgccagagg |
| SPNdeITsaA | ggcccccatatggacgaggtcgcgttcagcga |
| ASPPacITsaA | ctctagttaattaagctgcttgccattgtctaac |

| | |
|---|---|
| SPNdeITsaC | ggcccccatatgttgcgtctggtgcctgtcgc |
| ASPPacITsaC | ctctagttaattaagaacactgcgtcgtatctgc |
| SPNdeITsaD | ggcccccatatggagacagagccgggcactct |
| ASPPacITsaD | ctctagttaattaacccccggtctcctgggtgctcagatg |
| SPNdeITsaE | ggcccccatatgagcacccaggagaccggggg |
| ASPPacITsaE | ctctagttaattaacggtacctcagcggtctctt |
| SPNdeITsaG | ggcccccatatgctcaagctgggccagtccgc |
| ASPPacITsaG | ctctagttaattaacttgcgtggtggtatcggaa |
| SPNdeITsaH | ggcccccatatgaaactgcgccagagcgtccc |
| ASPPacITsaH | ctctagttaattaaaccgacggtggtggggatgc |
| SPNdeITsaI | ggcccccatatgagcatccccaccaccgtcgg |
| ASPPacITsaI | ctctagttaattaacaggttgttcaccgccgtcc |
| SPNdeITsaK | ggcccccatatgagcacgagcacggccgatcc |
| ASPPacITsaK | ctctagttaattaatgaagatgcagcttctgccg |
| SPNdeITsaL | ggcccccatatgctgtccgtcggccaggccgt |
| ASPPacITsaL | ctctagttaattaatgcgcccggaacactgcggg |
| SPNdeITsaMO | ggcccccatatgaatattcccttctccgtgct |
| ASPPacITsaMO | ctctagttaattaatccggcccggccgaccgccc |
| SPNdeITsa+1 | ggcccccatatgtgggtgttcatgaccggtgg |
| ASPPacITsa+1 | ctctagttaattaagcggtggaccggcacggcgg |
| SPNdeITsa+2 | ggcccccatatggctgtggacgccggggcggg |
| ASPPacITsa+2 | ctctagttaattaactggcaggaggggctgccgg |
| SPNdeITsaJ | ggcccccatatgaggacggcggtgaacaacct |
| ASPPacITsaJ | ctctagttaattaagggatcggccgtgctcgtgc |
| SPNdeITsaF | ggcccccatatgaaagagaccgctgaggtacc |
| ASPPacITsaF | ctctagttaattaagagcttggtcatgtctggga |
| SPNdeITsaMT | ggcccccatatgacggcagaagctgcatcttc |
| ASPPacITsaMT | ctctagttaattaacggcccaacgtgccgccgcc |
| SPNdeITsaAv2 | ggcccccatatgtctgaaaccacgacagcagc |
| SPNdeITsaAv3 | ggcccccatatggaaacccggcgacaacaacg |
| SPNdeITSaCv2 | ggcccccatatggttagacaatggcaagcagc |
| SPNdeITsaDv2 | ggcccccatatgagcagatacgacgcagtgtt |
| SPNdeITsaDv3 | ggcccccatatgttcgccagcatggcggagga |
| SPNdeITsaGv2 | ggcccccatatgatcccagacatgaccaagct |
| SPNdeITsaGv3 | ggcccccatatgaccaagctctcaccacagac |
| SPNdeITsaGv4 | ggcccccatatgaagatccacttcgaaggccg |
| SPNdeITaaCYP | ggcccccatatgatgacggaaaccgtgaatc |
| ASPPacITaaCYP | ctctagttaattaatcagcgtggcgtgaagag |
| SPNdeITaaRed | ggcccccatatgatgaccggcgagaaccag |
| ASPNdeITaaRed | ctctagttaattaatcaccggctcgtgctgctg |
| SPpIJ10257ins | cacgcggtcgatcttg |
| ASPpIJ10257ins | cgagctgaagaaagacaatc |

| | |
|---|---|
| SPTsaAassembly | ccgcacccgagaggtttcag |
| ASPTsaAassembly | gccttgcgcctcctcgtccg |
| TsaCoreTaa | ggagaaggccggcatatcgccggacgaggaggcgcaaggcagcgtcatcggcttcgcggtcaccatcgccgtgcactgctgagttagacaatggcaa gcagcgcttccgagcagaaggc |
| S1T | ggagaaggccggcatatcgccggacgaggaggcgcaaggcaccgtcatggccgccatcgccaccgtggcttaccactgctgagttagacaatggcaa gcagcgcttccgagcagaaggc |
| SAV | gagaaggccggcatatcgccggacgaggaggcgcaaggcagcgccgtcatggccgccatcgccaccgtggcttaccactgctgagttagacaatggc aagcagcgcttccgagcagaagg |
| M3I | ggagaaggccggcatatcgccggacgaggaggcgcaaggcagcgtcatcgccgccatcgccaccgtggcttaccactgctgagttagacaatggcaa gcagcgcttccgagcagaaggc |
| A5F | ggagaaggccggcatatcgccggacgaggaggcgcaaggcagcgtcatggccttcatcgccaccgtggcttaccactgctgagttagacaatggcaa gcagcgcttccgagcagaaggc |
| T8S | ggagaaggccggcatatcgccggacgaggaggcgcaaggcagcgtcatggccgccatcgcctccgtggcttaccactgctgagttagacaatggcaa gcagcgcttccgagcagaaggc |
| Y11V | ggagaaggccggcatatcgccggacgaggaggcgcaaggcagcgtcatggccgccatcgccaccgtggctgtccactgctgagttagacaatggcaa gcagcgcttccgagcagaaggc |
| H12A | ggagaaggccggcatatcgccggacgaggaggcgcaaggcagcgtcatggccgccatcgccaccgtggcttacgcctgctgagttagacaatggcaa gcagcgcttccgagcagaaggc |
| H12W | ggagaaggccggcatatcgccggacgaggaggcgcaaggcagcgtcatggccgccatcgccaccgtggcttactggtgctgagttagacaatggcaa gcagcgcttccgagcagaaggc |
| SPTsaCassembly | tgagttagacaatggcaagc |
| ASPTsaCassembly | ccagccgggctcggggtatc |

**Table S3** Details of complementation experiments.

| Gene | Sense Primer | Antisense Primer | Experiment | Result |
|---|---|---|---|---|
| *tsaA* | SPNdeITsaA | ASPPacITsaA | Complementation of *tsaA* deletion from potential start codon 1 | Complementation failed |
| | SPNdeITsaAv2 | ASPPacITsaA | Complementation of *tsaA* deletion from potential start codon 2 | Complementation failed |
| | SPNdeITsaAv3 | ASPPacITsaA | Complementation of *tsaA* deletion from potential start codon 3 | Complementation failed |
| *tsaC* | SPNdeITsaC | ASPPacITsaC | Complementation of *tsaC* deletion from potential start codon | Complementation failed |
| | SPNdeITSaCv2 | ASPPacITsaC | Complementation of *tsaC* deletion including the native promoter | **Complementation succeeded** |
| *tsaD* | SPNdeITsaD | ASPPacITsaD | Complementation of *tsaD* deletion from potential start codon 1 | Complementation failed |
| | SPNdeITsaDv2 | ASPPacITsaD | Complementation of *tsaD* deletion from potential start codon 2 | **Complementation succeeded** |
| | SPNdeITsaDv3 | ASPPacITsaD | Complementation of *tsaD* deletion from potential start codon 3 | Complementation failed |
| *tsaE* | SPNdeITsaE | ASPPacITsaE | Complementation of *tsaE* deletion from potential start codon | **Complementation succeeded** |
| *tsaF* | SPNdeITsaF | ASPPacITsaF | Complementation of *tsaF* deletion from potential start codon | **Complementation succeeded** |
| *tsaG* | SPNdeITsaG | ASPPacITsaG | Complementation of *tsaD* deletion from potential start codon 1 | Complementation failed |
| | SPNdeITsaGv2 | ASPPacITsaG | Complementation of *tsaD* deletion from potential start codon 2 | **Complementation succeeded** |
| | SPNdeITsaGv3 | ASPPacITsaG | Complementation of *tsaD* deletion from potential start codon 3 | **Complementation succeeded** |
| | SPNdeITsaGv4 | ASPPacITsaG | Complementation of *tsaD* deletion from potential start codon 4 | Complementation failed |
| *tsaH* | SPNdeITsaH | ASPPacITsaH | Complementation of *tsaH* deletion from potential start codon | **Complementation succeeded** |
| *tsaI* | SPNdeITsaI | ASPPacITsaI | Complementation of *tsaI* deletion from potential start codon | **Complementation succeeded** |
| *tsaJ* | SPNdeITsaJ | ASPPacITsaJ | Complementation of *tsaJ* deletion from potential start codon | **Complementation succeeded** |
| *tsaMT* | SPNdeITsaMT | ASPPacITsaMT | Complementation of *tsaMT* deletion from potential start codon | **Complementation succeeded** |
| *taaRED* | SPNdeITaaRed | ASPNdeITaaRed | Elucidating pyruvyl reduction in thioalbamide biosynthesis | **TaaRed was able to reduce the thiostreptamide S4 pyruvyl** |
| *taaCYP* | SPNdeITaaCYP | ASPPacITaaCYP | Elucidating phenylalanine hydroxylation in thioalbamide biosynthesis | No hydroxylation of thiostraptamide S4 was seen |

**Table S4** Untargeted metabolomics with P-values lower than 1E[-4] for the wild type cluster and clusters with deletions to *tsaC*, *tsaD*, *tsaE*, and *tsaF*. The value and intensity of shading in the cells reflects the MS intensity. Ion *m\z* cells are shaded green if their structure was proposed in this study. Values represent the average of three biological replicates.

| Ion *m/z* | Ion RT | PVal | Δ*tsaC* | Δ*tsaD* | Δ*tsaE* | Δ*tsaF* | WT |
|---|---|---|---|---|---|---|---|
| 1377.55 | 3.1 | 5.7E-16 | 0 | 0 | 0 | 0 | 696481 |
| 465.22 | 2.0 | 2.6E-15 | 0 | 0 | 823330 | 478565 | 126777 |
| 392.11 | 1.9 | 3.1E-15 | 783547 | 759679 | 830419 | 538018 | 428295 |
| 552.23 | 2.2 | 9.6E-14 | 0 | 0 | 271762 | 160966 | 0 |
| 370.12 | 1.9 | 1.9E-13 | 150997 | 136938 | 0 | 0 | 0 |
| 348.14 | 1.9 | 2.2E-13 | 168199 | 186354 | 0 | 0 | 0 |
| 493.73 | 2.0 | 2.6E-13 | 0 | 0 | 0 | 0 | 140435 |
| 487.73 | 1.6 | 2.1E-12 | 0 | 0 | 0 | 0 | 290551 |
| 479.73 | 1.6 | 2.3E-12 | 0 | 0 | 0 | 0 | 241452 |
| 1361.56 | 3.2 | 2.6E-12 | 0 | 0 | 0 | 0 | 318507 |
| 471.18 | 1.8 | 6.5E-12 | 0 | 94936 | 0 | 0 | 0 |
| 262.12 | 1.3 | 1.1E-11 | 89145 | 0 | 0 | 0 | 0 |
| 519.13 | 1.9 | 2.1E-11 | 192625 | 207592 | 247175 | 165182 | 142112 |
| 358.18 | 1.0 | 2.9E-11 | 0 | 0 | 78544 | 0 | 0 |
| 330.13 | 2.0 | 5.2E-11 | 48619 | 47760 | 521489 | 301795 | 0 |
| 503.15 | 1.9 | 7.0E-11 | 534560 | 588332 | 635199 | 406905 | 312217 |
| 453.16 | 2.0 | 9.3E-11 | 0 | 0 | 395727 | 254423 | 0 |
| 481.16 | 1.9 | 2.7E-10 | 325582 | 289713 | 324147 | 167661 | 183475 |
| 472.72 | 1.4 | 3.7E-10 | 0 | 0 | 0 | 0 | 248165 |
| 1347.55 | 2.9 | 3.9E-10 | 0 | 0 | 0 | 0 | 157561 |
| 364.15 | 1.9 | 4.0E-10 | 0 | 0 | 83880 | 0 | 0 |
| 469.16 | 1.9 | 1.1E-09 | 177134 | 185654 | 197363 | 114583 | 95596 |
| 333.07 | 2.6 | 2.4E-09 | 0 | 0 | 0 | 0 | 95864 |
| 531.16 | 2.3 | 4.3E-09 | 113319 | 106446 | 107331 | 0 | 0 |
| 259.09 | 1.9 | 4.9E-09 | 345602 | 319680 | 138403 | 63290 | 0 |
| 372.15 | 1.3 | 1.1E-08 | 0 | 0 | 0 | 64504 | 240572 |
| 330.13 | 2.2 | 1.2E-08 | 0 | 0 | 102379 | 0 | 0 |
| 480.23 | 1.6 | 1.3E-08 | 0 | 0 | 0 | 0 | 168832 |
| 352.12 | 1.9 | 3.8E-08 | 145120 | 162087 | 165483 | 97353 | 41173 |
| 305.08 | 2.2 | 1.1E-07 | 171183 | 173029 | 226425 | 127493 | 104898 |
| 676.25 | 1.1 | 1.8E-07 | 0 | 0 | 0 | 0 | 104863 |
| 564.25 | 1.3 | 3.4E-07 | 0 | 0 | 134702 | 21775 | 0 |
| 447.18 | 1.8 | 8.9E-06 | 152230 | 131107 | 145071 | 69117 | 0 |
| 479.21 | 1.5 | 1.8E-05 | 0 | 23672 | 0 | 0 | 104027 |
| 336.13 | 1.9 | 2.3E-05 | 131275 | 63697 | 0 | 0 | 0 |
| 283.13 | 1.4 | 2.4E-05 | 0 | 0 | 101784 | 0 | 0 |
| 616.18 | 3.4 | 3.0E-05 | 113118 | 21688 | 21543 | 0 | 100116 |
| 585.35 | 1.4 | 4.2E-05 | 81884 | 113695 | 32255 | 0 | 0 |
| 416.11 | 2.2 | 4.3E-05 | 114862 | 113974 | 129047 | 22101 | 27435 |
| 515.18 | 2.3 | 5.3E-05 | 168673 | 168939 | 193283 | 114637 | 53574 |
| 576.26 | 1.9 | 8.3E-05 | 0 | 0 | 103787 | 27383 | 0 |
| 360.23 | 1.4 | 8.8E-05 | 0 | 45420 | 128057 | 0 | 131936 |
| 619.17 | 3.3 | 9.1E-05 | 0 | 0 | 0 | 27907 | 99799 |

**Table S5** DEPTQ and $^{1}H$ NMR assignments for **12** in $CD_3OD$.

| **Residue** *Including atom number* | **δC (ppm)** *Including no. of attached protons* | **δH (ppm), mult. (*J*, in Hz)** |
|---|---|---|
| **Acyl** | | |
| 1 | 22.6, $CH_3$ | 1.99 (s) |
| 2 | 172.8, C | |
| **Ala5** | | |
| 3 | 56.7, CH | 4.71 (q, 7.2) |
| 4 | 21.1, $CH_3$ | 1.39 (d, 7.0) |
| 5 | 206.0, C | |
| NH | | ND[a] |
| **Ile6** | | |
| 6 | 68.9, CH | 5.16 (d, 8.9) |
| 7 | 41.0, CH | 2.08 (m) |
| 8 | 15.6, $CH_3$ | 0.95 (m) |
| 9 | 26.2, $CH_2$ | 1.25, 1.64 (br m) |
| 10 | 11.3, $CH_3$ | 0.91[b] (m) |
| 11 | 203.7, C | |
| NH | | ND[a] |
| **Ala7** | | |
| 12 | 55.8, CH | 5.05 (q, 7.2) |
| 13 | 17.5, $CH_3$ | 1.54 (d, 7.1) |
| 14 | 172.4, C | |
| NH | | ND[a] |
| **Dhb8** | | |
| 15 | 129.1[c] | |
| 16 | 136.4, CH | 6.83 (q, 7.2) |
| 17 | 14.4, $CH_3$ | 1.77 (d, 7.3) |
| 18 | 167.7[c] | |
| NH | | ND[a] |
| OH | | ND[a] |

a. The NH and OH protons are not visible due to deuterium exchange with the $CD_3OD$.
b. The signal for this hydrogen is masked by a contaminant compound; COSY and HMBC correlations are visible (Figures S13 and S15).
c. Due to low signal these carbons are below the level of noise in the DEPTQ spectrum (Figure S12), however their HMBC correlations are visible (Figure S15).

**Table S6** Yeast assemblies of pCAPtsa variants and their constituent parts.

| Construct | Backbone | Enzymes used | Parts |
|---|---|---|---|
| pCAPtsaTsaCoreTaa | pCAPtsa | AflII and SrfI | A, B, K |
| pCAPtsaS1T | pCAPtsa | AflII and SrfI | A, C, K |
| pCAPtsaSAV | pCAPtsa | AflII and SrfI | A, D, K |
| pCAPtsaM3I | pCAPtsa | AflII and SrfI | A, E, K |
| pCAPtsaA5F | pCAPtsa | AflII and SrfI | A, F, K |
| pCAPtsaT8S | pCAPtsa | AflII and SrfI | A, G, K |
| pCAPtsaY11V | pCAPtsa | AflII and SrfI | A, H, K |
| pCAPtsaH12A | pCAPtsa | AflII and SrfI | A, I, K |
| pCAPtsaH12W | pCAPtsa | AflII and SrfI | A, J, K |

**Table S7** Parts used in cluster assemblies.

| Part | Primer pairs / oligonucleotides | Size of PCR products (bp) or oligonucleotides (bases) | Contains | Left linker (towards *btmA* or *tsaA*) | Right linker (towards *btmM* or *tsaL*) |
|---|---|---|---|---|---|
| A | SPTsaAassembly<br>ASPTsaAassembly | 473 bp | *tsaA* | Plasmid backbone | No linker |
| B | TsaCoreAlba | 119 bases | Mutated *tsaA* core | *tsaA* | *tsaC* |
| C | S1T | 119 bases | Mutated *tsaA* core | *tsaA* | *tsaC* |
| D | SAV | 120 bases | Mutated *tsaA* core | *tsaA* | *tsaC* |
| E | M3I | 119 bases | Mutated *tsaA* core | *tsaA* | *tsaC* |
| F | A5F | 119 bases | Mutated *tsaA* core | *tsaA* | *tsaC* |
| G | T8S | 119 bases | Mutated *tsaA* core | *tsaA* | *tsaC* |
| H | Y11V | 119 bases | Mutated *tsaA* core | *tsaA* | *tsaC* |
| I | H12A | 119 bases | Mutated *tsaA* core | *tsaA* | *tsaC* |
| J | H12W | 119 bases | Mutated *tsaA* core | *tsaA* | *tsaC* |
| K | SPTsaCassembly<br>ASPTsaCassembly | 1081 bp | *tsaC* | No linker | *tsaD* |

### SUPPLEMENTARY FIGURES

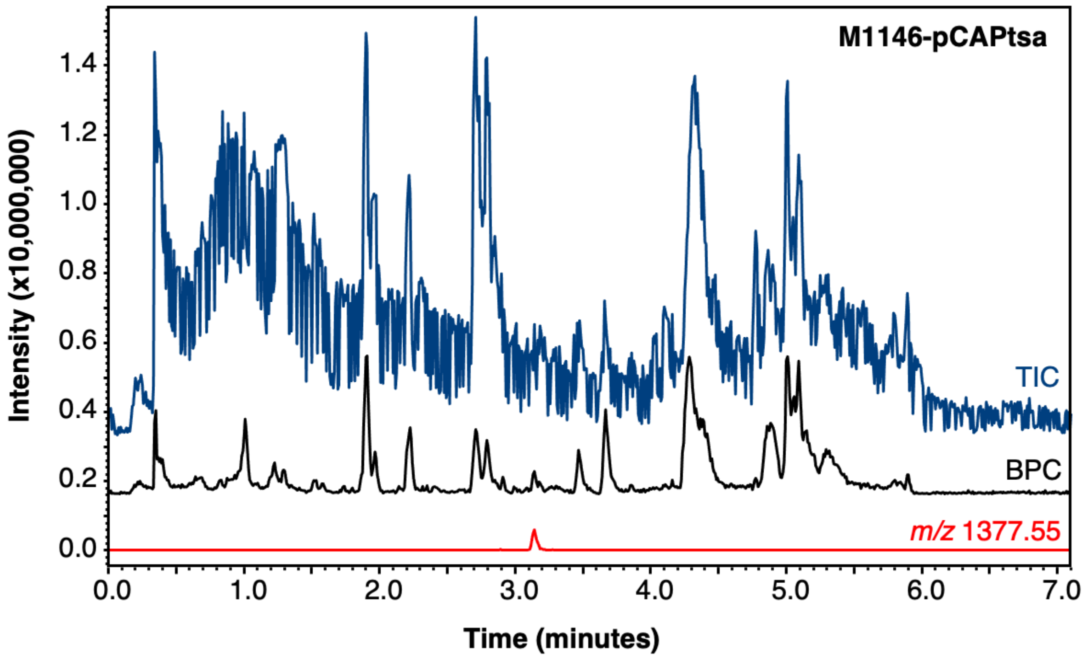


**Figure S1** LC-MS detection of thiostreptamide S4 (**1**, *m/z* 1377.55) produced by *S. coelicolor* M1146-pCAPtsa in comparison to the total ion chromatogram (TIC) and the base peak chromatogram (BPC).

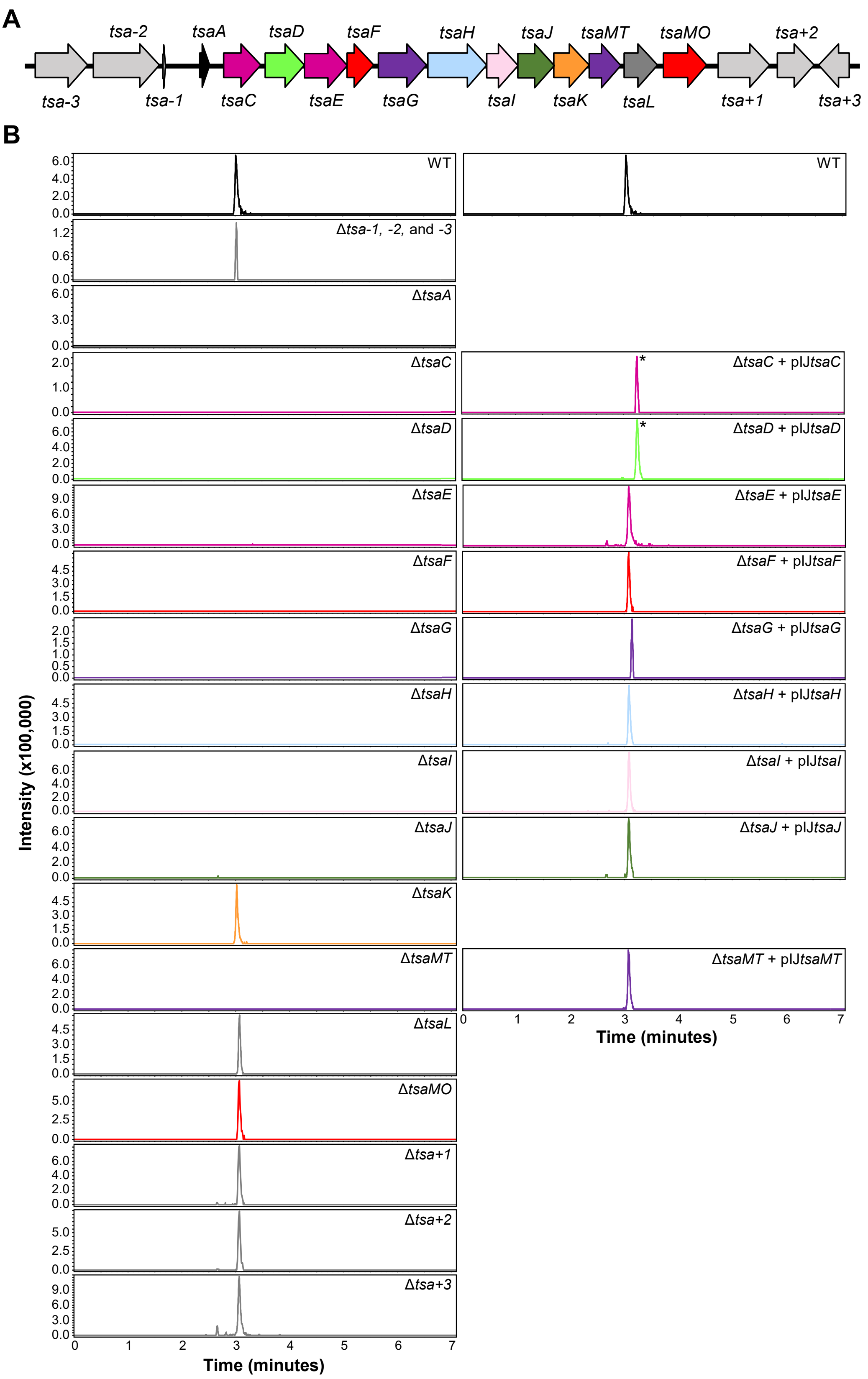


**Figure S2** A. The genes captured on pCAPtsa. The genes in the predicted BGC are colour-coded as in the main paper. B. LC-MS analysis of each gene deletion cluster and their successful complementations in *S. coelicolor* M1146. An extracted ion chromatogram (EIC) of thiostreptamide S4 (**1**; *m/z* 1377.55) is shown for each mutant. The wild type (WT) EIC is duplicated as a reference chromatogram for each set of peaks. An asterisk highlights the Δ*tsaC* + pIJ*tsaC* and Δ*tsaD* + pIJ*tsaD* peaks, whose retention times are shifted due to the addition of a guard to the column.

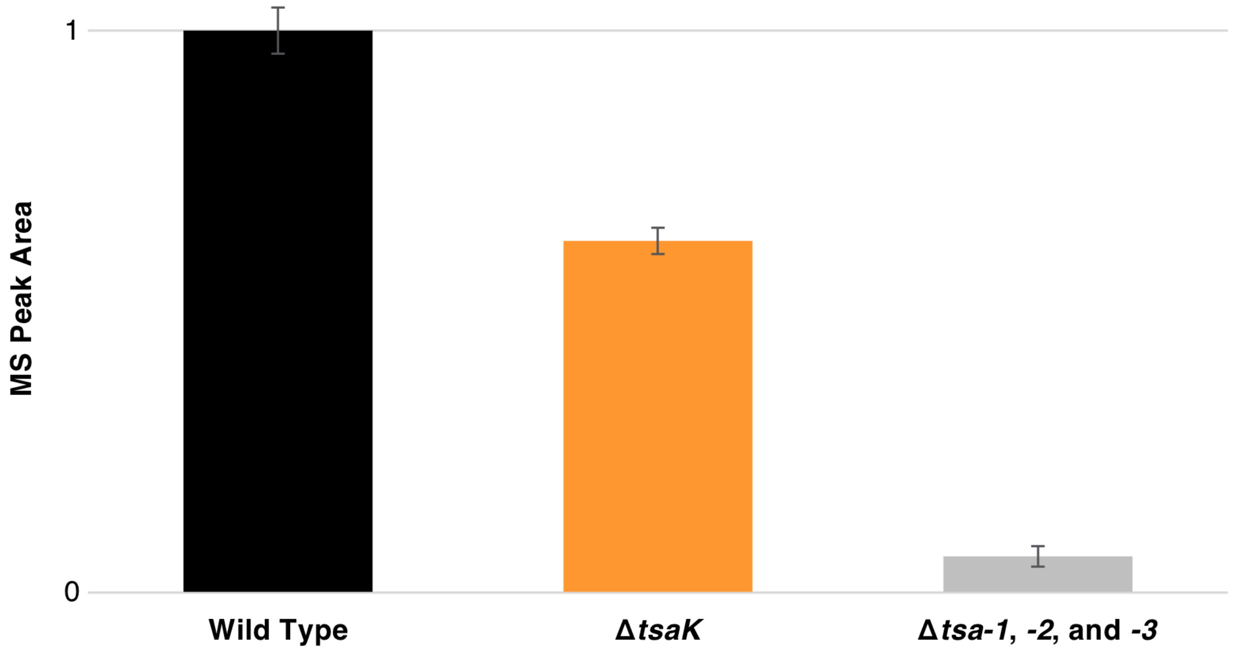


**Figure S3** Mutants that produced thiostreptamide S4 (**1**; *m/z* 1377.55), but at lower levels than the wild type BGC. MS peak areas are shown for *S. coelicolor* M1146 containing either the wild type *tsa* BGC, the Δ*tsaK* BGC, or the Δ*tsa-1*, *-2*, and *-3* BGC. Values represent the average of three biological replicas and the error bars represent the standard error.

**A**

```
tsaC       ArgAla***
tsaD            MetSerArgTyrAspAlaValPheAlaSerMetAlaGluAspIleGluValLeuAsp
      1    CGCGCATGAGCAGATACGACGCAGTGTTCGCCAGCATGGCGGAGGACATAGAGGTCCTGGAC

tsaD       GlnAlaThrPheArgHisArgGluTrpGlyAspLeuSerProAlaAlaGlyValGluThr
     63    CAGGCAACTTTCCGTCACCGGGAGTGGGGTGACCTGTCCCCTGCTGCTGGGGTGGAGACA
```

**B**

```
tsaF       GlyGln***
tsaG            ValIleProAspMetThrLysLeuSerProGlnThrValLeuTyrArgAlaProSer
      1    GGACAGTGATCCCAGACATGACCAAGCTCTCACCACAGACGGTTCTCTACCGGGCGCCGTCC

tsaG       PheValAlaGluValAspThrSerAsnGluValLysIleHisPheGluGlyArgValLeuLys
     63    TTTGTCGCCGAGGTCGACACCAGTAACGAGGTGAAGATCCACTTCGAAGGCCGGGTGCTCAAG
```

**Figure S4** Identification of functional start codons in *tsaD* and *tsaG* (see Table S3). A. Region of the *tsa* BGC containing three potential start codons (red) for *tsaD* that were tested by complementation of Δ*tsaD*. The first start codon (bold), coupled to the *tsaC* stop codon, is the only one that successfully complemented the *tsaD* deletion. The third highlighted start codon is the originally annotated start codon. B. Region of the *tsa* BGC containing four potential start codons (red) for *tsaG* that were tested by complementation of Δ*tsaG*. The first start codon (bold), coupled to the *tsaF* stop codon, and the second start codon (bold) are the only start codons that successfully complemented the *tsaG* deletion.

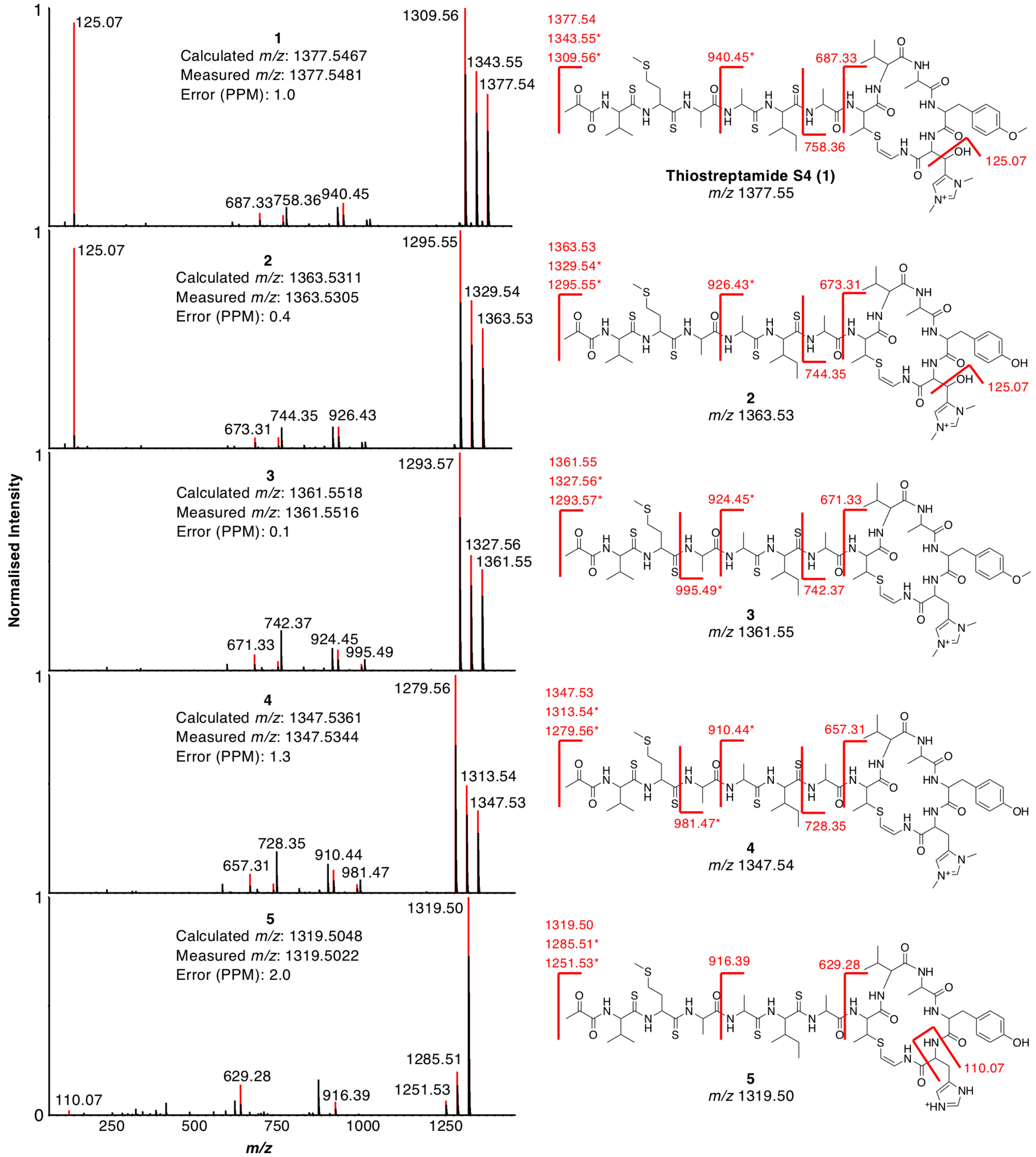


**Figure S5** Accurate mass and MS/MS fragmentation data for compounds **1** - **5**. Annotated fragments are coloured red in the MS/MS spectra. Fragments marked with an asterisk show a loss of 33.99, characteristic of the loss of $SH_2$ from thioamide bonds in MS/MS fragmentation (see Figure S6). Exact mass measurements are shown within the mass spectra.

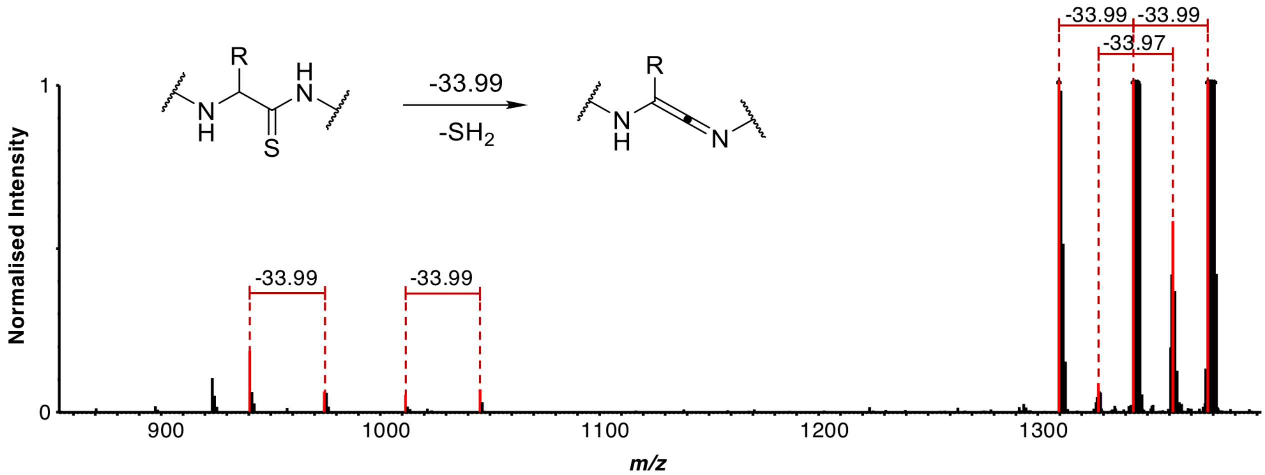


**Figure S6** Selected MS/MS fragmentation of thiostreptamide S4 (**1**) showing losses of 33.99 Da, which is characteristic of thioamide bond fragmentation. The mass spectrum is zoomed for clarity and a proposed reaction scheme is shown.

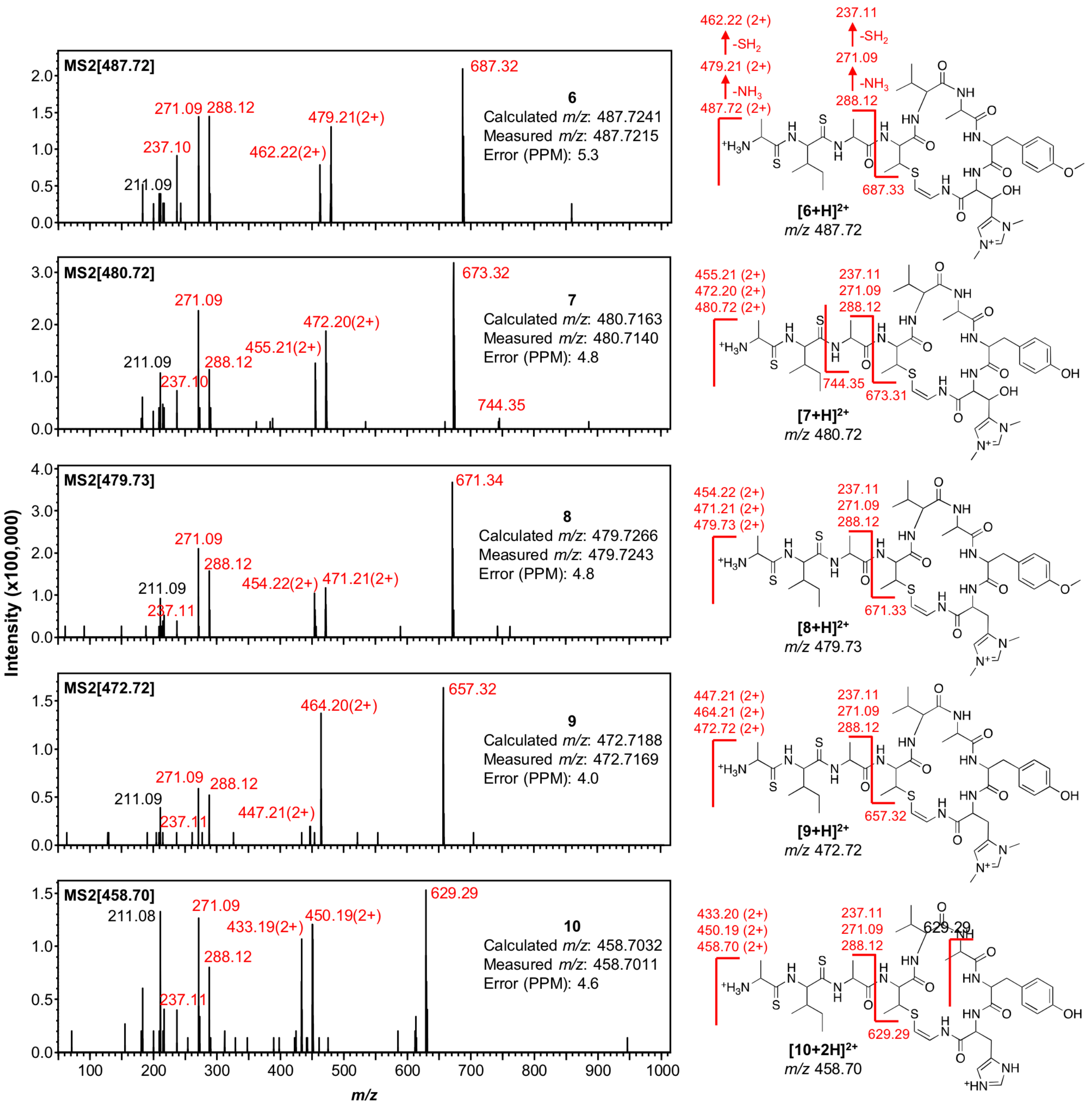


**Figure S7** MS/MS fragmentation data for compounds **6** - **10** alongside predicted molecule structures. Exact *m/z* measurements for doubly charged molecules are shown within the mass spectra; note that compounds **6** - **9** are protonated once to become doubly charged. Characteristic macrocycle fragments are highlighted, as are other common fragments (losses of $NH_3$ and $SH_2$ are shown for **6**). Calculated fragment masses shown with the structures.

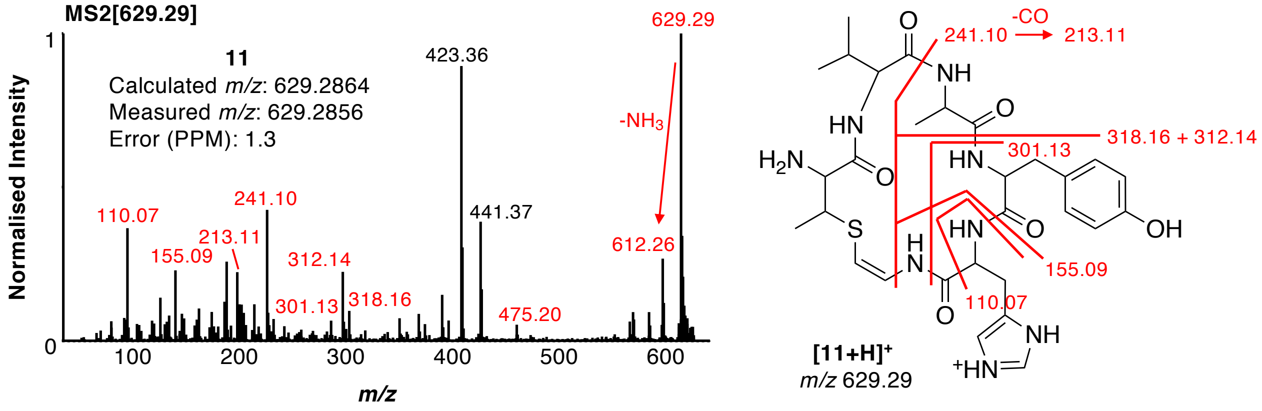


**Figure S8** MS/MS fragmentation data for compound **11** alongside predicted molecule structure. An exact *m/z* measurement is shown within the mass spectrum. Fragments are highlighted, where calculated fragment masses are shown with the structure.

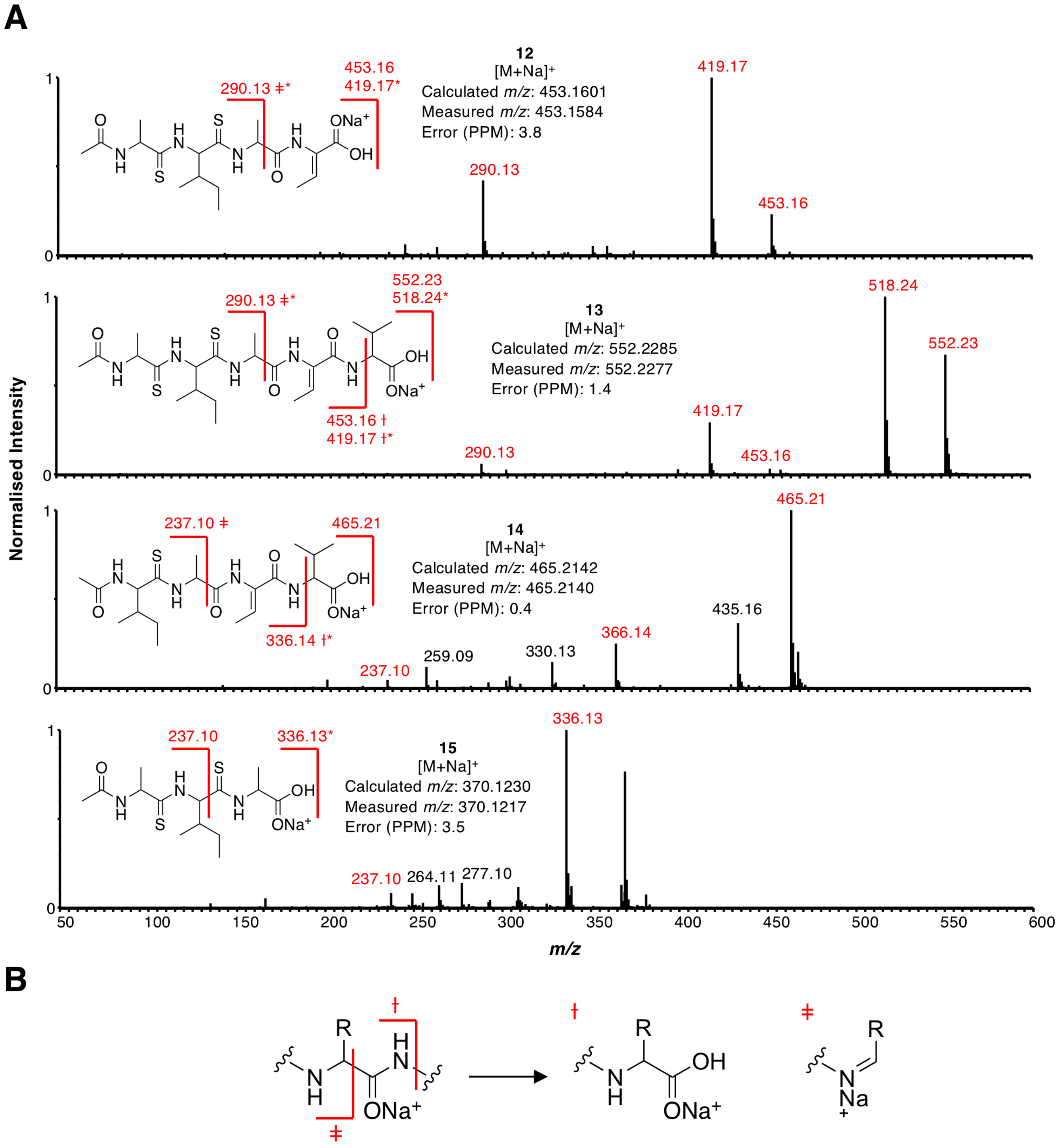


**Figure S9** A. MS/MS spectra showing fragmentation of compounds **12** - **15** alongside predicted structures as sodium adducts. The *m/z* of the parent molecule is shown in the top right of each chromatogram. Fragments corresponding to unusual sodiated peptide fragments (panel B) are highlighted by symbols, and fragments marked with an asterisk show a loss of 33.99, characteristic of the loss of $SH_2$ from thioamide bonds. B. The unusual position of bond breakage common in the fragmentation of the sodium adduct of peptides[23,24].

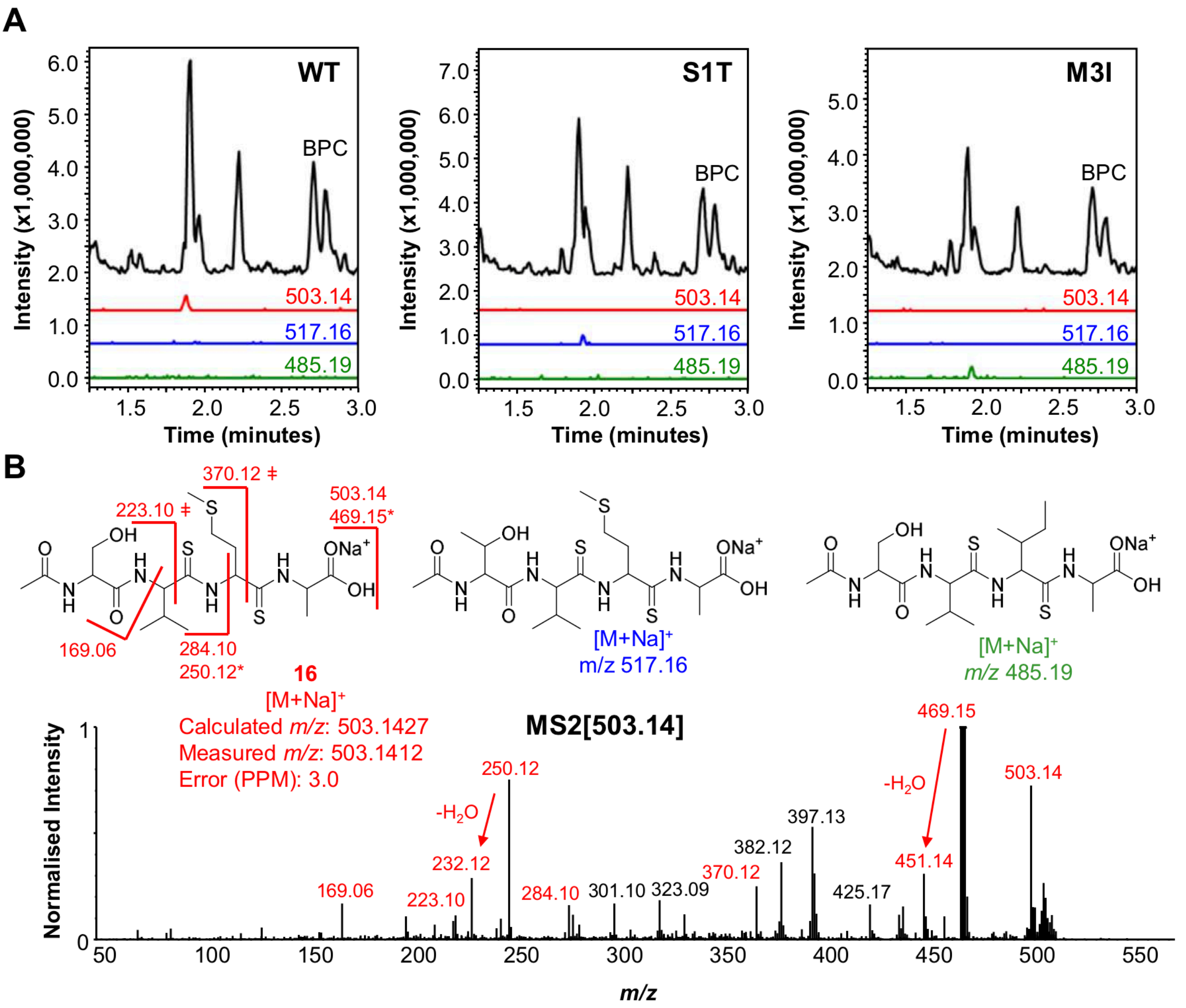


**Figure S10** A. LC-MS analysis of *S. coelicolor* M1146 expressing pCAPtsa (wild type, WT), pCAPtsa S1T and pCAPtsaM3I. The BPC and EICs of **16** and two related metabolites (*m/z* 503.14, 517.16 and 485.19, respectively) are shown. The EICs are magnified 2x compared to the BPCs for clarity. B. Predicted structures of **16** and the two related metabolites labelled with their *m/z*. Fragments of **16** observed in the MS/MS spectrum are shown on the molecule and highlighted in the spectrum in red. Fragments marked with an asterisk show a loss of 33.99 (loss of $SH_2$) and symbols are used to highlight the unusual sodiated peptide fragmentation described in Figure S9.

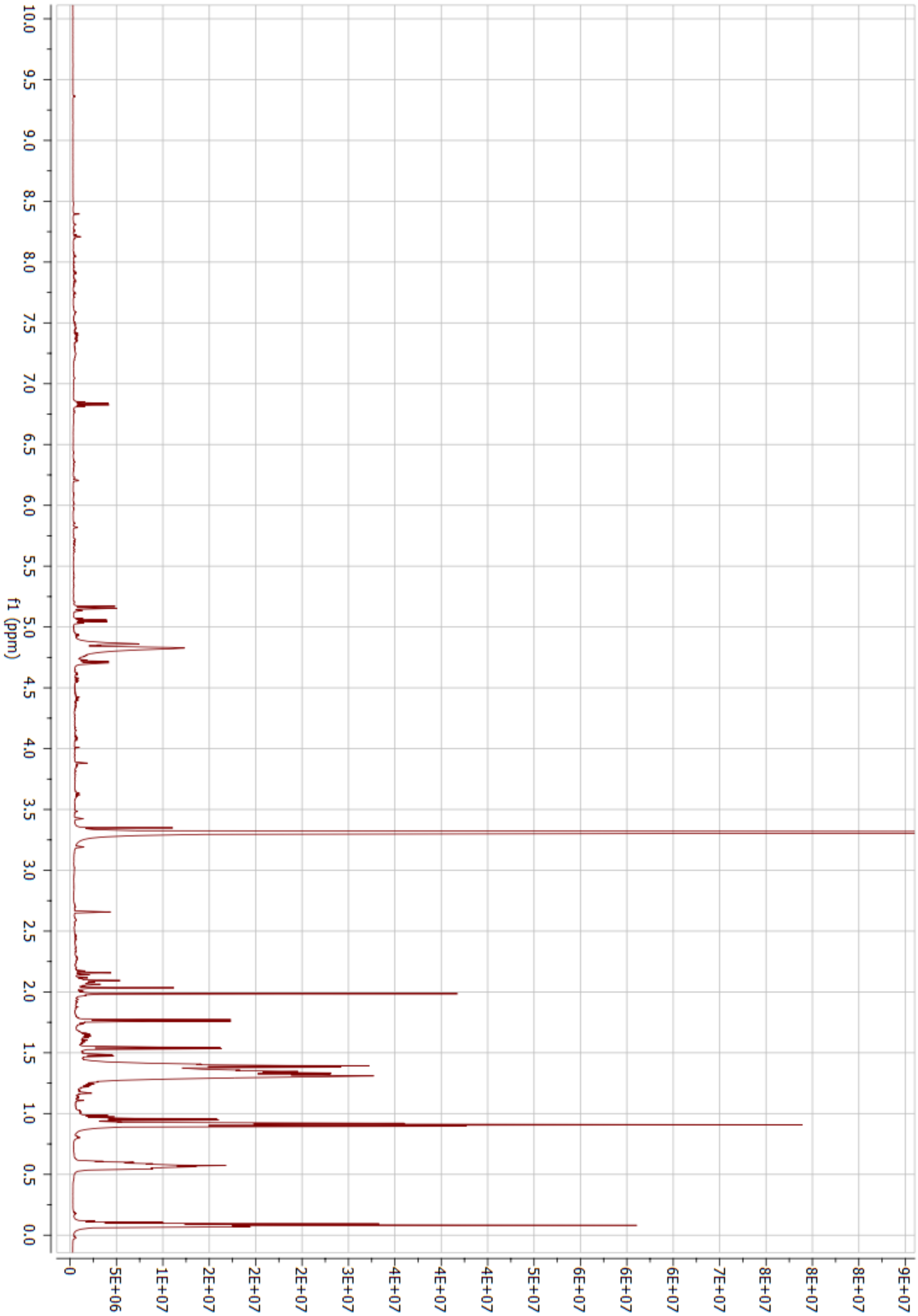


**Figure S11** [1]H NMR spectrum for molecule **12** (600 MHz, $CD_3OD$, 298 K).

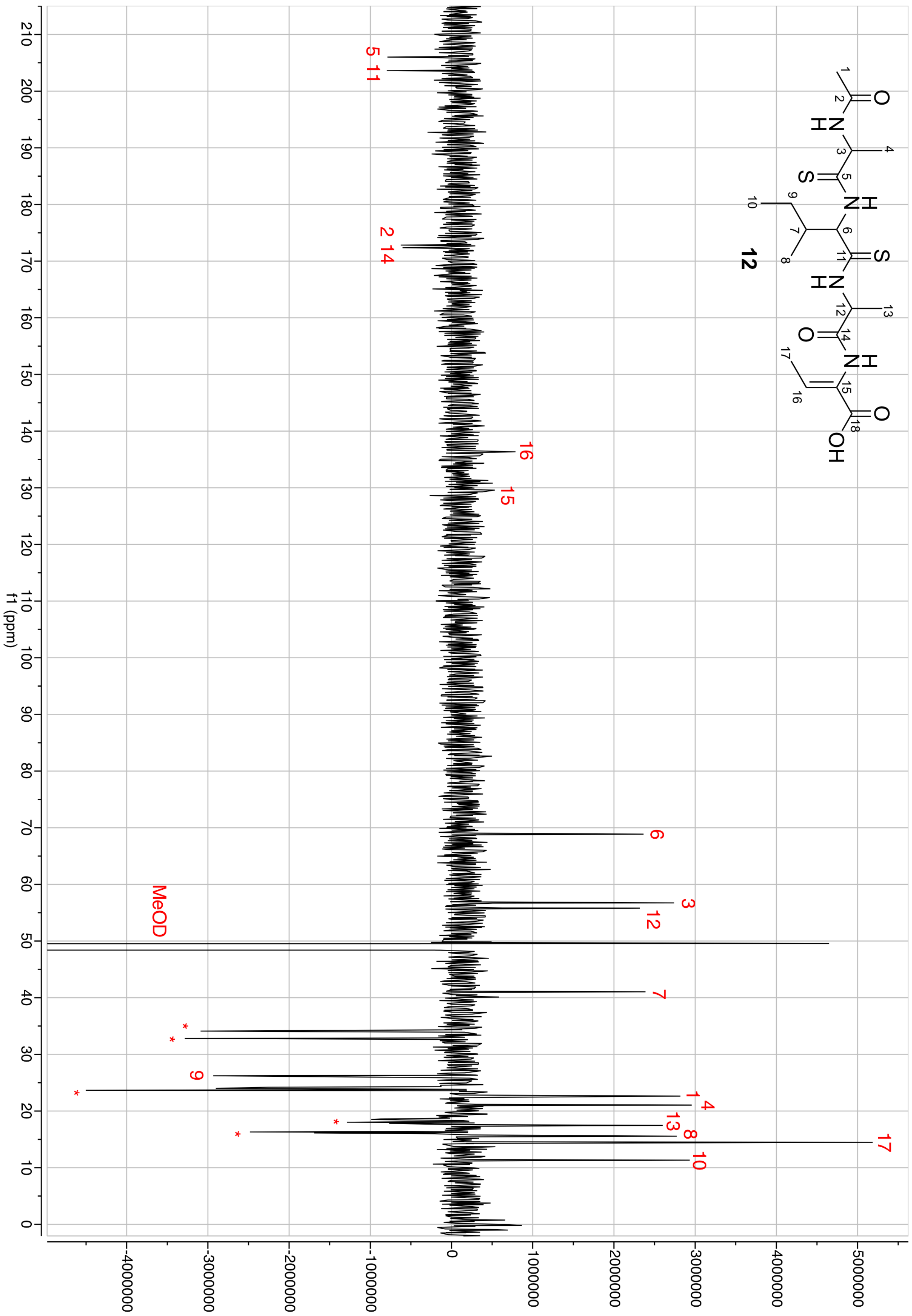


**Figure S12** DEPTQ NMR spectrum for molecule **12** (150 MHz, $CD_3OD$, 298 K). Peaks are labelled. * Impurity peaks.

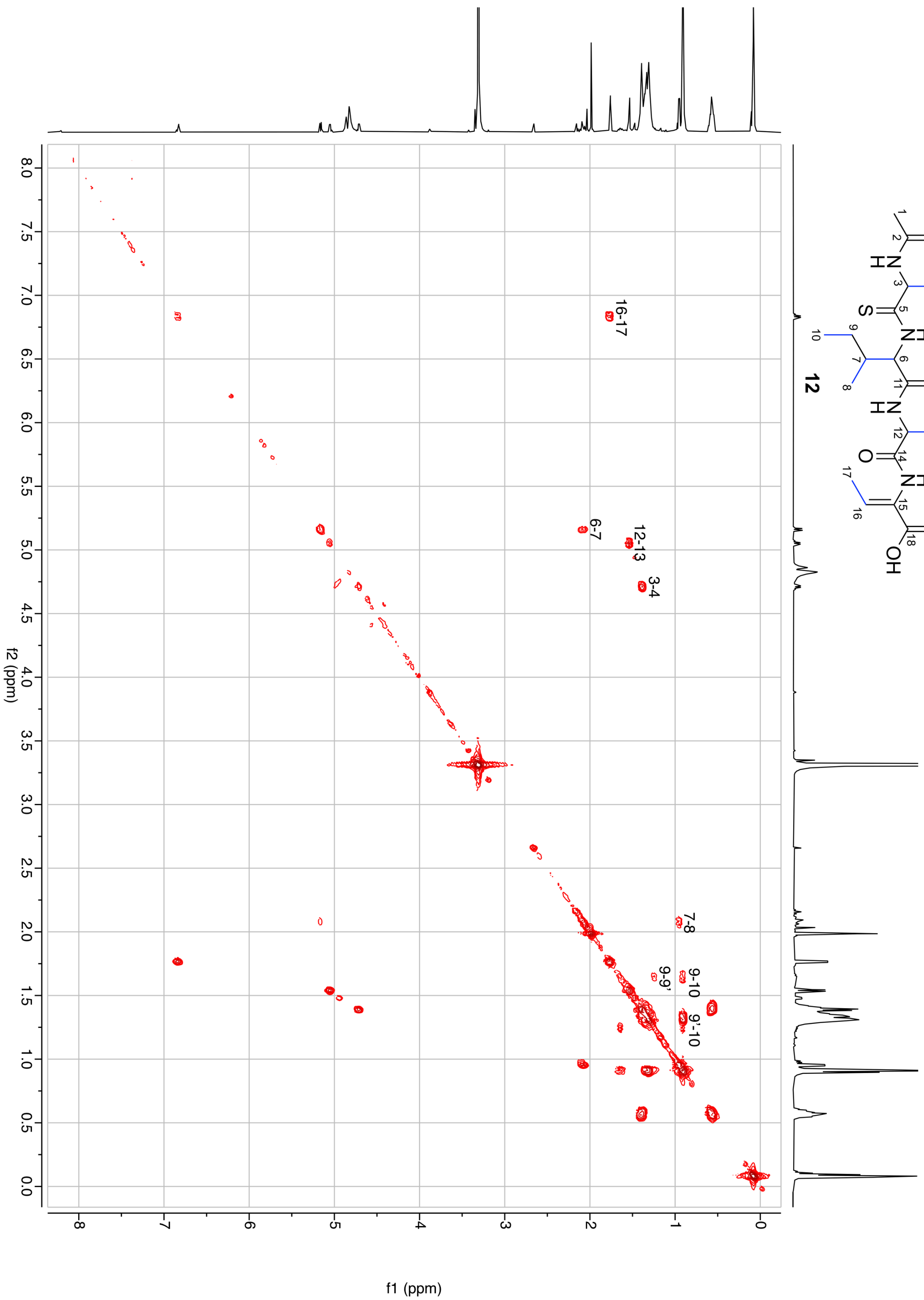


**Figure S13** 2D COSY NMR spectrum for molecule **12** ($CD_3OD$, 298 K). Correlations are annotated. 9 and 9' relate to diastereotopic protons on carbon 9. Visible correlations are shown as blue bonds in **12**.

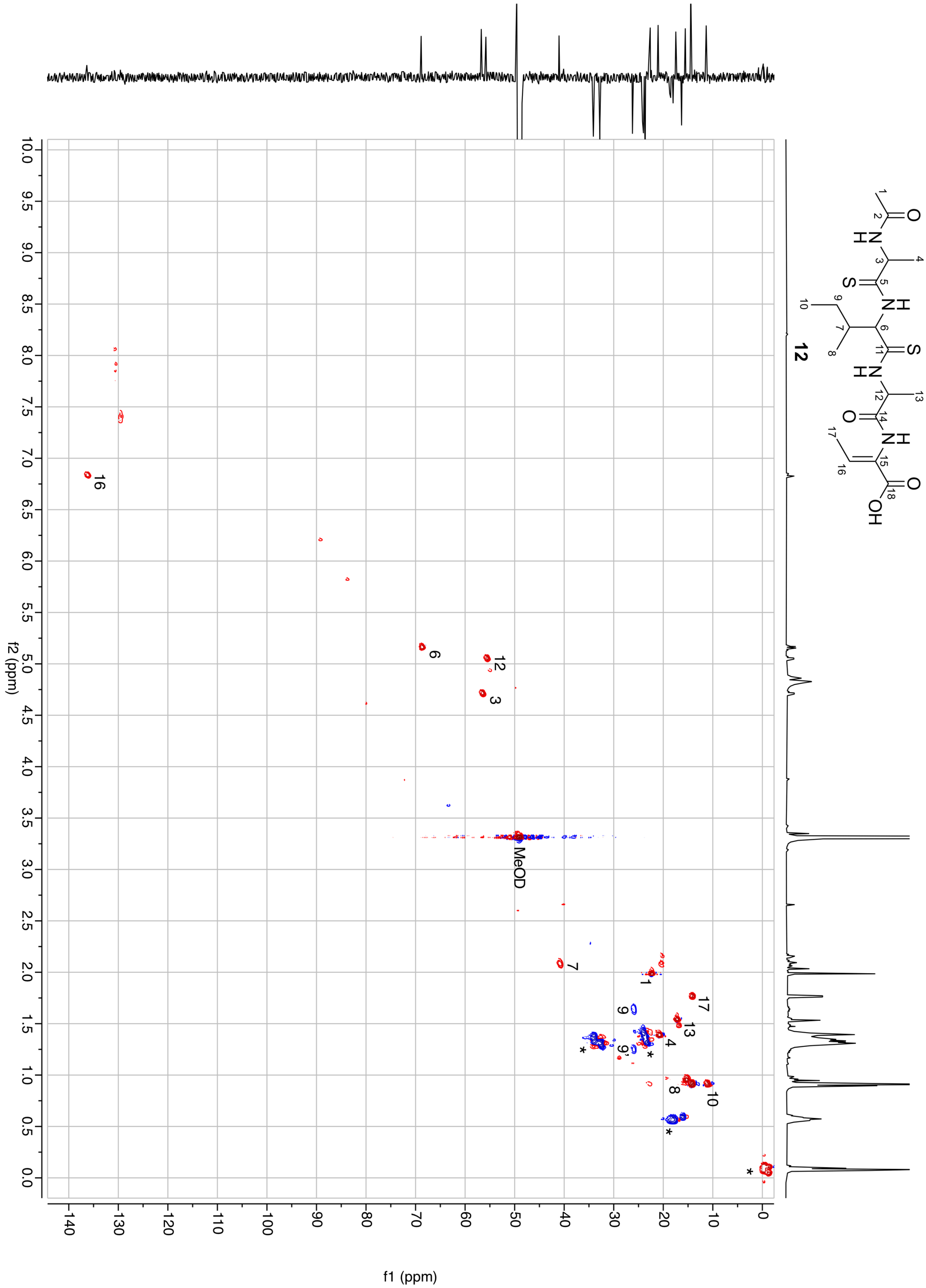


**Figure S14** 2D HSQCed NMR spectrum for molecule **12** ($CD_3OD$, 298 K). * Impurity peaks.

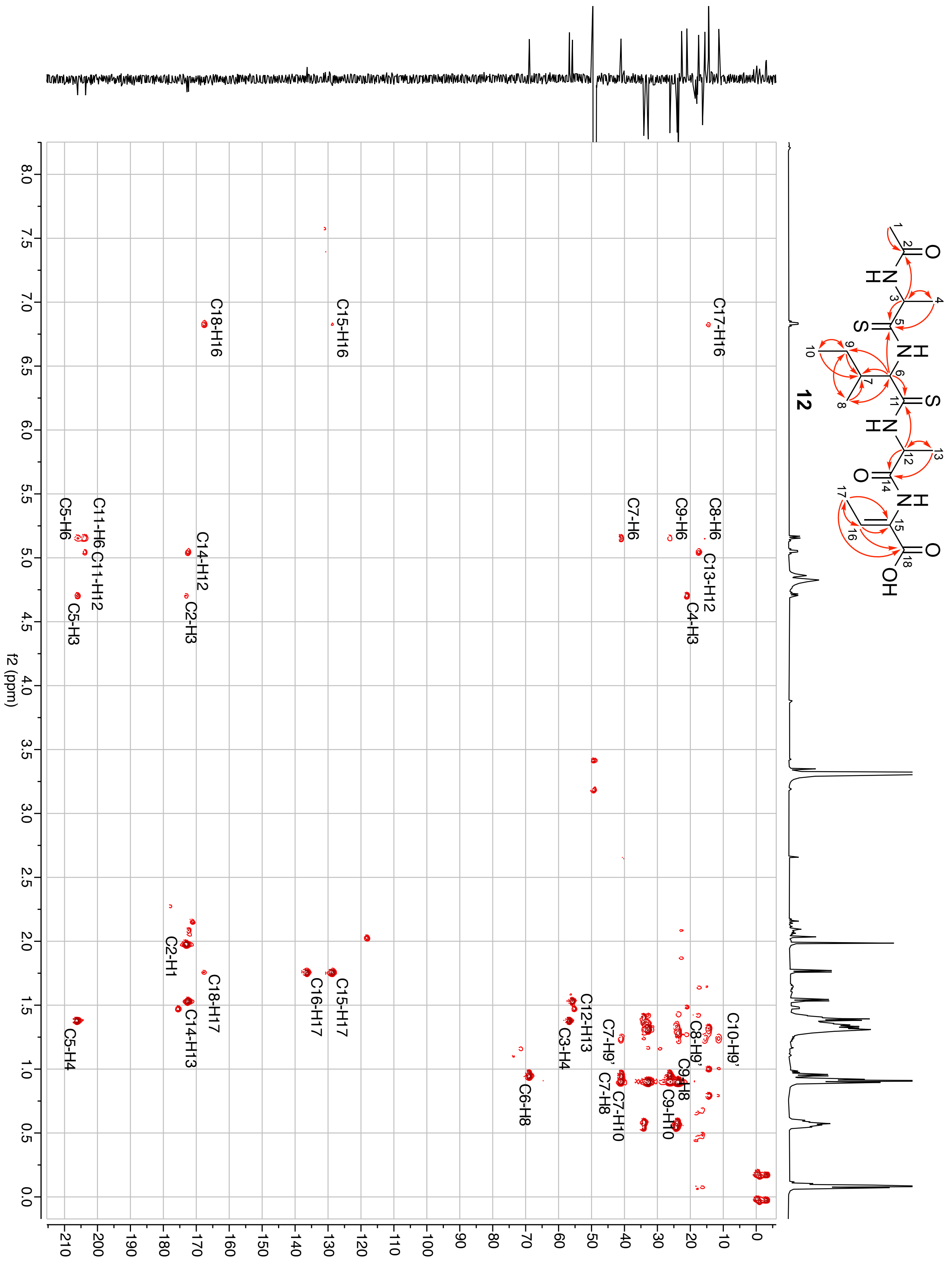


**Figure S15** 2D HMBC NMR spectrum for molecule **12** ($CD_3OD$, 298 K). Correlations are annotated. Visible correlations are shown as red arrows in **12**, where arrows represent H to C direction of correlation; double headed arrow indicates that correlations in both directions were detected.

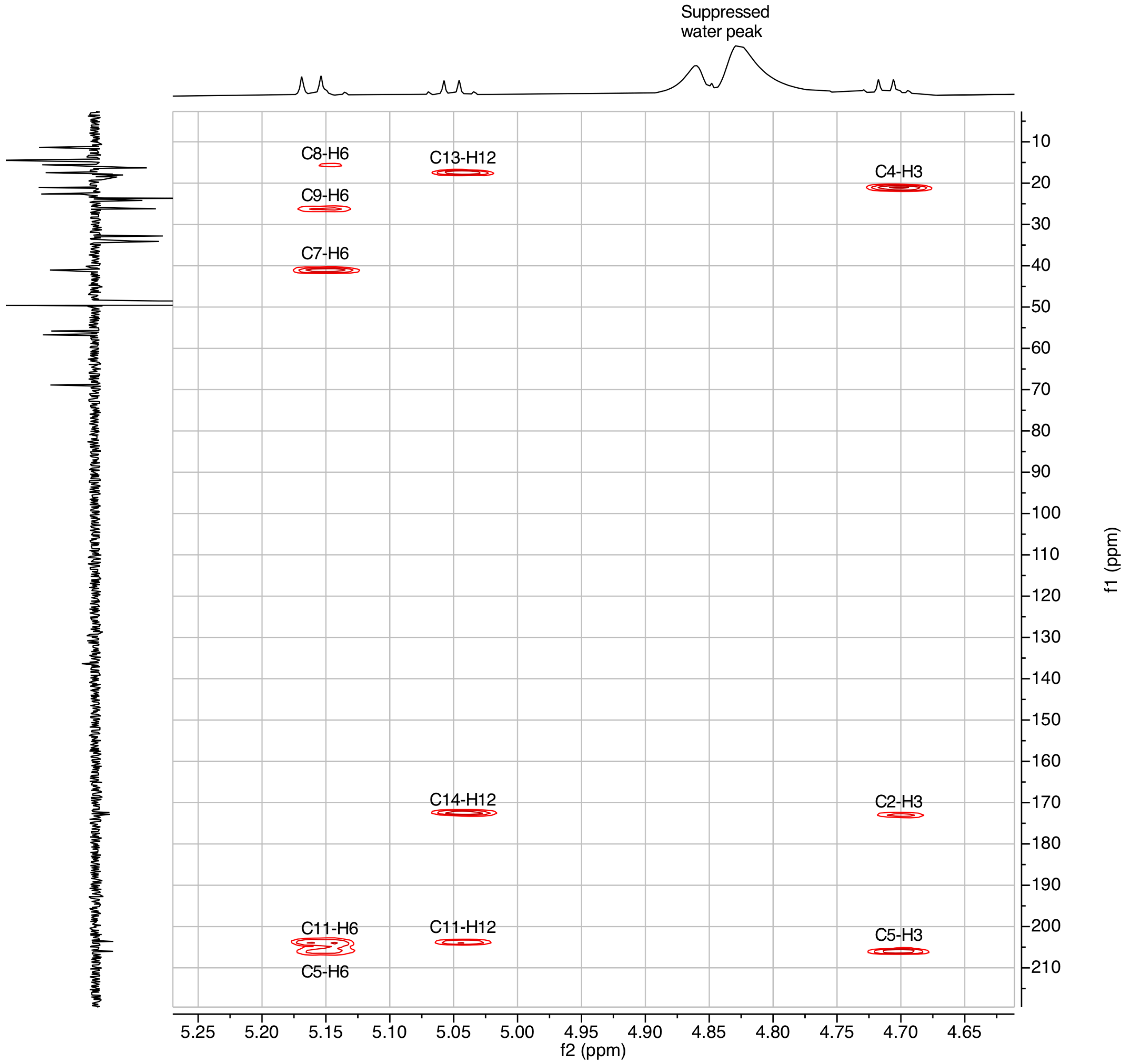


**Figure S16** Expanded detail of 2D HMBC NMR spectrum for molecule **12** ($CD_3OD$, 298 K) showing key thioamide correlations.

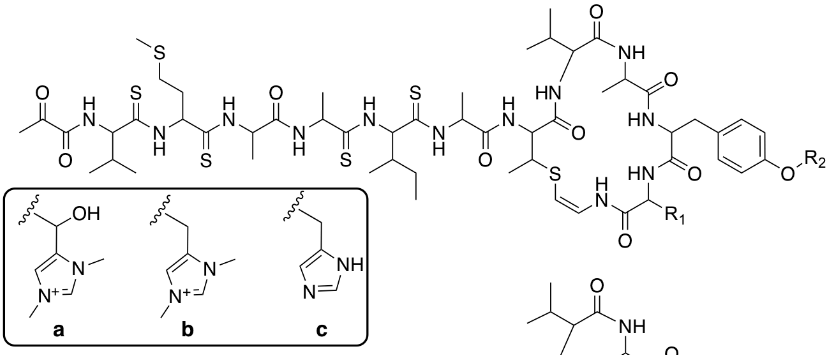


| Molecule | | Major detected version | *m/z* |
|---|---|---|---|
| **1** | $R_1$ = **a**, $R_2$ = $CH_3$ | **M+** | 1377.55 |
| **2** | $R_1$ = **a**, $R_2$ = H | **M+** | 1363.53 |
| **3** | $R_1$ = **b**, $R_2$ = $CH_3$ | **M+** | 1361.55 |
| **4** | $R_1$ = **b**, $R_2$ = H | **M+** | 1347.54 |
| **5** | $R_1$ = **c**, $R_2$ = H | **[M+H]+** | 1319.50 |

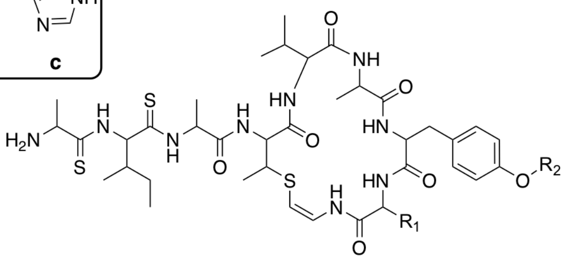


| Molecule | | Major detected version | *m/z* |
|---|---|---|---|
| **6** | $R_1$ = **a**, $R_2$ = $CH_3$ | **[M+H]2+** | 487.72* |
| **7** | $R_1$ = **a**, $R_2$ = H | **[M+H]2+** | 480.72* |
| **8** | $R_1$ = **b**, $R_2$ = $CH_3$ | **[M+H]2+** | 479.73* |
| **9** | $R_1$ = **b**, $R_2$ = H | **[M+H]2+** | 472.72* |
| **10** | $R_1$ = **c**, $R_2$ = H | **[M+2H]2+** | 458.70 |

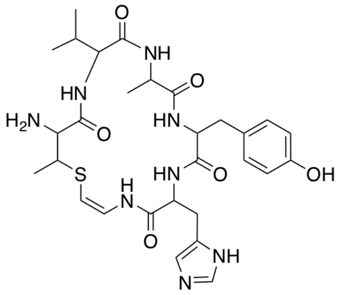


**11** **[M+H]+** 629.29

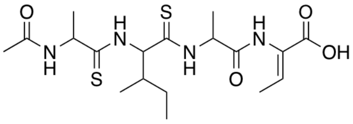

**12** **[M+Na]+** 453.16

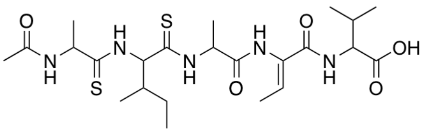

**13** **[M+Na]+** 552.23

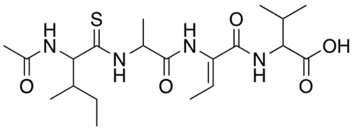

**14** **[M+Na]+** 465.21

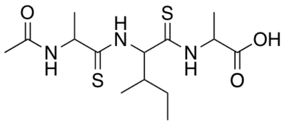

**15** **[M+Na]+** 370.12

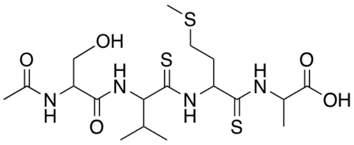

**16** **[M+Na]+** 503.14

**Figure S17** Confirmed structures of compounds **1** and **12**, and structures of compounds **2-11** and **13-16** proposed by detailed MS/MS analysis. *Permanent charge on bis-methylated histidine means that a single protonation generates a doubly charged molecule.

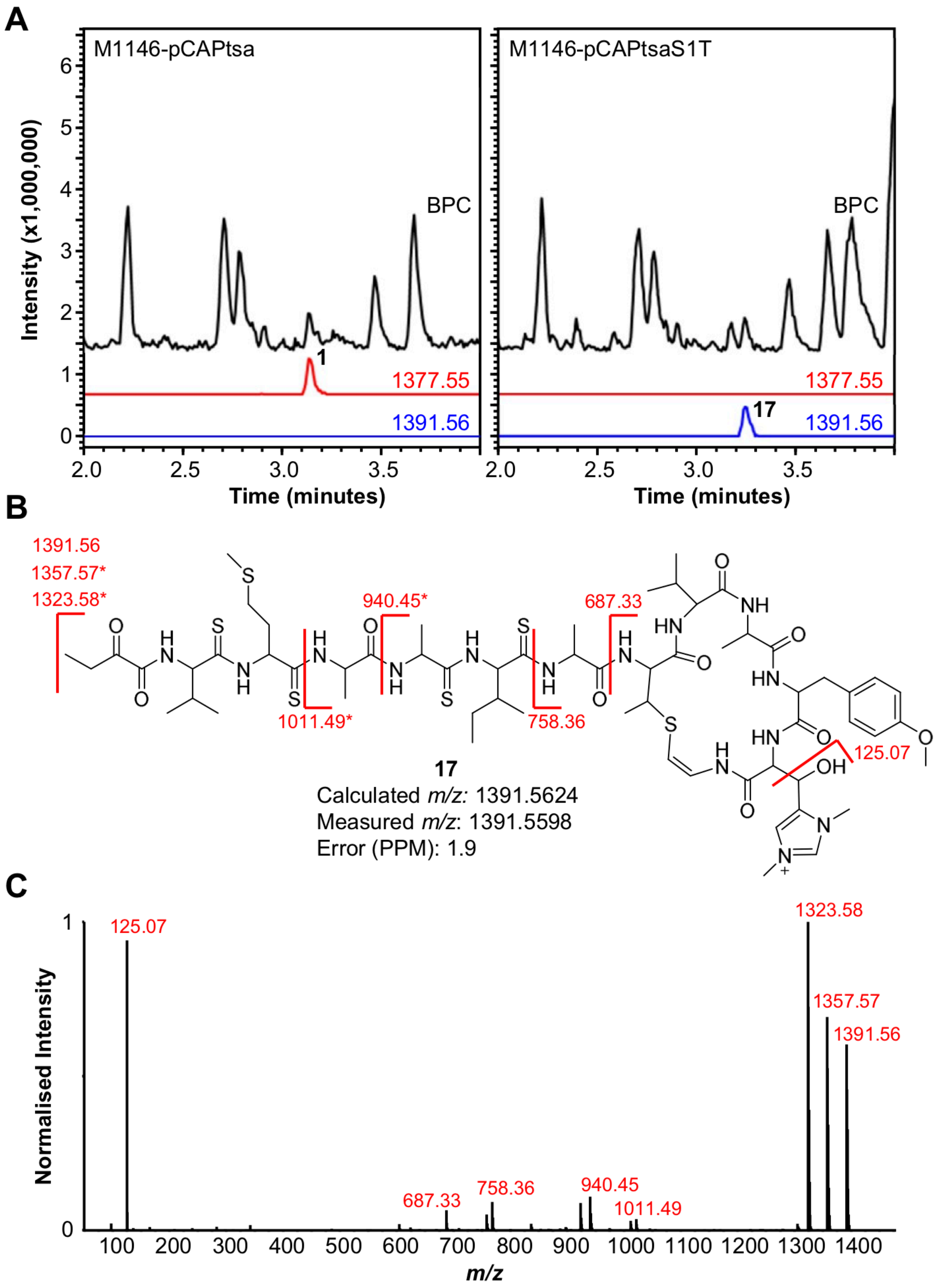


**Figure S18** A. LC-MS analysis of M1146-pCAPtsa and M1146-pCAPtsaS1T. EICs of *m/z* 1377.55 and *m/z* 1391.56 are shown. B. Predicted structure of **17** annotated with fragments observed via MS/MS analysis. Fragments marked with an asterisk show an additional loss of 33.99, characteristic of the loss of $SH_2$ from thioamide bonds. C. MS/MS spectrum of *m/z* 1391.56 from M1146-pCAPtsaS1T.

**A**

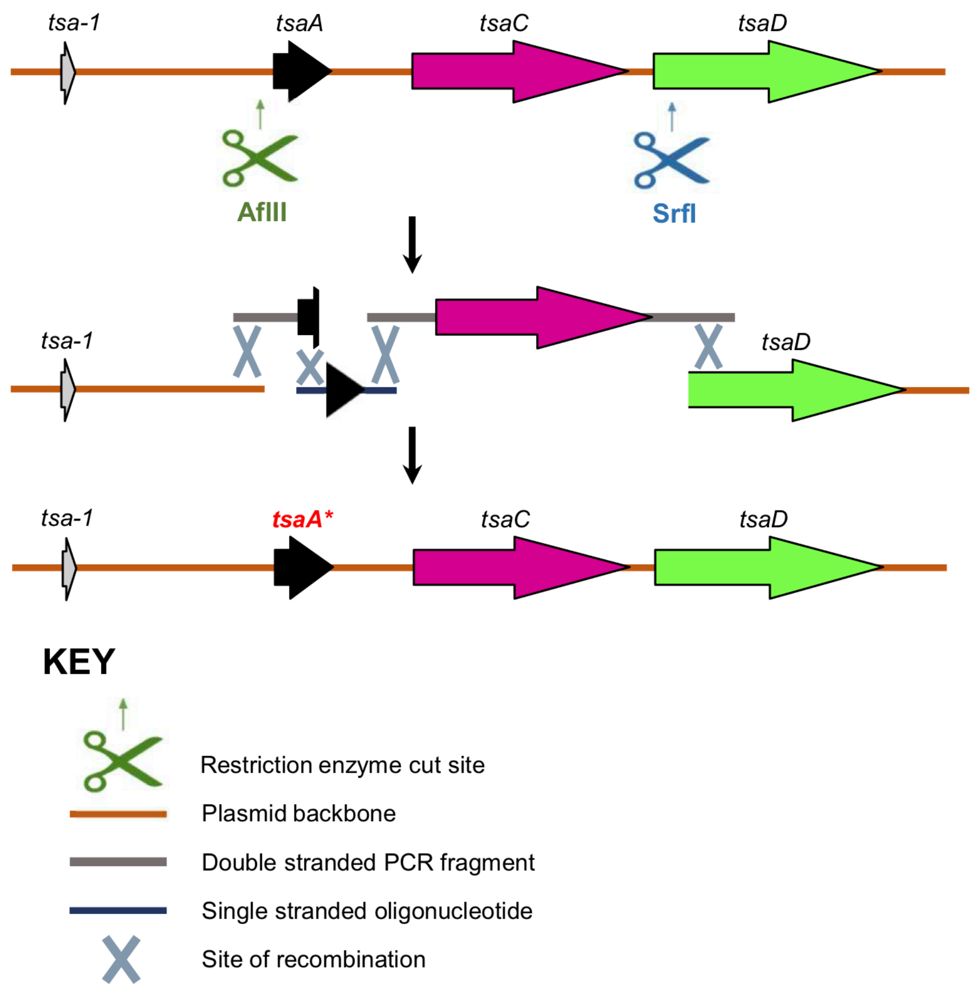


**B**

| | 1 5 10 | |
|---|---|---|
| **Wild type** | SVMAAIATVAYHC | ✓ |
| **S1T** | TVMAAIATVAYHC | ✓ |
| **M3I** | SVIAAIATVAYHC | ✓ |
| **A5F** | SVMAFIATVAYHC | ✗ |
| **T8S** | SVMAAIASVAYHC | ✓ |
| **Y11V** | SVMAAIATVAVHC | ✗ |
| **H12A** | SVMAAIATVAYAC | ✗ |
| **H12W** | SVMAAIATVAYWC | ✗ |
| **TsaCoreTaa** | SVIGFAVTIAVHC | ✗ |
| **SAV** | SAVMAAIATVAYHC | ✗ |

**Figure S19** A. Schematic of yeast assembly-based modification of the *tsaA* core peptide gene in pCAPtsa. B. Mutants generated in this study. A tick indicates that a fully modified thiostreptamide-like compound was detected, while a cross indicates that no fully modified compound was detected.

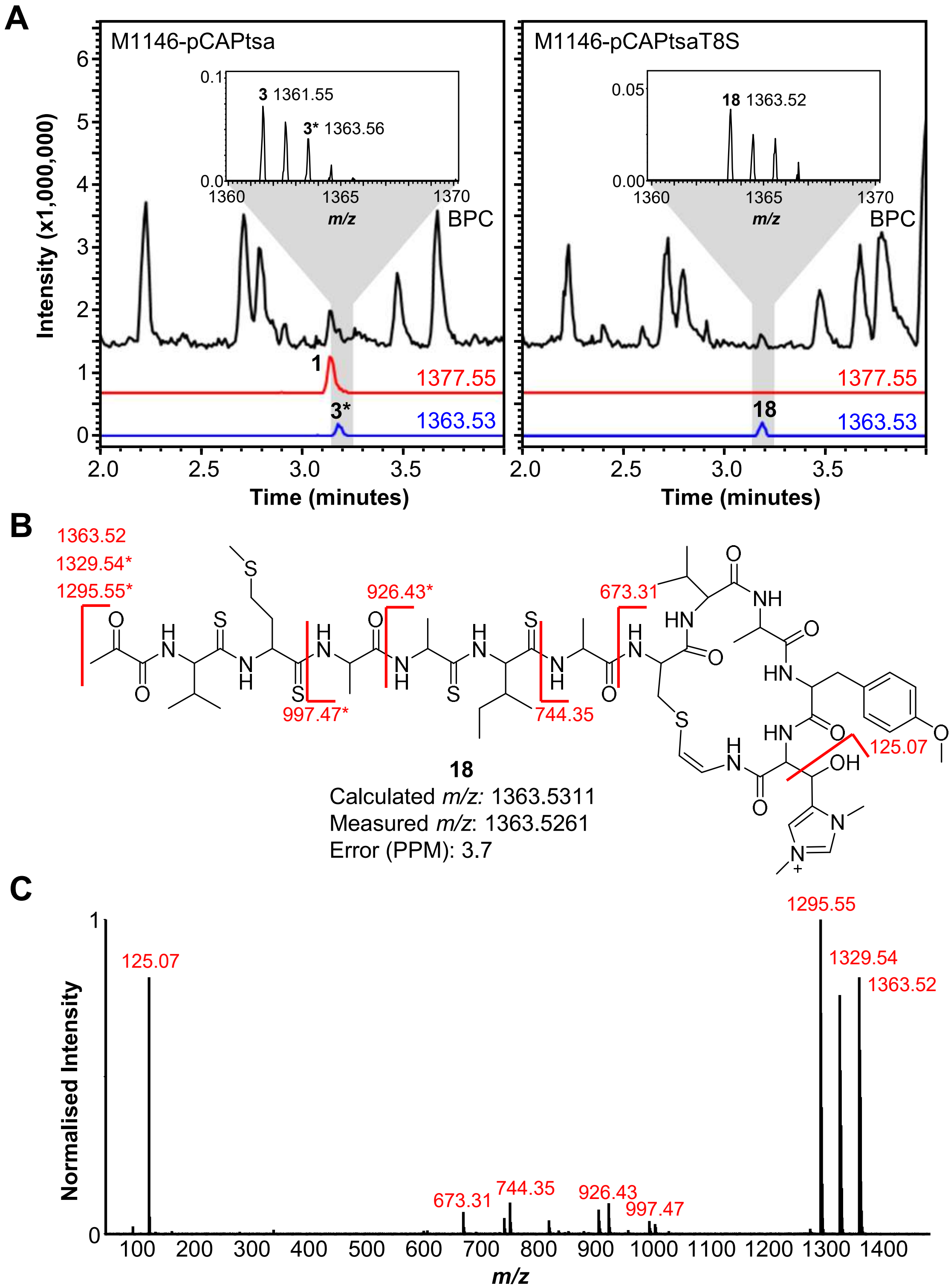


**Figure S20** A. LC-MS analysis of M1146-pCAPtsa and M1146-pCAPtsaT8S. EICs of *m/z* 1377.55 and *m/z* 1363.53 are shown. **3*** labels the second isotope peak of compound **3** (see Figure S5). Mass spectrums are shown to enable the distinguishing between **3** and **18**, as their retention times are the same. B. Predicted structure of **18** annotated with fragments observed via MS/MS analysis. Fragments marked with an asterisk show an additional loss of 33.99, characteristic of the loss of $SH_2$ from thioamide bonds. C. MS/MS spectrum of *m/z* 1363.53 from M1146-pCAPtsaT8S.

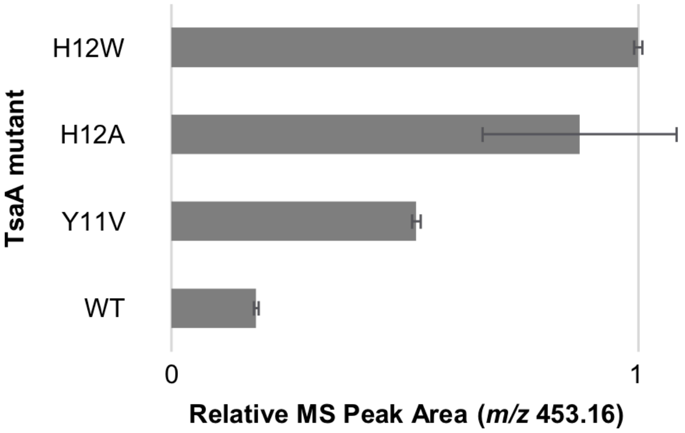


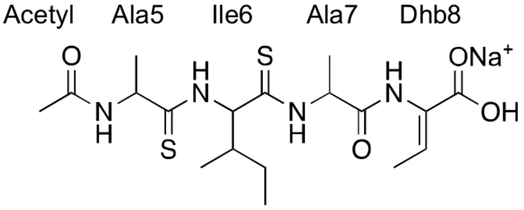


**12**
$[M+Na]^+$
*m/z* 453.16

**Figure S21** MS peak areas of **12** (*m/z* 453.16) in *S. coelicolor* M1146 expressing H12W, H12A, Y11V and wild type (WT) clusters. The bar chart is normalised to the highest mass spectral area (H12W). Values represent the average of three biological replicas and the error bars represent the standard error.

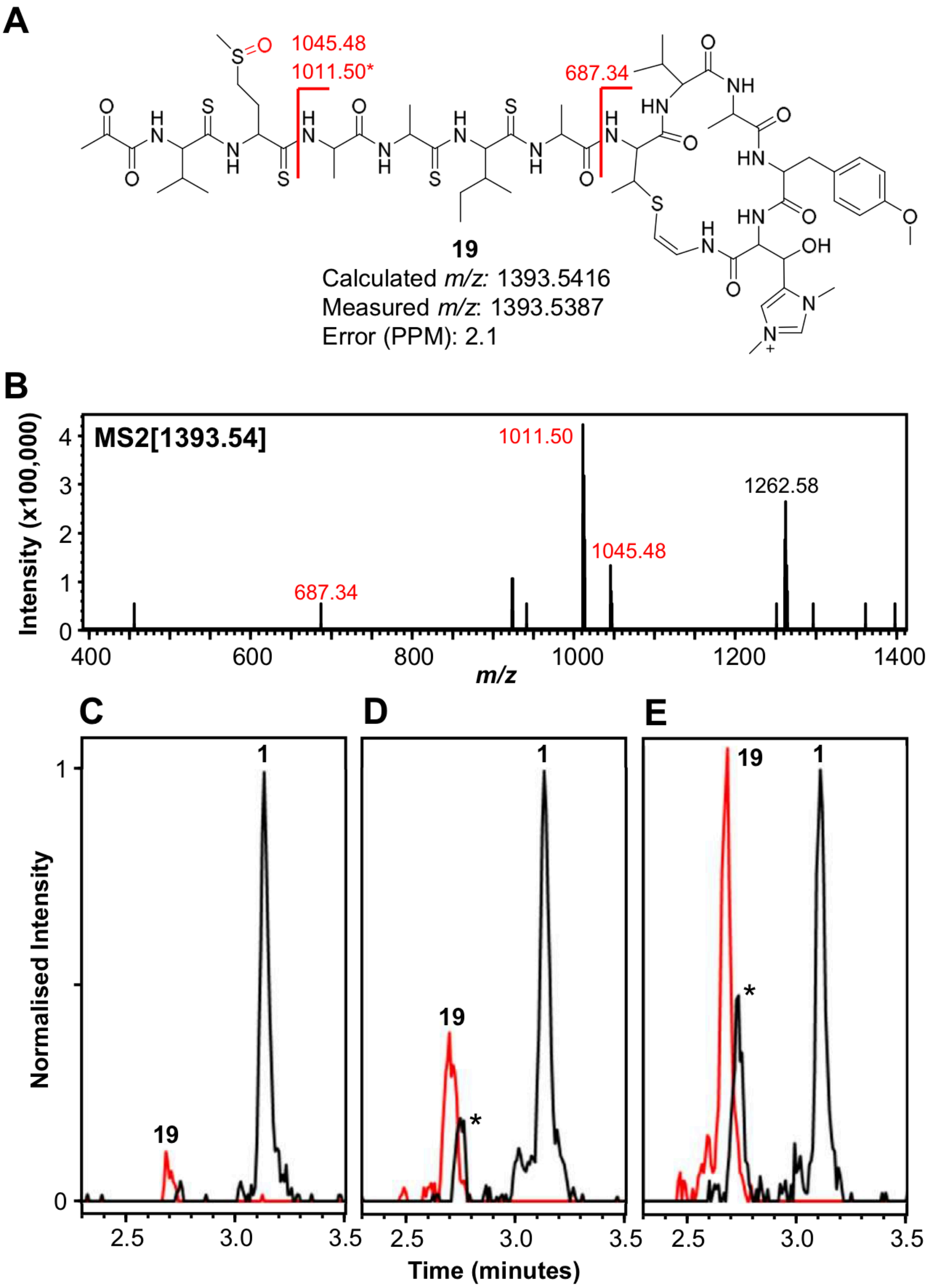


**Figure S22** A. Proposed structure of **19**, the methionine sulphoxide version of **1**. The oxidation is highlighted in red and the fragments seen during fragmentation are marked on the molecule. Fragments marked with an asterisk show a loss of 33.99 (loss of $SH_2$ from thioamide bonds). B. MS/MS spectrum of **19**. Panels C, D and E show EICs of *m/z* 1377.55 (black) and *m/z* 1393.54 (red). Molecules **1** and **19** are labelled, where the peaks labelled with an * are predicted to be a methionine sulphoxide version of **3** (see Figure S5). Each chromatogram is normalised to the intensity of **1**, as a quantitative comparison between these samples was not possible. C. EICs following step one of **1** purification (methanol extraction from M1146-pCAPtsa). D. EICs following step two of **1** purification (liquid-liquid extraction using EtOAc). E. EICs following step three of **1** purification (Sephadex chromatography).

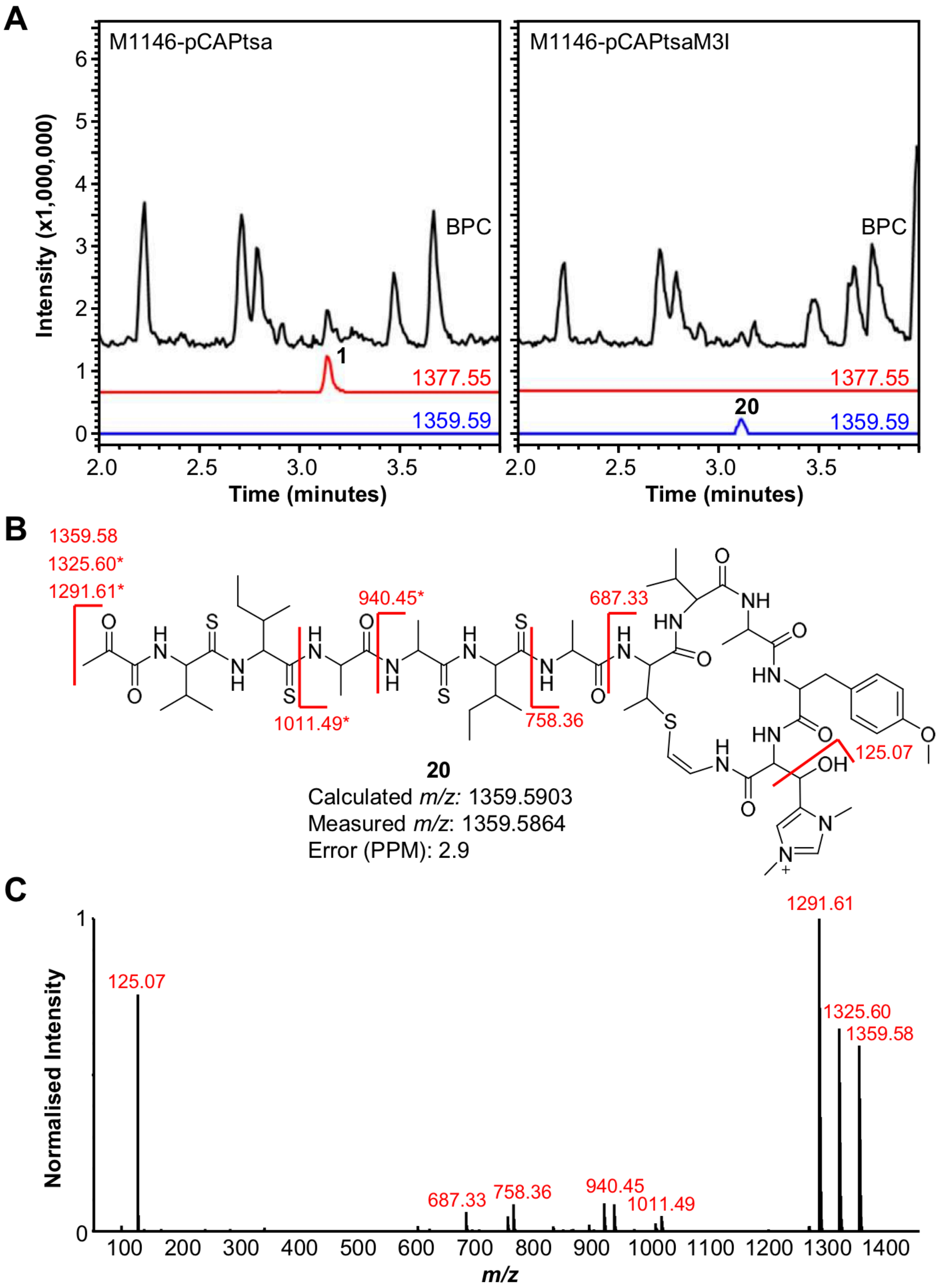


**Figure S23** A. LC-MS analysis of M1146-pCAPtsa and M1146-pCAPtsaM3I. EICs of *m/z* 1377.55 and *m/z* 1359.59 are shown. B. Predicted structure of **20** annotated with fragments observed via MS/MS analysis. Fragments marked with an asterisk show an additional loss of 33.99, characteristic of the loss of $SH_2$ from thioamide bonds. C. MS/MS spectrum of *m/z* 1359.59 from M1146-pCAPtsaM3I.

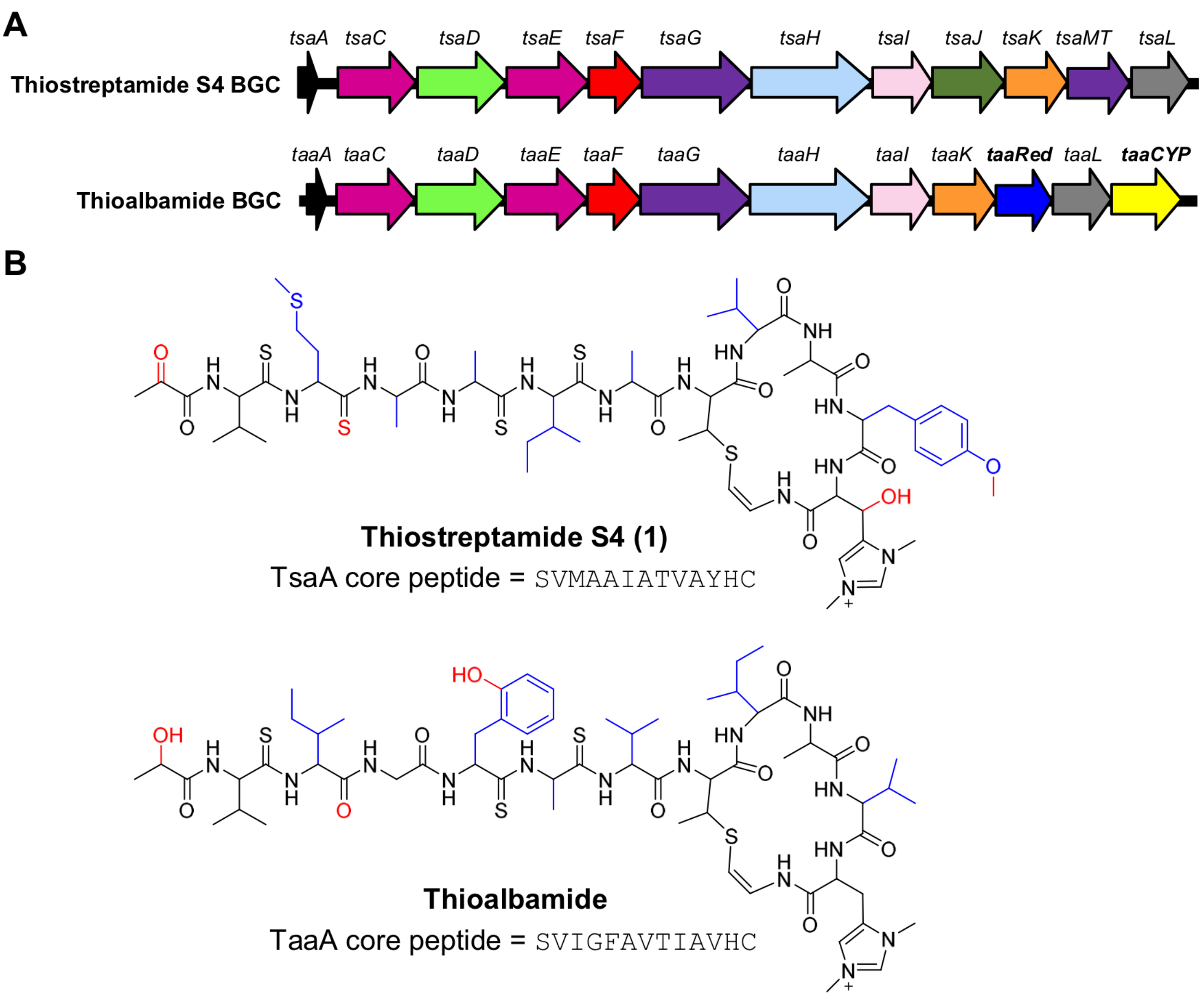


**Figure S24** Comparison between thiostreptamide S4 and thioalbamide[25]. A. BGCs with genes unique to thioalbamide highlighted in bold. B. Structures of thiostreptamide S4 and thioalbamide with structural differences due to amino acid changes are highlighted in blue, while differences due to tailoring enzymes are highlighted in red.

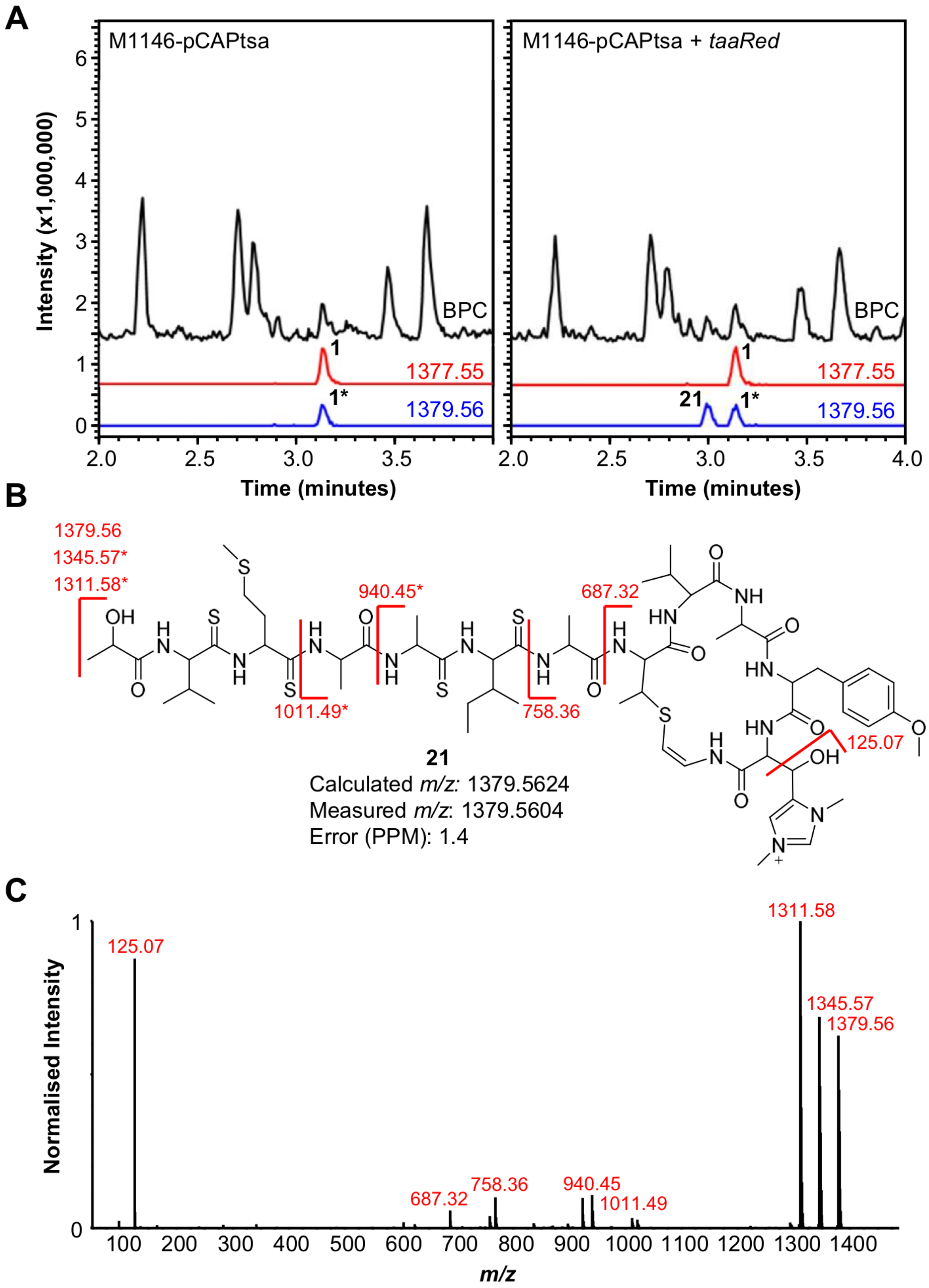


**Figure S25** Activity of TaaRed from thioalbamide pathway expressed alongside pCAPtsa. A. LC-MS analysis of M1146-pCAPtsa and M1146-pCAPtsa + *taaRed*. EICs of *m/z* 1377.55 and *m/z* 1379.56 are shown. **1*** labels the second isotope peak of **1**, indicating that TaaRed does not fully reduce **1** in these production conditions. B. Predicted structure of **21** annotated with fragments observed via MS/MS analysis. Fragments marked with an asterisk show an additional loss of 33.99, characteristic of the loss of $SH_2$ from thioamide bonds. C. MS/MS spectrum of *m/z* 1379.56 from M1146-pCAPtsa + *taaRed*.

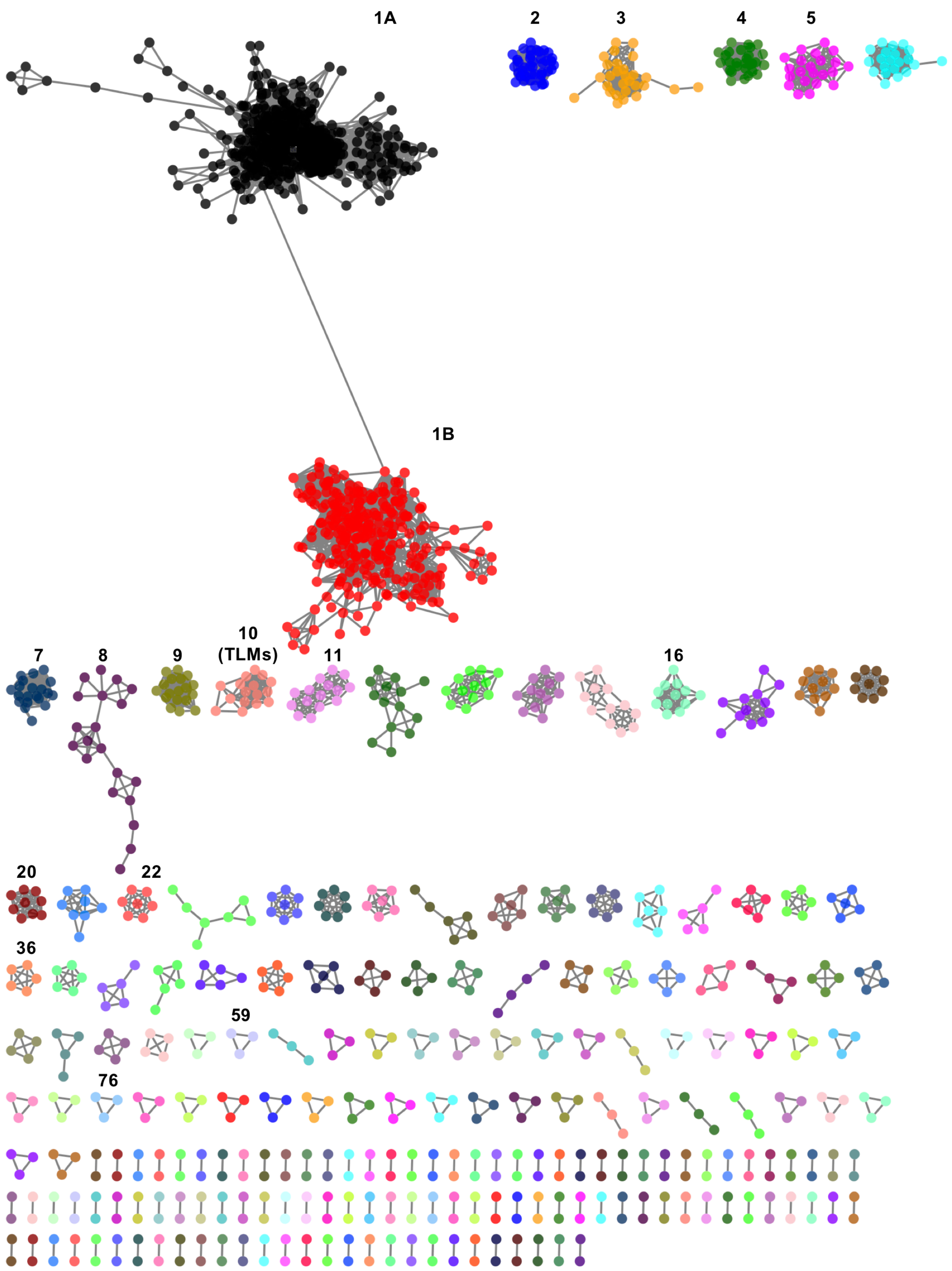


**Figure S26** Short peptide networks associated with HopA1 domain proteins detected by RiPPER. Each of the networks is composed of short peptides sharing a minimum 40% identity. Networks that are shown in Figure 7 of the main paper or in Supplementary Dataset 3 are labelled. Network image generated in Cytoscape[15].

**Network 1A**

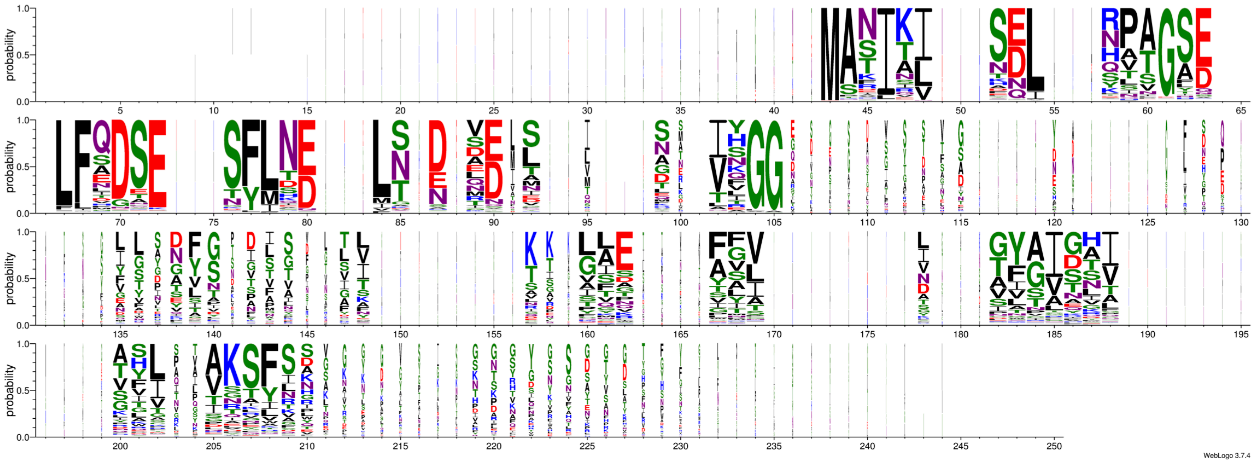

**Network 1B**

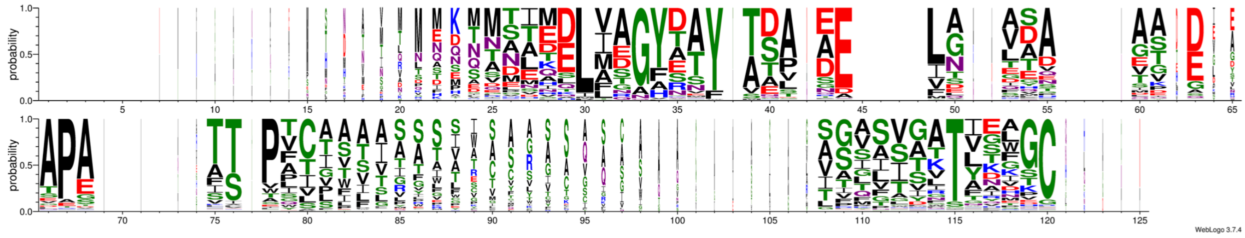

**Figure S27** Sequence logos for all peptides in Networks 1A and 1B showing amino acid composition at each position of aligned sequences. Figure generated using WebLogo 3, where the amino acid colour corresponds to the chemistry (polar = green, neutral = purple, basic = blue, acidic = red, hydrophoblic = black) and the width of each stack is proportional to the fraction of residues in that position (i.e. if a position in an alignment has many gaps and is therefore not conserved it will be visualised as a thin stack).

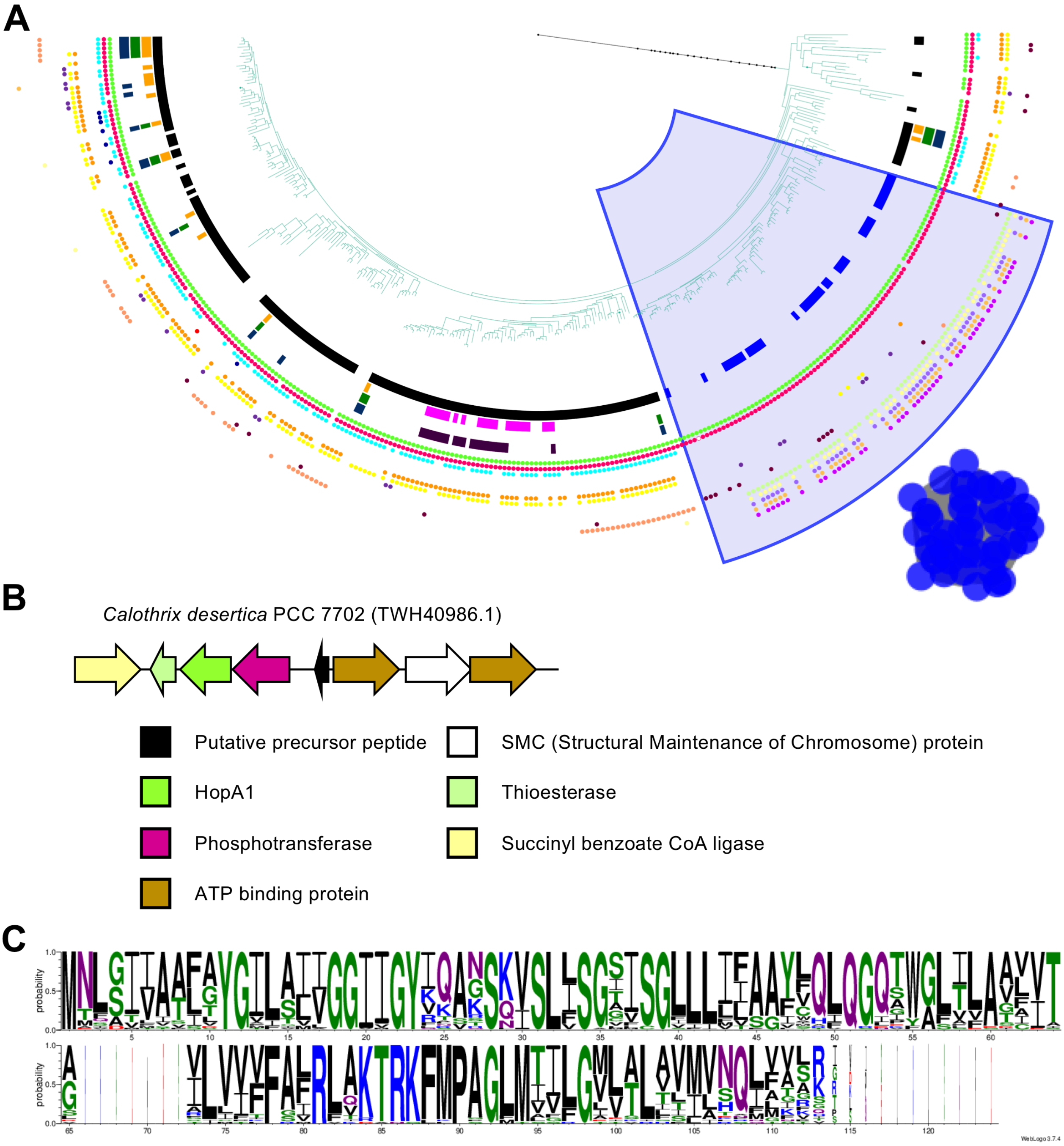


**Figure S28** Analysis of peptide network 2 and its genetic context. A. Partial view of the HopA1 phylogenetic tree showing its association to peptide network 2. The tree branches are colour coded as in Figure 7 of the main text. All branches shown here are cyanobacterial proteins. Conserved pfam domains in the genetic context of each HopA1 proteins are shown as coloured dots in the outer rings of the tree, and network 2 is shown. B. Genetic organisation of a representative gene cluster containing a peptide from network 2. The genes present in the clusters are colour coded to match the pfam annotation on the phylogenetic tree. C. Sequence logo representation of an alignment of precursor peptides belonging to network 2. See Figure S27 for details of logo visualisation.

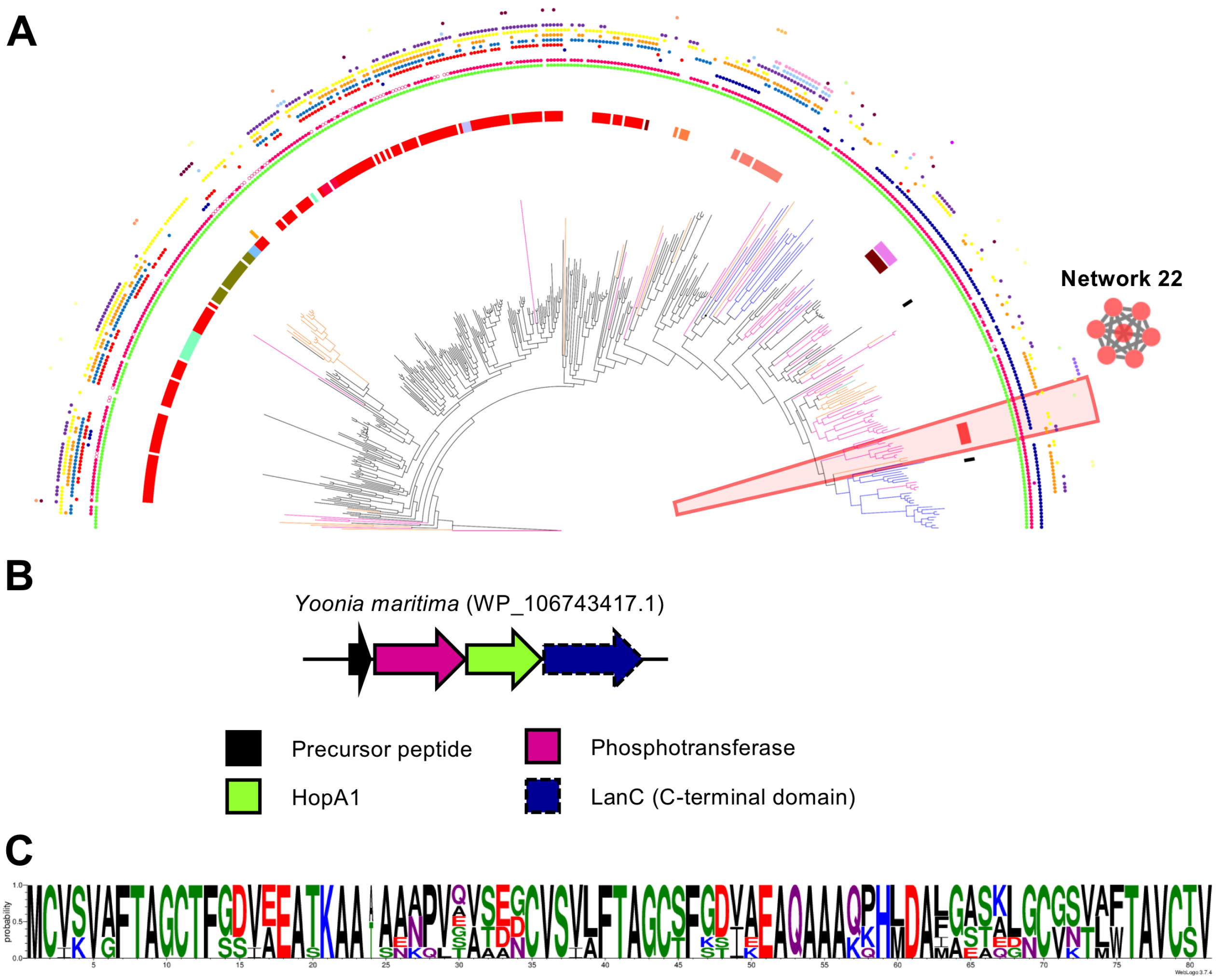


**Figure S29** Analysis of peptide network 22 and its genetic context. A. Partial view of the HopA1 phylogenetic tree showing its association to peptide network 22. The tree branches are colour coded as in Figure 7 of the main text. Conserved pfam domains in the genetic context of each HopA1 proteins are shown as coloured dots in the outer rings of the tree, and network 2 is shown. B. Genetic organisation of a representative gene cluster containing a peptide from network 2. The genes present in the cluster are colour coded to match the pfam annotation on the phylogenetic tree. C. Sequence logo representation of an alignment of precursor peptides belonging to network 22. See Figure S27 for details of logo visualisation.

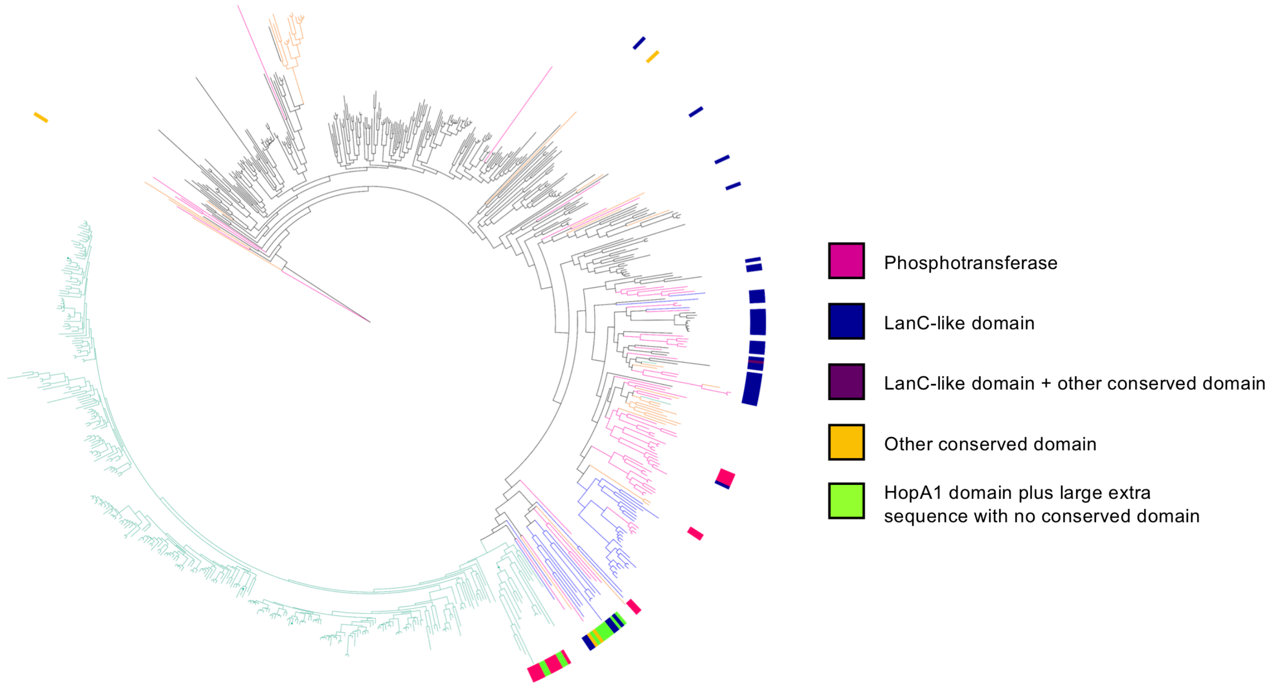


**Figure S30** HopA1 protein fusions. Proteins where the HopA1 domain is physically fused to another functional domain are mapped onto the HopA1 protein phylogenetic tree, with different colour strips for fusions containing each associated domain. Here, all proteins longer than 500 amino acids (AA) are labelled and assessed for additional conserved domains. This identified proteins between 511-976 AA, with an average of 732 AA. The tree branches are colour coded according to the taxonomic origin of the protein as in Figure 7 of the main paper.

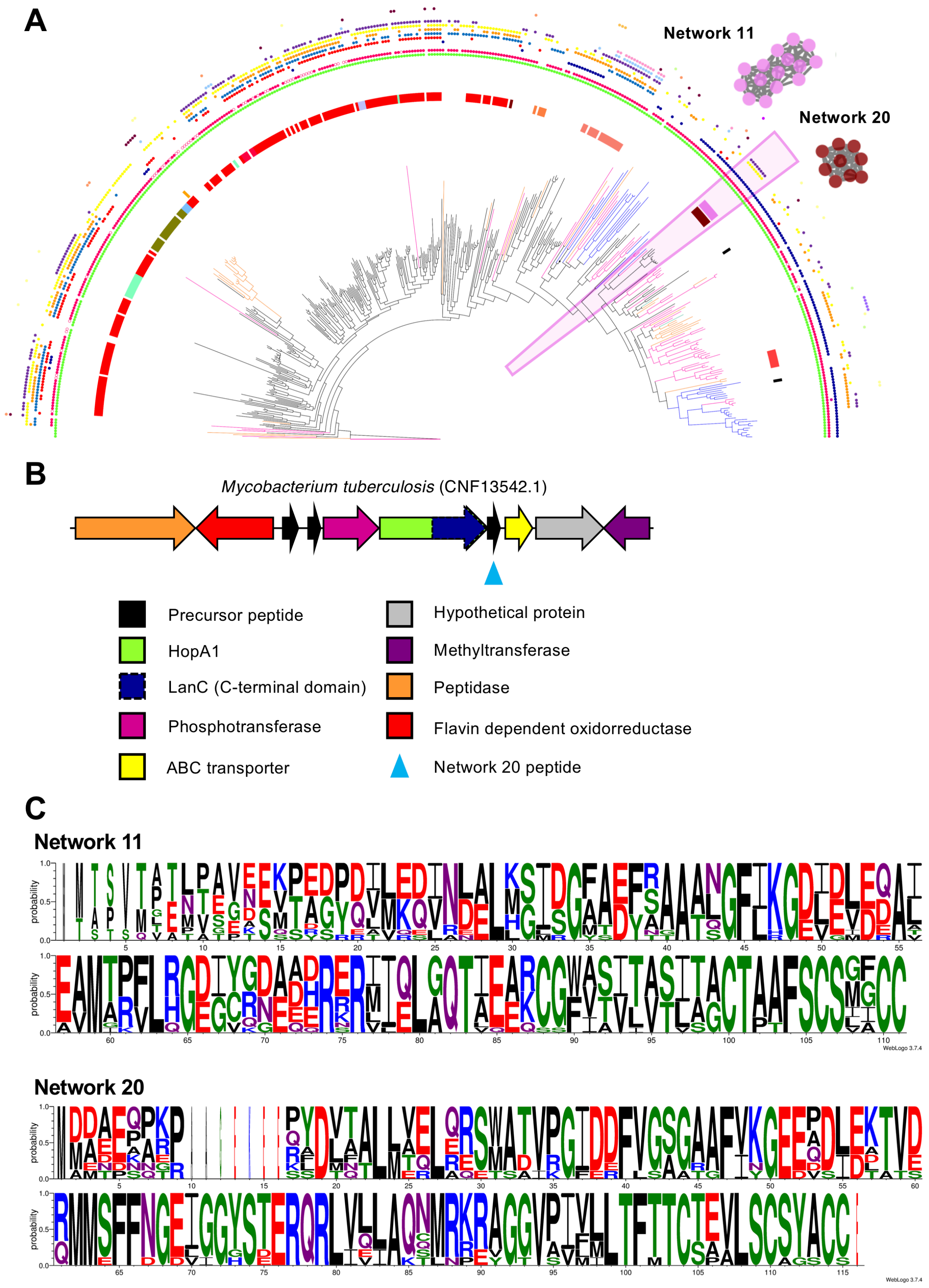


**Figure S31** Analysis of peptide networks 11 and 20 and their genetic context. A. Partial view of the HopA1 phylogenetic tree showing its association to peptide networks 11 and 20 (pink and burgundy bars annotated on tree, as in Figure 7 of the main paper). The tree branches are colour coded as in Figure 7 of the main text. Conserved pfam domains in the genetic context of each HopA1 proteins are shown as coloured dots in the outer rings of the tree, and networks 11 and 20 are shown. B. Genetic organisation of a representative gene cluster containing peptides from each network. The genes present in the clusters are colour coded to match the pfam annotation on the phylogenetic tree. C. Sequence logo representation of an alignment of precursor peptides belonging to networks 11 and 20. See Figure S27 for details of logo visualisation.

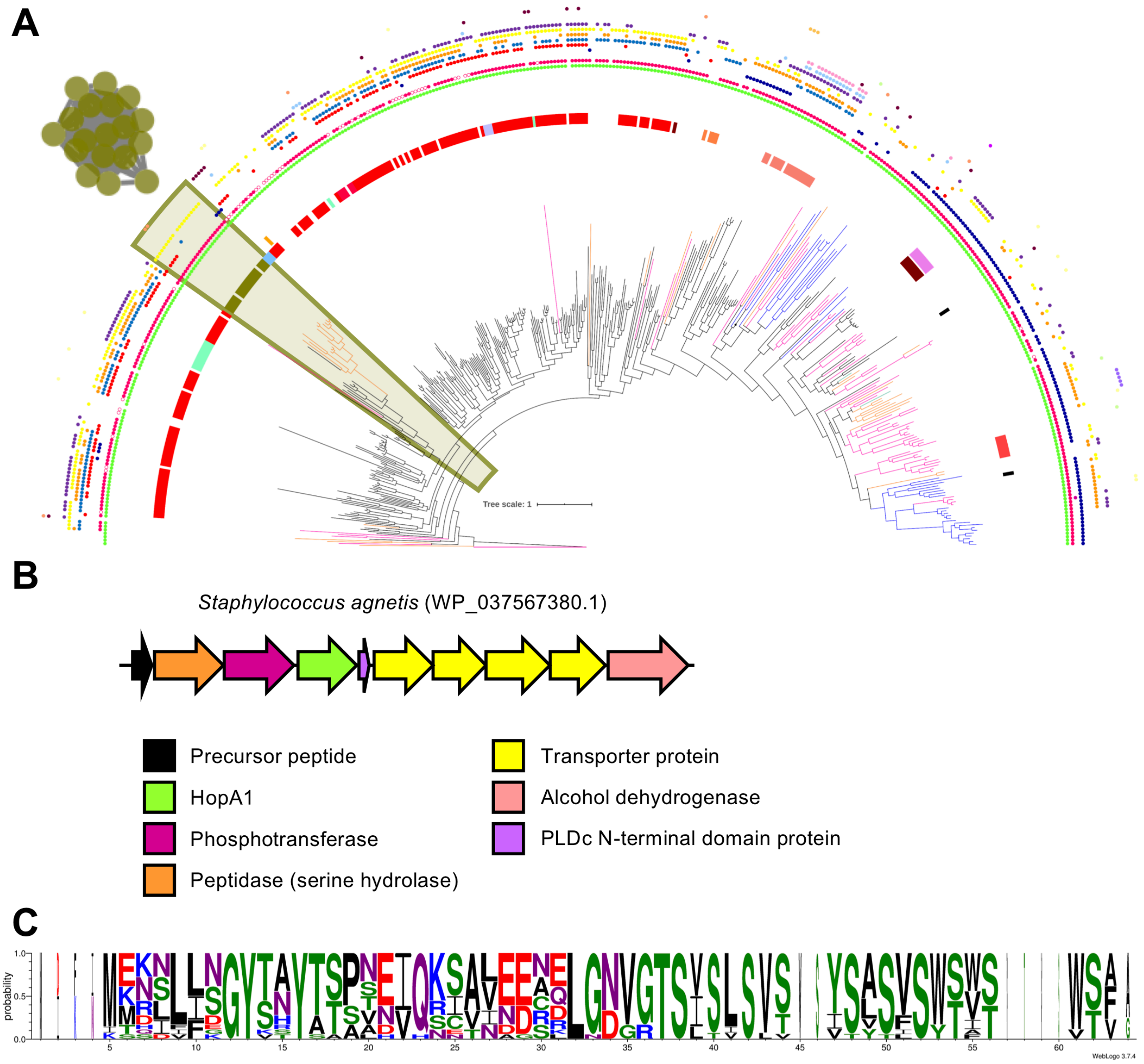


**Figure S32** Analysis of peptide network 9 and its genetic context. A. Partial view of the HopA1 phylogenetic tree showing its association to peptide network 9. The tree branches are colour coded as in Figure 7 of the main text. Conserved pfam domains in the genetic context of each HopA1 proteins are shown as coloured dots in the outer rings of the tree, and network 2 is shown. B. Genetic organisation of a representative gene cluster containing a peptide from network 2. The genes present in the cluster are colour coded to match the pfam annotation on the phylogenetic tree. C. Sequence logo representation of an alignment of precursor peptides belonging to network 22. See Figure S27 for details of logo visualisation. PLD = phospholipase D nuclease.

### Network 3

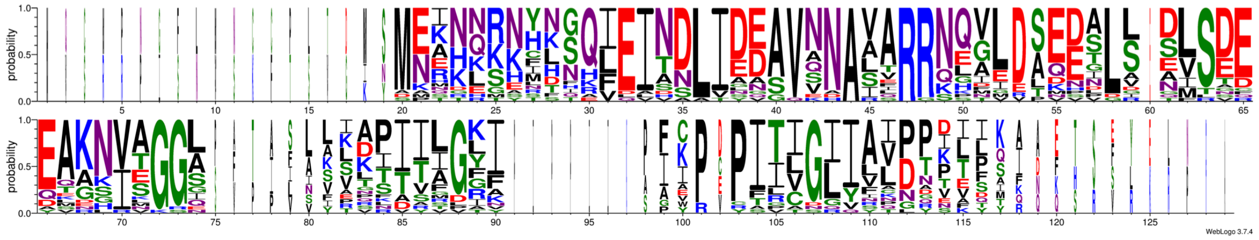

### Network 4 (Nif11 domain peptides)

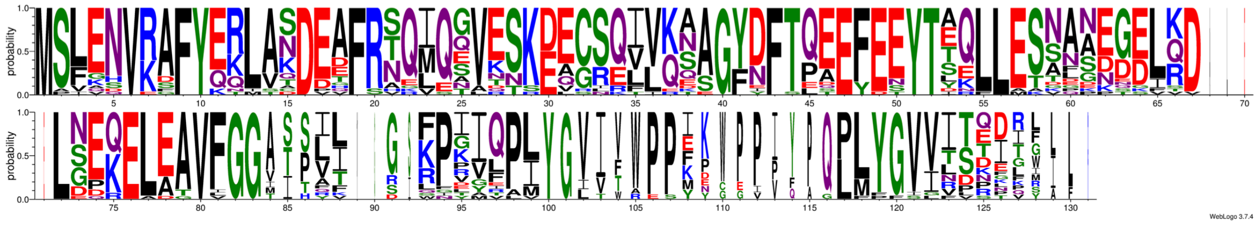

### Network 5

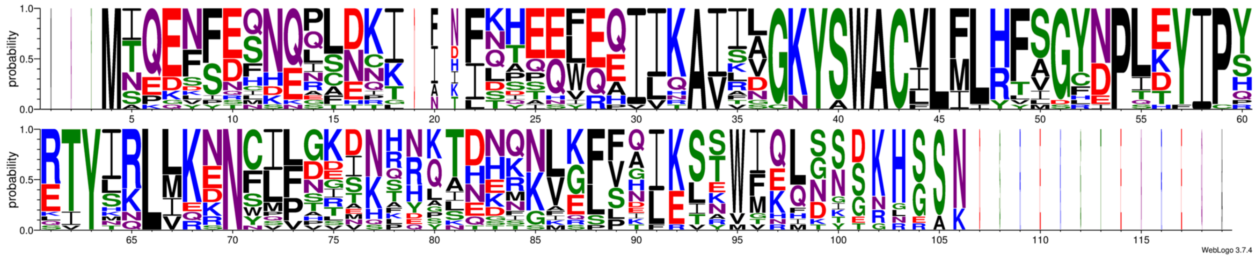

### Network 7

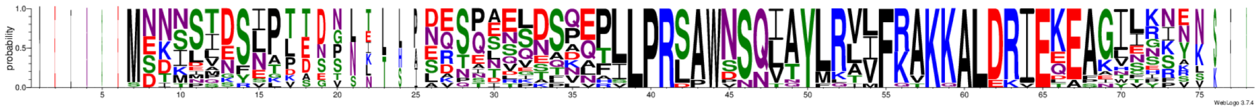

### Network 8

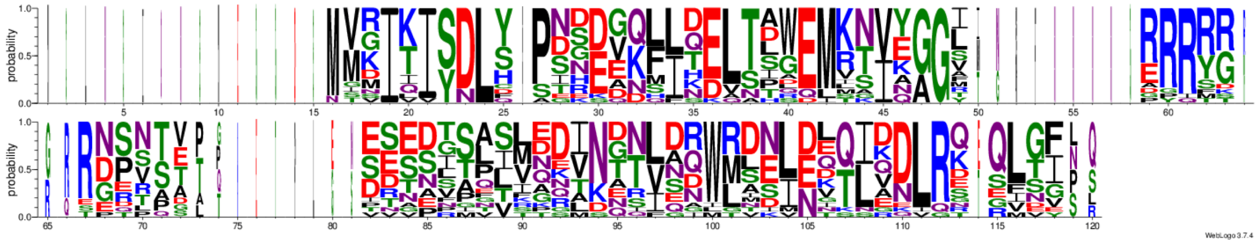

**Figure S33** Logo representation of the precursor peptides belonging to networks 3, 4, 5, 7 and 8. These peptides appear in close proximity with peptides belonging to other networks, as part of the same biosynthetic gene clusters (see Figure 8 of main paper). See Figure S27 for details of logo visualisation.
