## Supplementary figures and images for "Understanding Thioamitide Biosynthesis Using Pathway Engineering and Untargeted Metabolomics"

### Supplementary Dataset 3

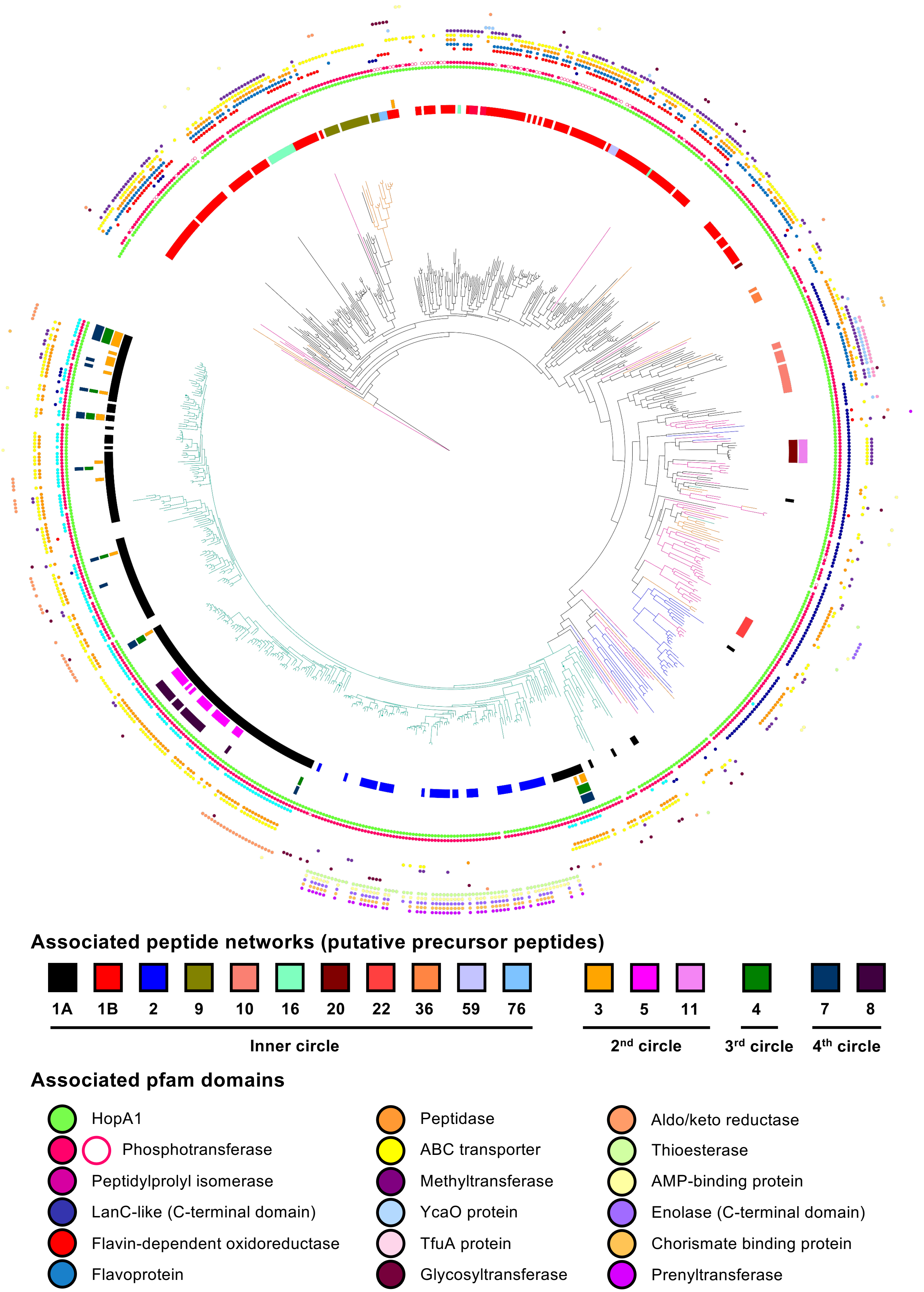
